# Bradykinesia and postural instability in a model of prodromal Synucleinopathy with α-Synuclein aggregation in the gigantocellular nuclei

**DOI:** 10.1101/2024.09.05.610956

**Authors:** Vasileios Theologidis, Sara A. Ferreira, Nanna M. Jensen, Diana Gomes Moreira, Ole A. Ahlgreen, Mads W. Hansen, Emilie D. Rosenberg, Mette Richner, Islam Faress, Hjalte Gram, Poul H. Jensen, Per Borghammer, Jens R. Nyengaard, Marina Romero-Ramos, Christian B. Vægter, Wilma D. J. van de Berg, Nathalie Van Den Berge, Asad Jan

**Affiliations:** Department of Clinical Medicine, Aarhus University, Palle Juul-Jensens Boulevard 35, DK-8200 Aarhus; Department of Biomedicine, Aarhus University, Høegh-Guldbergs Gade 10, DK-8000 Aarhus Denmark; Danish Research Institute of Translational Neuroscience (DANDRITE), Department of Biomedicine, Aarhus University, Ole Worms Allé 3, DK-8000 Aarhus C, Denmark; Core Center for Molecular Morphology, Section for Stereology and Microscopy, Department of Clinical Medicine, Aarhus University, Palle Juul-Jensens Boulevard 35, DK-8200 Aarhus; Department of Anatomy and Neurosciences, Amsterdam Neuroscience, VU University Medical Center Amsterdam, De Boelelaan 1117, 1081 HV Amsterdam, The Netherlands

**Keywords:** Parkinson Disease, Alpha-Synuclein, Lewy pathology, Gigantocellular nucleus

## Abstract

α-Synuclein (aSyn) accumulation within the extra-nigral neuronal populations in brainstem, including the gigantocellular nuclei (GRN/Gi) of reticular formation, is a recognized feature during the prodromal phase of Parkinson disease (PD). Accordingly, there is a burgeoning interest in animal model development for understanding the pathological significance of extra-nigral synucleinopathy, in relation to motor and/or non-motor symptomatology in PD. Here, we report an experimental paradigm for the induction of aSyn aggregation in brainstem, with stereotaxic delivery of pre-formed fibrillar (PFF) aSyn in the pontine GRN of transgenic mice expressing the mutant human Ala53Thr aSyn (M83 line). Our data show that PFF aSyn-induced aggregate pathology in GRN leads to progressive decline in spontaneous locomotion and an early phenotype of postural instability. This early phase of bradykinesia was followed by a moribund stage, characterized by worsening motor performance and impaired survival with substantial aSyn aggregation in several brain regions beyond the GRN. Collectively, our observations suggest an experimental framework for studying the pathological significance of aSyn aggregation in GRN in relation to features of movement disability in PD. With further refinements, we anticipate that this model holds promise as a test-bed for translational research in PD and related disorders.

## INTRODUCTION

Idiopathic Parkinson disease (PD) is the most common cause of movement disability, clinically defined by the acronym TRAP: resting Tremor, Rigidity, difficulty in movement initiation (bradykinesia/Akinesia), and Postural instability (1, 2). The prevalent notion concerning the movement disability in PD implicates the progressive decline of dopaminergic neurotransmission in the nigro-striatal circuitry, arising due to the loss of dopaminergic neurons in the midbrain substantia nigra-*pars compacta* (SN*pc*) (1, 2). In addition, a substantial number of PD patients report non-motor symptoms (olfaction, autonomic, sleep and pain-related), which significantly impair the quality of life (3). In a larger context, it is increasingly being recognized that the nigro-centric view is not sufficient to account for the heterogeneity in clinical presentation of PD and related disorders (2–4). The nature of pathological process(es) which trigger neuronal dysfunction and/or neurodegeneration in SN*pc* remains an active subject of investigation. In this regard, aggregation of α-Synuclein (aSyn; gene symbol *SNCA*) in SN*pc* and several extra-nigral regions (ie. outside SN*pc*) is considered to be a potent aggravating factor in the pathogenesis of PD and related synucleinopathies (5, 6).

According to the Braak staging scheme, neuronal populations within the dorsal motor nucleus of vagus (*dmX*), locus coeruleus (LC) and the nuclei of reticular formation including the gigantocellular nuclei (GRN/Gi) bear the brunt of cellular aSyn pathology during the early stages of PD (7–11). These observations ushered novel efforts in the development of refined animal models for studying the neurological basis of PD symptomatology, which are not confined by the prevalent nigro-centric view. The utility of these efforts is highlighted by the studies showing that extra-nigral brainstem synuclienopathy (eg. in *dmX* and/or LC) recapitulates PD-like non-motor symptoms and autonomic dysfunction in rodents (12–15). Intriguingly, the significance of cellular aSyn pathology affecting GRN and nearby nuclei of brainstem reticular formation in the context of PD symptomatology remains largely unexplored.

The GRN is a prominent collection of neurons within the paramedian parts of pontomedullary reticular formation (16). It has been suggested that neuronal populations of GRN, in concert with basal ganglia, are involved in smooth execution of complex movements, including turning, gait stance and stopping the locomotion in freely moving animals (17–19). Neuroanatomically, the GRN is one of the major sources of input into the reticulospinal tracts, which converge on motor and premotor neurons at all levels of spinal cord, and modulates the excitability of spinal motor system (9, 20, 21). Moreover, in concert with raphe magnus and periaqueductal grey (PAG), the neurons in GRN plausibly mediate the modulation of pain perception, through descending projections onto the spinal nociceptors in the dorsal horn (16, 22–24). In addition, these nuclei also receive substantial input from cerebellum and spinal cord, and are integral components in the coordination of reflex motor activity in the maintenance of posture and balance (9, 16, 25).

Therefore, we hypothesized that direct induction of aSyn aggregation within GRN of rodents will lead to the emergence of unique sensorimotor phenotypes, which could potentially be relevant to PD symptomatology. In particular, we wanted to study the patterns of locomotion, movement coordination and nociception in relation to the emergence and propagation of aSyn pathology in brainstem, with GRN as the initial nidus of aSyn aggregation. In transgenic mice expressing the human mutant Ala53Thr aSyn ((M83 line, (26)), we observed that *de novo* induction of aSyn aggregation in GRN, -by stereotaxic delivery of pre-formed fibrillar (PFF) aSyn-, was associated with progressive reduction in spontaneous locomotion and subtle defects in postural motor coordination, long before phenotypes reflecting motor weakness were seen. With the progression of aSyn pathology into additional nuclei in the brainstem, the animals exhibited worsening deficits in movement coordination and defective response in tests of nociception, with consequent decline in survival. With this context, we highlight the implications of our findings *vis-a-vis* further refinements in model development for PD and related disorders, and also discuss the limitations of the model.

## MATERIALS AND METHODS

### Generation and characterization of mouse aSyn fibrils

Mouse aSyn fibrils were prepared and characterized *in vitro*, essentially as described (27, 28). Briefly, full length recombinant (wild type) mouse aSyn was expressed in BL21(DE3) competent cells and purified using ion-exchange and reverse phase chromatography. The purified aSyn was re-suspended in phosphate-buffered saline (PBS, pH 7.4) at 1 mg/mL and passed through a 100kDa filter. The monomeric (non-aggregated) aSyn was then incubated with sonicated mouse pre-formed fibrillar (PFF; 5% by mass in PBS) in a seeded aggregation assay (28). The sample was incubated at 37°C with continuous shaking at 1050 r.p.m. in a tabletop microtubes shaker (Eppendorf) for 72h. The insoluble PFF were collected by centrifugation (15,600 g at 25°C for 30 min) and then re-suspended in PBS. Protein concentration was determined by the BCA assay (Pierce) and a stock solution consisting of 5 mg/mL protein was prepared (in PBS). Subsequently, PFF were sonicated for 20 min using a Branson 250 Sonifier at 30% intensity, and then aliquoted and frozen at□−□80°C until further use.

### Animal Studies

#### Animal care and husbandry

Transgenic M83 mice [B6;C3-Tg(Prnp-SNCA*A53T)83Vle/J]- (26) were housed at the Skou animal facility at Aarhus University in accordance with Danish regulations and the European Communities Council Directive for laboratory animals (license # 2022-15-0201-01294 issued to CBV, co-author). The animals were housed under a 12 hours light/dark cycle and fed with regular chow diet *ad libitum*. For the study, adult mice (12-14 weeks of age) were used, and the cohorts included both male and female animals.

#### Intracerebral aSyn injection in the pontine GRN

PFF aSyn, monomeric aSyn or PBS were bilaterally delivered into the pontine GRN under isoflurane anesthesia (2-5%), using the stereotaxic coordinates: [AP (y): −□6; ML (x): 0.5 relative to bregma; DV (z)- 2 locations:−□5 mm (1 µl) and -5.2 mm (1 µl) relative to dura]. Hence, 2 µl of aSyn preparations (PFF or monomeric, amounting to 10 µg total protein) were injected bilaterally, at a flow rate of 0.2 µl/min through a 5 μl Hamilton syringe (33-gauge needle), connected to a stereotaxic frame. The needle was left in place (at DV -5.2 mm) for an additional 1 min, and then gently withdrawn over 15 sec. The main study consisted of the following cohorts of heterozygous M83^+/-^ mice: i) PFF aSyn (n=18), ii) monomeric (non-aggregated) aSyn (n=5) and iii) PBS (n=5). As a proof-of-concept, a small pilot study involving stereotaxic delivery of PFF aSyn in GRN of homozygous M83^+/-+^ mice (n=4) was also performed using identical experimental setup. After the surgical procedure, the animals were allowed to recover in their home cage (placed on a heated blanket), and received appropriate analgesia based on the veterinarian’s recommendations.

### Behavioral Assessments

After recovery (14 days), the animals were tested in a battery of sensorimotor tasks (described below) periodically over a period of 120 days post-injection (DPI). Prior to the tests, the animals were acclimatized to the testing environment with standard lighting conditions and ambient background noise for 1 hour, unless indicated otherwise.

### General Locomotion

#### Non-invasive monitoring of spontaneous activity in home cage

Patterns of locomotion and spontaneous activity were monitored in home cage through specialized digitally ventilated cages (DVC) platform (Tecniplast, Italy). This platform is based on the electrical capacitance sensing technology, with incorporates a sensor board equipped with an integrated circuit comprised of 12 electrodes directly beneath the floor of home cage (29). The DVC circuit measures changes in the electrical capacitance signal from each electrode in response to the movement of a water-filled body (animal) close to or away from a given electrode. The measurements, performed approximately 4 times per second, are remotely relayed to the centralized DVC analytics platform (Tecniplast, Italy). In this web-based interface, time-stamped data for each cage can be visualized using in-built tools (e.g. daily rhythms, cumulative activity/locomotion index aggregated per minute/hour, bedding status, light or dark period activity, heatmaps etc.). In the default setup, the DVC analytics web-interface plots the animal locomotion index as arbitrary units normalized between 0% and 100%, representing the overall activity performed in the cage by the animals, ie. the signal is measured for each cage and not each animal, unless the animals are singly housed. The detailed description of DVC working principle and DVC analytics platform is included in the Supplementary Information.

#### Open-Field Test

General locomotor activity and exploratory behavior were assessed in an open-field chamber (40 cm × 40 cm × 30 cm) with video recordings over a period of 10 min obtained through an overhead USB camera, operated by the ANY-maze analytical software (Vendor: Stoelting Europe). The test parameters included distance traveled (m), mean speed (m/s), time freezing (s), and number of entries into defined zones (center, periphery and intermediate).

### Balance, Movement coordination and Motor strength

#### Balance beam Test

Fine motor coordination and balance were assessed by the balance beam test with slight modifications (30). The test apparatus consisted of flat surface metal beams (length: 1 meter; width: 8 cm or 16 cm), supported by two poles (height: 60 cm), and equipped with a nylon hammock underneath (10 cm above the ground). A source of bright light (lamp) was used as an aversive stimulus on the starting end. The mice were trained to traverse the 2 beams (3 attempts on 2 consecutive days; each attempt separated by 15 min), towards a clean cage with some bedding from the home cage. During the test, video recordings of the behavior were obtained through a USB camera (operated by Microsoft Windows), placed within 5 cm from the starting end and at the same height as the beam. Each animal was tested 2 times on each beam with inter-trial duration of 15-20 min, and the mean of the measurements was calculated. Video recordings in slow motion were analyzed for the traversal time (s), number of hindpaw slips and mean speed of traversal (m/s).

#### Pole Test

Fine motor coordination was assessed in a pole test (31), with a test apparatus consisting of a wooden pole (height: 50 cm, diameter: 1 cm), which was supported by a circular base stand (diameter: 10 cm) placed in a clean cage containing some bedding from the home cage). The mice were gently placed within 2 cm of the top of the pole facing up and away from the testing personnel. Video recordings of the behavior were obtained through a USB camera (operated by Microsoft Windows), placed within 25 cm from the pole such that the whole length of pole could be recorded (from a side view, approx. 90°). Each animal was trained over 2 consecutive days with 5 trials, with inter-trial duration of 15 min. During the test, each animal was subjected to 3 trials, with inter-trial duration of 15-20 min, and average of the measurements was calculated. Video recordings in slow motion were analyzed for assessing the turning time (t1), traversal time after turning and reaching the base stand (t2) and total time on the pole (t1+t2). If the animal paused while descending, the trial was repeated. If the animal fell off during the descent, a maximum score of 25 sec was assigned to the traversal time (t2) and 30 sec to the total time (t1+t2).

#### Rotarod Test

Fore- and hindlimb motor coordination and balance were assessed by placing the mice on an accelerating rod (Rotarod; LE8500 Harvard Apparatus)- (32). The mice were trained to walk in a forward direction on the rotarod for 60-90 sec with a fixed rotation rate of 4 rpm (3 attempts on 3 consecutive days; each attempt separated by 10 min). If the animal fell off before 60 s, it was returned to home cage and the training attempt was resumed after approx. 15 min. During the test, the animal was placed on the rotarod and acceleration from 4 to 40 rpm in 120 sec was initiated. Each animal was tested 3 times, with inter-trial duration of 15-20 min, and average of the measurements was calculated. The analyses included time (sec, latency) to fall and speed at fall (rpm).

#### Grip strength Test

Limb motor strength was assessed by a grip strength test apparatus (BIOSEB, BIO-GS4), according to the manufacturer’s instructions. During the test, the mice were held by the tail and lowered onto a horizontal metal grid connected to a sensor detecting peak tension. After the animal grabbed the metal grid, it was pulled backwards by the tail in horizontal plane with a gentle constant pressure. Average maximal peak force (grams) exerted by the paws was recorded from 3 consecutive trials, with an inter-trial interval of approx. 10 sec.

#### Footprint Test

Gait stance was assessed by a footprint test with slight modifications (32). The apparatus consisted of a flat wooden platform (length: 50 cm, width: 5 cm) supported on each end by two wooden poles attached underneath (height: approx. 10 cm). The forepaws and the hindpaws were coated with red and black nontoxic paint, respectively. The mice were placed on a cut sheet of white paper (length: 40 cm, width: 5 cm) lightly affixed to the platform, and trained to traverse the platform to a clean cage with some bedding material from home cage. The training consisted of 3 consecutive attempts on 2 consecutive days (each attempt separated by 2-3 min) one day prior to the test (except, the pre-terminal stage; see Results). A fresh sheet of paper was placed for each mouse during each training session.

On the testing day, the mice were allowed to traverse the platform with a fresh sheet of paper and the paint was air-dried for 30 min. Then, the footprint patterns were analyzed for measures of gait stance from 2-3 consecutive steps made in the forward direction, excluding the footprints made at the beginning and end of the platform. The measures (all in cm) include: (1) Stride length: average distance of forward movement between each stride. (2) Step width: average diagonal distance between alternating front and rear paws. (3) Rear base of support (RBOS) and (4) Frontal base of support (FBOS): average distance between left and right footprints (rear or front paws, respectively), represented by a perpendicular line connecting the center of a given footstep to its opposite preceding and proceeding steps. (5) Step alternation: Overlap between left or right footprints in consecutive steps, ie. distance between the center of the hind footprint and the center of the preceding front footprint. For the gait stance parameters, the mean value of each set comprising 2-3 values in each measure was used in the subsequent analyses.

#### Hindlimb Clasping Test

Assessment of hindlimb clasping was performed by a modified tail suspension test (33, 34). Freely moving, non-anesthetized, mice were held by the tail and lifted in air for 10 sec. Severity of clasping was scored on a scale of 0-3, as follows: a) *Score 0, No clasping* (both hindlimbs were consistently splayed outwards, and away from the abdomen for more than 50% of the time suspended); b) *Score 1, Mild clasping* (one hindlimb was retracted toward the abdomen for more than 50% of the time suspended); c) *Score 2, Moderate clasping* (both hindlimbs were partially retracted toward the abdomen for more than 50% of the time suspended); and d) *Score 3, Severe clasping* (both hindlimbs were entirely retracted, and touching the abdomen for more than 50% of the time suspended).

### Nociception

#### Hot plate

Thermal nociception/allodynia was assessed using a hot plate apparatus (VWR), preheated to a stable temperature of 55°C ± 0.5 (35). For the test, a bottomless plexiglass chamber (15 cm x 15 cm x 15 cm) was placed on a flat metal surface and the animals were individually lowered into the chamber. The latency to response (sec) was recorded manually, when the animal licked the paws or jumped. A cut-off maximum time (30 sec) was used and animals were immediately removed after the response in the test.

#### Von Frey

Mechanical allodynia was assessed by manually applying the calibrated Semmes– Weinstein monofilaments (Stoelting) of ascending force (0.16-2.00 g) onto the plantar surface of the hindpaws, avoiding food pads (36). During the test, the animals were acclimatized in a plexiglas container placed over a mesh metal grid for approximately 10 min prior to testing in an ambient lit room and quiet surroundings. A positive response to given filament application was characterized by sudden paw withdrawal, paw licking or jerky flaying of toes. Each filament was applied 5 times for max. duration of 5 sec to the test subject. The threshold response (g) to a given filament was recorded at a positive response to at least three out of 5 applications of the same filament.

### Histological Analyses

#### Human studies

##### Study cohort, tissue selection and pathological assessment

Post-mortem brain tissue from subjects with clinical parkinsonism and neurologically normal controls were acquired from the Netherlands Brain Bank (NBB; Amsterdam, The Netherlands, http://brainbank.nl). Donors or their next of kin signed informed consent for brain autopsy, the use of brain tissue and the use of medical records for research purposes. The brain donor program of the NBB and NABCA is approved by the local medical ethics committee of the VUmc, Amsterdam (NBB 2009.148). Demographic features and clinical symptoms were retrieved from the clinical files, including sex, age at symptom onset, age at death, disease duration, presence of dementia and parkinsonism. Braak stages for aSyn pathology were determined using the BrainNet Europe (BNE) criteria. Braak neurofibrillary stages were determined according to on NIA-AA consensus criteria. A summary of the clinical and pathological characteristics for all cases can be found in the supplementary file (Table S1).

##### Human tissue processing and Immunohistochemistry (IHC) analyses

IHC detection of p-aSyn (S129) was performed on 6 µm thick formalin-fixed paraffin-embedded (FFPE) tissue sections of medulla oblongata, following deparaffinization and blocking of endogenous peroxidise, according to previously established methods (37). Nonspecific binding was blocked by incubating the sections in in Tris-Buffered Saline (TBS) containing 3% normal donkey serum for 30 min at room temperature (RT). Then, the sections were incubated (overnight, at 4°C) with the primary antibody for detecting p-S129 aSyn (rabbit mAb EP1536Y Abcam, #ab51253-1:4000). Then, the sections were stained with secondary detection solution Envision anti-rabbit (DAKO cat# K4003) for 30 min at RT. Color was developed using the DAB (3,3’-diaminobenzidine) chromogen for 10 min at RT. Nuclear counterstaining was performed in hematoxylin for 20 sec, after which sections were washed under running tap water for 5 min. Sequential dehydration in ethanol was performed in series: 1 x 2 min 70%, 1 x 2 min 80%, 2 x 2 min 96%, 2 x 2 min 100%, followed by 3 x 2 min xylene. Entellan (Merck, cat# 107960) was used as mounting medium when cover-slipping. After cover-slipping, sections were left to dry overnight in the fume hood. Whole slide digital scans of the immunostained sections were obtained using the brightfield mode in Olympus VS200 upright microscope at UMC Amsterdam (20X magnification). Slide scans from PD and control cases were imported in Qupath (v. 0.5.1) and p-aSyn (S129) immunopositivity was computed using a pixel classifier on the DAB channel (38). The data were normalized to the area of the region of interest (ROI) covering GRN.

#### Animal studies

##### Immunofluorescence (IF) and IHC analyses on mouse brain sections

IHC or IF on 10 µm thick sections from FFPE tissue was performed after deparaffinization and antigen retrieval in citrate buffer pH 6.0, essentially as described (33). Nonspecific binding was blocked by incubating the sections in 5% normal donkey serum in TBS (1 hour, RT). Then, the sections were incubated (overnight, at 4°C) with the following primary antibodies (also indicated in the relevant figure legends): phospho-S129 aSyn antibodies (rabbit mAb EP1536Y Abcam, #ab51253-1:400; rabbit mAb D1R1R, Cell Signaling #23706-1:400), Neuronal nuclei marker (NeuN, mouse mAb A60, Millipore #MAB377-1:1000), astroglial marker, glial fibrillary acidic protein (GFAP, chicken polyclonal, Abcam #4674-1:200), phagocyte marker CD68 (LAMP4, rat mAb FA-11, Novus Biologicals #NBP2-33337-1:200), and Sequestosome 1/p62 (guinea pig, Nordic Biosite GP62-C-1:200). For double IF co-detection, fluorophore conjugated secondary antibodies were used (Thermo Fisher: AlexaFluor488, AlexaFluor568, AlexaFluor647, 1:1000). For IHC, DAB chromogen detection was performed following prior incubation with biotin conjugated secondary antibody (anti-mouse, Sigma #B7264-1:100) and Extra-Avidin peroxidise (Sigma #E2886-1:200). Sections were counterstained with hematoxylin (Vector Labs, #H-3401).

For the image analyses, whole slide digital scans of the tissue sections were obtained using the Olympus VS120 upright microscope (at AU) equipped for brightfield scanning and fluorescence single-band emitters for DAPI, FITC, Cy3 and Cy5. High resolution IF views were exported using Qupath (v. 0.5.1), for further analysis with ImageJ. ROI for neurons (NeuN+) or microglia (CD68+) were identified using Cellpose (39), followed by manual corrections to determine cell profile numbers expressed as cell profiles/area in mm^2^. NeuN masks defined the neuronal area in each tissue section. P-aSyn (S129) and GFAP area fractions were defined by thresholding at the same level across all sections. P-aSyn (S129) area fraction was also quantified both within and outside neuronal masks (NeuN).

##### Mouse neuroanatomical topography

Panoramic view of digital slide scans were mapped onto Mouse Brain Atlas (Paxinos and Franklin’s The Mouse Brain in Stereotaxic Coordinates, Elsevier Publishing, 4^th^ Edition)- (40). Information about neuroanatomical tracts and nuclei in mouse CNS was primarily derived from The Mouse Nervous System (Elsevier Publishing, 1^st^ Edition)- (25).

##### Statistics

The data were statistically analyzed in Graphpad Prism software version 8, and the final graphs were prepared in Graphpad or Microsoft Excel. Statistical significance in datasets was calculated following the guidelines from relevant literature, as indicated in the figure legends. P values were set at: *p<0.05, **p<0.01, ***p<0.001, ****p<0.0001.

##### Data availability

All the data generated and analyzed during this study are included in the main manuscript and the associated supplementary files. A video montage of select behaviors (Table S2) is accessible on the figshare repository: https://doi.org/10.6084/m9.figshare.26936968

## RESULTS

In this study, we aimed to refine an experimental paradigm of *in vivo* aSyn aggregation for studying the significance of prodromal aSyn pathology in GRN, particularly in the context of sensorimotor phenotypes relevant to PD. Lewy related aSyn pathology affecting the GRN and nearby nuclei of reticular formation in PD has been reported previously, using silver staining methods and/or immunodetection of pathological aSyn deposition using antibodies/anti-sera (8, 9, 11). In post-mortem sections of medulla oblongata, obtained from cases with clinically diagnosed parkinsonism (Table S1), we assessed the phosphorylation of aSyn on the serine residue S129 (p-aSyn, S129), which is one of the widely used biochemical marker of aSyn pathology (6, 41, 42). Our data corroborate the findings from pioneer studies, such that we observed significant accumulation of p-aSyn (S129) in the GRN of PD cases compared to the controls (Fig. 1A-B; antibody, EP1536Y).

**Figure 1.**
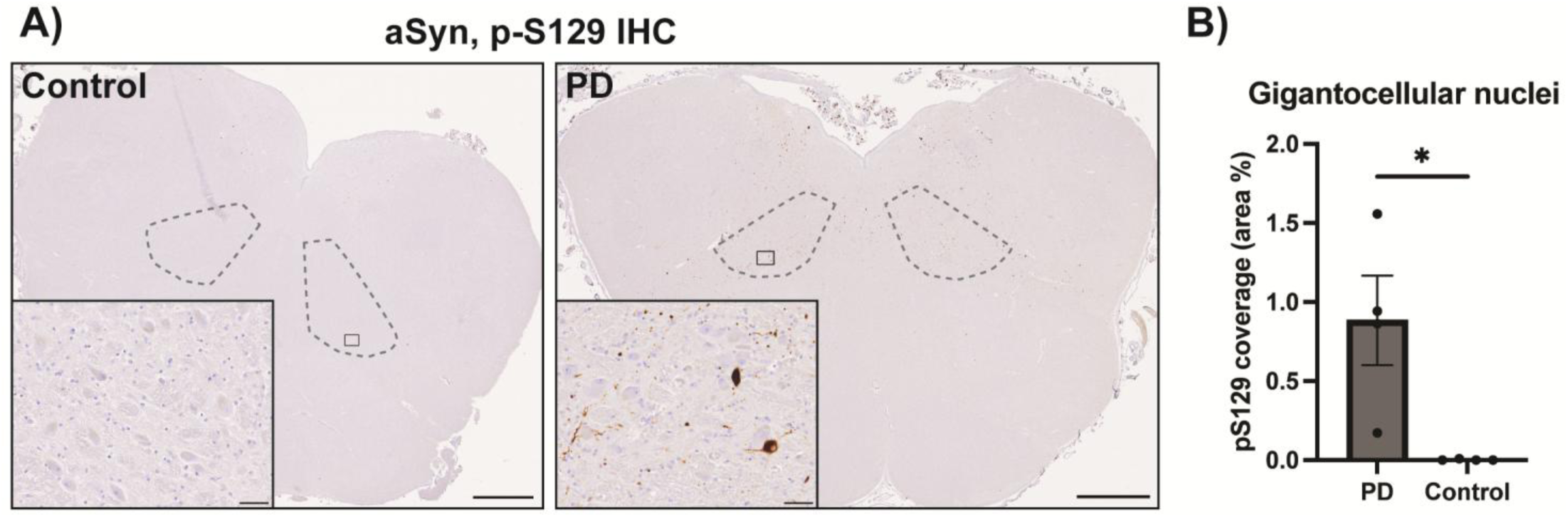
Immunostaining of phospho-alpha synuclein (p-aSyn, S129) in post-mortem human brain sections. **(A)** Representative images showing p-aSyn (S129) immunostaining in the transverse sections of medulla oblongata from controls and PD cases, with the gigantocellular nuclei (GRN) outlined by the dashed grey lines. Insets show high magnification images from the panoramic view, reflecting prominent Lewy pathology in PD and visible lack of staining in the controls. Scale bar = 2 mm (insets = 50 µm). Primary antibody in Figure 1A: p-aSyn (S129)-abcam EP1536Y. (B) Quantification shows that PD cases contain significantly more pS129-positive Lewy pathology than controls (p = 0.0286). Graph displays mean ± SEM of area % covered in pS129 staining. Groups were compared with a Mann-Whitney test, as they did not pass normality (Shapiro-Wilk test). *p < 0.05, n = 4 per group.

To evaluate the consequences of aSyn aggregation in GRN in rodent brain, we performed stereotaxic delivery of murine aSyn PFF into the pontine GRN of adult (12-14 weeks old) transgenic M83 mice, expressing the aggregation prone human mutant A53T aSyn (26). This approach (ie. delivery of exogenous PFF as seeding agents) for promoting *de novo* aSyn aggregation has been reproducibly used for studying the effects of aSyn-induced proteopathic stress in the nervous system, in both transgenic models and in wild type rodents (33, 43–48). Accordingly, we designed an experimental protocol in which we incorporated longitudinal assessment of select sensorimotor behaviors in heterozygous M83^+/-^ mice, following PFF-mediated induction of aSyn aggregation in brain, with GRN as the initial nidus (Fig. 2A). Our main cohort consisted of hetereozygous M83^+/-^ mice injected with PFF aSyn (n=18), while monomeric aSyn and PBS vehicle injection were used as controls (n=5/group). Using p-aSyn (S129) as a surrogate marker of cellular aSyn pathology (6, 41, 42), we also studied the induction of aSyn aggregation in GRN and its propagation into additional brain regions over time.

**Figure 2.**
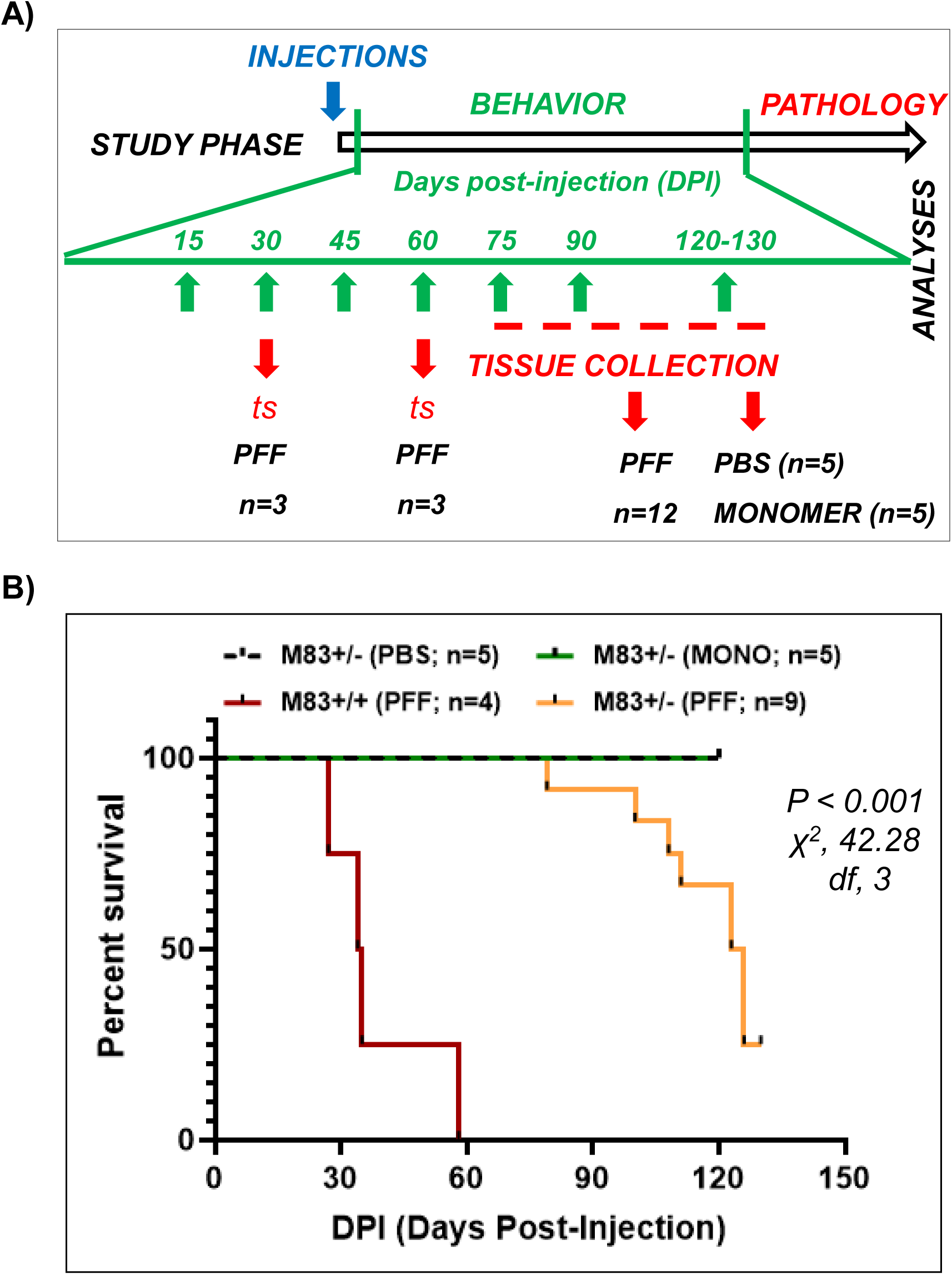
Overview of the study design in cohorts of heterozygous M83^+\-^ mice. **(A)** In cohorts of transgenic M83^+\-^ mice (both male and female. 12-14 weeks of age), phosphate buffered saline (PBS) vehicle (n=5), monomeric (ie., non-aggregated) aSyn (n=5) or pre-formed fibrillar (PFF) aSyn (n=18) were sterotaxically delivered bilaterally in the pontine GRN/Gi. Post-recovery, the mice were periodically assessed over time in tests of sensorimotor behaviors (green arrows, median interval between measurements: 15 days, unless indicated otherwise in the figure legends). Timed tissue collection was performed from the PFF-injected cohort after sacrifice at 30 and 60 days post-injection (DPI) (n=3/time-point), or in the event of a humane endpoint (period is highlighted by the dashed red line). The PBS and monomer cohorts were euthanized at the termination of the study (Between DPI-120 and DPI-130). **(B)** Kaplan–Meier plot showing survival of transgenic M83^+/-^ mice injected with PBS, monomeric aSyn or PFF aSyn bilaterally in the pontine GRN/Gi. The survival analyses do not include mice from the M83^+/-^ PFF aSyn cohort terminated at DPI-30 and DPI-60 (timed-sacrifice, mentioned in A). The median time to moribund state for the heterozygous M83^+/-^ was 124.5 days post-injection (9 out of 12 animals reached terminal stage by DPI-129; the 3 remaining animals were euthanized at DPI-130 and not included/censored in the survival analyses). A small cohort of homozygous M83^+/+^ mice (n=4) was also studied for comparison, in which the median time to moribund state was 34.5 days. The PBS and monomeric injected mice M83^+/-^ remained comparatively asymptomatic in gross motor performance (ie. no foot-drop, paralysis or significant weight loss) over the duration of the experiment. Statistics in B: Log-rank Mantel-cox test (M83^+/-^ cohorts: PBS, n=5; monomeric aSyn, n=5; PFF aSyn, n=12 and M83^+/+^ cohort: n=4; \*\*\*\**p*<0.0001; *^2^*, 42.28; *df*, 3).

Here, we show that PFF aSyn delivery in rodent GRN leads to the emergence of unique sensorimotor phenotypes, including altered patterns of spontaneous activity and progressive deterioration of motor performance in association with cellular aSyn pathology. In brief, 9 out of the 12 PFF-injected heterozygous M83^+/-^ mice reached a terminal stage within 129 days post-injection (DPI-129), with median time to moribund state of 124.5 days (Fig. 2B). In comparison, the control M83^+/-^ cohorts (PBS and monomeric aSyn) remained asymptomatic in gross appearance (ie. no foot-drop, freezing or significant weight loss) over the duration of the experiment. Prior to initiating the study in heterozygous M83^+/-^ (Fig. 2A), we performed a pilot study in cohorts of homozygous M83^+/+^ mice (n=4; 12-16 weeks old), in which PFF aSyn delivery was associated with a highly unfavorable outcome with median time to a moribund state of 34.5 days (Fig. 2B). Albeit, the pilot study (using M83^+/+^ mice) lacked appropriate controls, and a detailed behavioral characterization was not performed. Nevertheless, these observations are in congruence with previous reports that PFF aSyn delivery in transgenic M83 mice is associated with progressive motor impairment and reduced survival, with significantly earlier onset in the homozygous mice compared with the heterozygous animals (33, 47, 49).

### Direct PFF aSyn delivery in GRN leads to an early-onset phenotype characterized by progressive decline in spontaneous activity and decreased locomotion in M83^+/-^ mice

In order to assess the effects of aSyn aggregation in GRN on spontaneous activity and locomotion in M83^+/-^ mice, we employed non-invasive 24-hour monitoring using digitally ventilated cages (DVC, Tecniplast). For this purpose, we tracked the animals’ activity (DVC locomotion index, see Methods) over a longitudinal period, starting at DPI-36 till DPI-90. Our results show a progressive decline in the spontaneous locomotion of PFF aSyn-injected M83^+/-^ mice (n=12, in 4 cages), which could be clearly distinguished from the controls by DPI-60 (Fig. 3A; compare PBS and monomeric aSyn; n=5/in 2 cages/group). Moreover, evaluation of daily rhythms and home cage activity during light/dark periods revealed that the PFF-injected M83^+/-^ mice were significantly less active during the dark period (night time, active time for rodents), compared to the controls (PBS and monomeric aSyn; Fig. 3B-C; complete data on daily rhythms are presented in Fig. S1).

**Figure 3.**
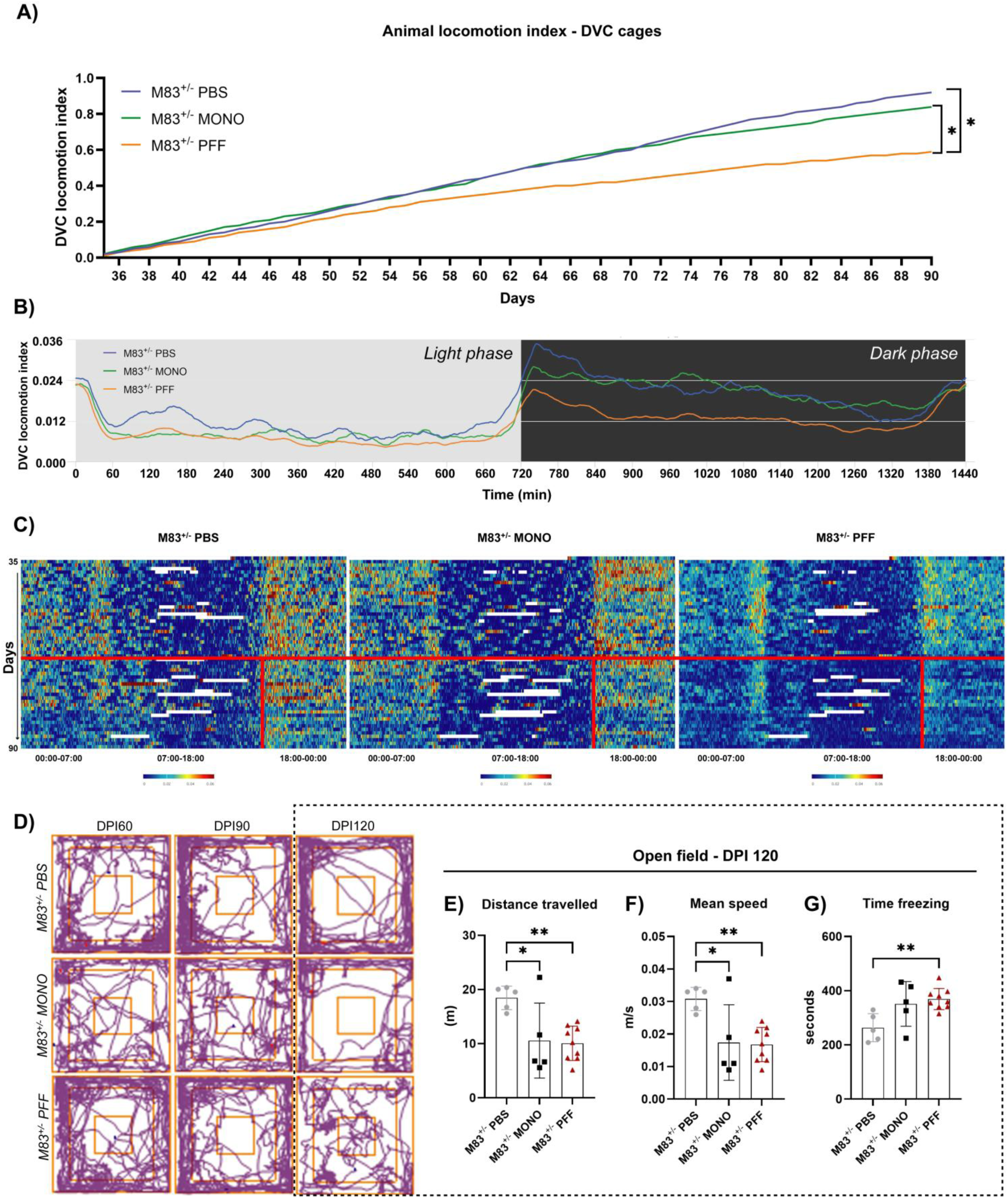
Spontaneous activity and locomotion in cohorts of heterozygous M83^+\-^ mice. **(A-C)** Non-invasive monitoring of spontaneous activity by cohorts of M83^+/-^ mice in digitally ventilated cages (DVC) over a longitudinal period up to 56 days post-injection (DPI) starting at DPI-36. The line chart (in A) represents the cumulative locomotion index (x-axis, days), which is represented for light and dark periods (in B, x-axis showing time in minutes) and heatmaps (in C, x-axis showing time in hours over 24-hour period each day, y-axis showing days with DPI-36 as the starting point on top). Also see line charts in S1A-D displaying daily locomotion index for each day, including light and dark periods. Statistics in 2A: One-Way ANOVA followed by Dunn’s multiple column comparisons (PBS, n=5 housed in 2 cages; monomeric aSyn, n=5 housed in 2 cages and PFF aSyn, n=12 housed in 4 cages; *p<0.05; Error bars, Mean ± SD). Only significant differences are highlighted. (**D-G**) Measurements of spontaneous activity by cohorts of M83^+/-^ mice in the open-field arena recorded over 10 minutes (ANY-maze, see Material and Methods) and representative tracking plots (in D) at the indicated time points (Days post-injection: DPI-60, DPI-90 and DPI-120. Bar graphs with individual points represent quantitative measurements of the overall distance travelled (in E), mean speed (in F) and time freezing (in G). Statistics in 2E-G: One-Way ANOVA followed by Tukey’s multiple column comparisons (PBS, n=5; monomeric aSyn, n=5 and PFF aSyn, n=9; *p<0.05, **p<0.01; Error bars, Mean ± SD). Only significant differences are highlighted. Also see Fig. S2A-B for the measurements in the open field arena at time-points DPI-60 and DPI-90.

In parallel, we also assessed the locomotion pattern and exploratory behavior of the mice in open-field arena at DPI-60, DPI-90 and DPI-120. Intriguingly, while the findings in DVC suggest a progressive reduction in activity by the PFF aSyn-injected cohort as early as DPI-60, all the experimental groups showed similar pattern of activity in the open-field arena at DPI-60 and DPI-90 (Fig. S2A-B). However, at the advanced stage of the study (DPI-120), distinct locomotion patterns between the groups could be distinguished (Fig. 3D-G; S2C). Near the termination, the PFF and monomeric aSyn-injected mice moved less distance (Fig. 3E), were overall slower (Fig. 3F), and exhibited frequent freezing episodes compared to the PBS cohort. Intriguingly, we also observed some differences in the entries to periphery or center of the arena between the monomeric aSyn and PFF aSyn injected animals (Fig. S2B-C). A video montage of the open-field behavior is presented in the supplementary files accessible on the figshare repository (Table S2): VIDEO 1-5 (M83^+/-^) and VIDEO 18 (M83^+/+^)- see Data availability.

### PFF aSyn-injected M83^+/-^ mice exhibit progressive defects in fine control of posture and balance

The experimental cohorts of M83^+/-^ mice were also subjected to longitudinal assessment of motor coordination and balance using a battery of standard tests (13, 50–52). During these assessments, PFF aSyn-injected M83^+/-^ mice exhibited progressive deterioration of performance in the balancing beam test (Fig. 4A-C). These deficits were observed as early as DPI-45 on the narrow beam (diameter: 8 mm), such that the animals took longer time to traverse, experienced frequent slips of the hindpaws and had overall slower speed of movement, compared to the controls (Fig. 4A). Strikingly, the performance of all cohorts was relatively comparable on the wider beam (diameter: 16 cm) till DPI-90 (Fig. 4B), suggesting relatively intact postural reflexes and lack of gross motor weakness. The latter is also corroborated by the findings in the grip strength test (S2D), which ruled out major defects in motor strength and the ability to grab and hold surfaces. In the balance beam test, we also observed freezing of movement in the PFF aSyn-injected M83^+/-^ mice, such that the mice would hesitate to initiate the traversal and adapt a hunched posture (Fig. 4C). Near the terminal stage (DPI-120), slower movement and frequent hindpaw slips were eventually observed on the wider beam as well (Fig. 4B). Thus, these data (Fig. 4A-C) indicate a progressive nature of the deficits in fine control of postural adaptations in the PFF aSyn-injected M83^+/-^ mice. A video montage of the performance in balance beam test is presented in the supplementary files accessible on the figshare repository (Table S2): VIDEO 6-9 (M83^+/-^) and VIDEO 19 (M83^+/+^)- see Data availability.

**Figure 4.**
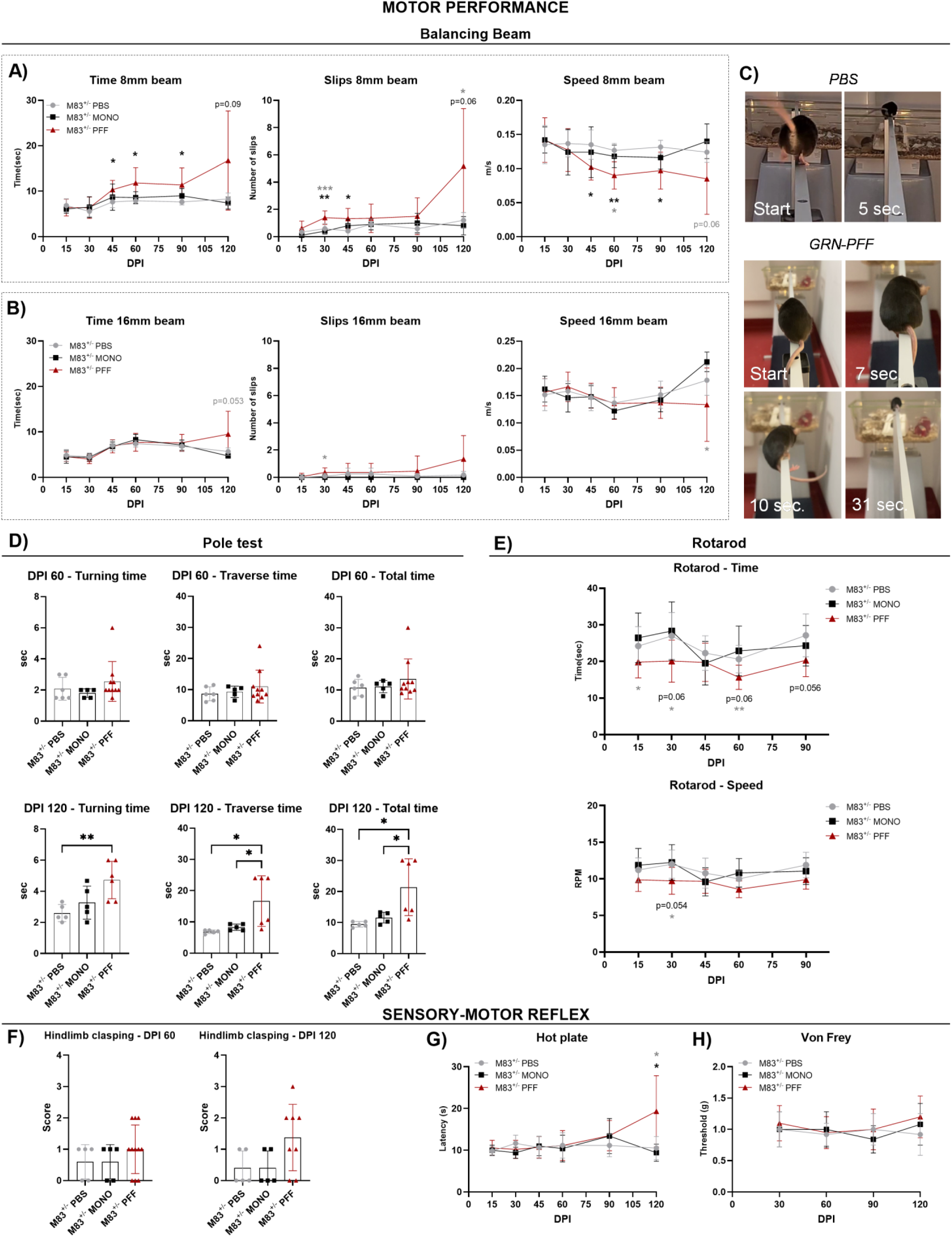
Measurements of sensorimotor behaviors in cohorts of heterozygous M83^+\-^ mice. **(A-B)** Assessment of motor coordination of M83^+/-^ cohorts (injected with PBS, monomeric aSyn or PFF aSyn) in the balancing beam test over a longitudinal period (shown on x-axis as DPI, days post-injection). The line graphs show the traversal time (seconds), number of slips and speed (m/s) on 8mm wide beam (in A) and 16mm wide beam (in B). **(C)** Representative images showing postural adaptations and performance on 8mm beam by PBS or PFF-injected M83^+/-^ mice. **(D)** Bar graphs depicting performance of M83^+/-^ cohorts (injected with PBS, monomeric aSyn or PFF aSyn) in the pole test with individual points representing the quantitative measurement of the turning time, traversal time and total time on pole at DPI-60 and DPI-120 (DPI, days post-injection). **(E)** Assessment of motor coordination of M83^+/-^ cohorts (injected with PBS, monomeric aSyn or PFF aSyn) in the rotarod test over a longitudinal period (shown on x-axis as DPI, days post-injection). The line graphs depict the quantitative measures of the latency to fall (time, in seconds) and speed at fall (rotations per minute, RPM). **(F)** Postural reflex assessment in modified tail-suspension test with bar graphs depicting hindlimb clasping by cohorts of M83^+/-^ mice (injected with PBS, monomeric aSyn or PFF aSyn) at DPI-60 and DPI-120 (DPI, days post-injection). **(G-H)** Assessment of thermal nociception (hot plate, in G) and mechanical allodynia (von frey, in H) in cohorts of M83^+/-^ mice (injected with PBS, monomeric aSyn or PFF aSyn). Statistics in 3A-H: One-Way ANOVA followed by Tukey’s multiple column comparisons (PBS, n=5; monomeric aSyn, n=5 and PFF aSyn, n=6-9; *p<0.05, **p<0.01; Error bars, Mean ± SD). Black asterisks (all panels) represent significant differences between PBS and PFF aSyn-injected cohorts, while the grey asterisks (in A and G) indicate significant differences between monomeric aSyn and PFF aSyn-injected cohorts. Only significant differences are highlighted.

### Impaired performance in complex sensorimotor tasks and defective response to nociceptive stimuli is a late-stage phenotype in the PFF aSyn-injected M83^+/-^ mice

We also subjected the experimental cohorts of M83^+/-^ mice to tests requiring more complex sensorimotor skills and motor coordination. Among these, the pole test and the accelerating rotarod are commonly employed methods of phenotype assessment in models of basal ganglia disorders, including aSyn overexpressing rodents (51, 52). In the pole test, the performance of experimental cohorts was overall comparable during the early stage of the study (DPI-60) (Fig. 4D, top panel). In contrast, significant differences between PFF aSyn-injected M83^+/-^ mice and controls were seen near the terminal stage (DPI-120; Fig. 4D, bottom panel). At DPI-120, the PFF aSyn cohort exhibited increased latency in the turning response; albeit, the increased latency in descending and more time spent on the pole is due to assigning the cut-off test values to 3 out of 6 mice (see Methods). A video montage of the performance in the pole test is presented in the supplementary files accessible on the figshare repository (Table S2): VIDEO 10-13 (M83^+/-^)- see Data availability. The severity of movement incoordination was further substantiated by the findings in rotarod test, in which the performance of PFF aSyn cohort at DPI≥60 was significantly impaired in comparison with the controls (Fig. 4E; latency to fall, speed at fall).

Taken in conjunction, these findings indicate that gross defects in movement coordination following PFF aSyn delivery in the GRN are features defining the advanced stages of motor disability. This notion is further supported by assessments in the footprint test (gait stance) and additional tasks requiring sensorimotor coordination (hindlimb clasping and response to nociceptive stimuli- see below). In the footprint test, we did not observe significant alterations in stride length, step width and base of support measurement between the cohorts (Fig. S3B-C) However, there were signs of uncoordinated gait at the pre-terminal stage (1-2 days before euthanasia, DPI≥106) in 4 PFF aSyn-injected mice, reflected by the changes in step alternation during locomotion (Fig. S3B, S3D; compare instances indicated by green arrows and red arrows, reflecting coordinated gait or gait incoordination respectively). In one female (euthanized DPI-123), signs of imminent unilateral foot-drop could also be seen (Fig. S3B).

Another prominent phenotype in M83 mice, reported as a harbinger of motor collapse in the presence of established brainstem aSyn pathology, is moderate-to-severe degree of hindlimb clasping (33, 53, 54). This sensorimotor reflex is reliant upon an intact ocular-postural-motor coordination, and is also imapired in models of basal ganglia, cerebellar or motor neuron dysfunction (55). In the early stage (DPI-60), we observed a normal response (score 0) or mild degree (score 1) of hindlimb clasping in the controls, and 50% of the PFF aSyn-injected mice (Fig. 4F). Around DPI-120, ∼30% animals in the PFF aSyn cohort progressed to moderate-severe degree of clasping (Fig. 4F; scores 2-3), suggesting potential dysfunction in pathways mediating this reflex response. A video montage of the hindlimb clasping test is presented in the supplementary files accessible on the figshare repository (Table S2): VIDEO 14-17 (M83^+/-^) and VIDEO 20 (M83^+/+^)- see Data availability.

Lastly, a previous study suggested that PFF aSyn delivery, using intramuscular injections, impaired nociception and mechanical allodynia in M83 mice (27). Therefore, we also assessed sensorimotor reflexes of nociception/allodynia using thermal or tactile stimuli (Hot plate and Von Frey filaments, respectively). In these tests, we observed that all experimental cohorts responded similarly over the duration of the study (ie. exhibited an intact nociceptive response, in Fig. 4G-H). Although, we observed a relatively increased latency in the Hot plate in the PFF aSyn-injected cohort around DPI-120 (Fig. 4G), we consider that these results most likely reflect defects in complex movement coordination at this stage, rather than a pure sensory defect in nociception.

### PFF aSyn delivery induces *de novo* aSyn aggregation in GRN with spatiotemporal progression in brainstem

In addition to the behavioral characterization, we wanted to contextualize the nature of neuronal aSyn pathology that potentially underlies the early and late phenotypes in movement coordination in M83^+/-^ mice, following PFF aSyn delivery in GRN (Fig. 3-4). Accordingly, we assessed the emergence and spatiotemporal progression of aSyn pathology in select brain regions (Fig. S4). Our analyses were guided by previous studies in the M83 mice, showing substantial p-aSyn (S129) accumulation (spontaneous age-related, or PFF-induced) in the nuclei of reticular formation including GRN, midbrain PAG and vestibular nuclei (VN) (26, 33, 43, 47, 49). Moreover, a mild-moderate degree of aSyn aggregation also has been detected in the deep cerebellar nuclei (DCN), red nucleus (RN) and motor cortex (M1/M2). Notably, pathological involvement of substantia nigra (SN), striatum (caudatoputamen, CPu) and parts of thalamus is rarely observed in the M83 line (26, 33, 43, 47, 49).

By IF analyses (antibody: EP1536Y), we detected a sparse degree of p-aSyn (S129) accumulation in the GRN of PBS-injected mice at DPI-120 (Fig. 5A, 6B), likely due to the spontaneous aSyn aggregation in the ageing transgenic M83^+/-^ mice. In contrast, substantially more p-aSyn (S129) was detected in the GRN of monomeric aSyn-injected cohort at DPI-120, in both neuronal cell bodies and in the surrounding neuropil (Fig. 5B, 6B). In the PFF aSyn injected mice, localized p-aSyn (S129) was detected within GRN and adjacent pontine reticular formation as early as DPI-30 (Fig. 5C, 6A-B, S7B), and appeared to predominantly affect the neuropil (ie. based on minimal overlap with neuronal nuclei marker, NeuN; also see, Fig. S7A). Outside the GRN, we did not detect appreciable degree of p-aSyn (S129) accumulation at this early stage (Fig. 6A, S5A-E, S7B; regions shown in S5: VN, DCN, RN, PAG, and subthalamic nucleus-STh). By DPI-60, there was further increase in p-aSyn (S129) within GRN, characterized by a mixed pattern of accumulation in neuronal perikarya and surrounding neuropil (Fig. 5C, 6A-B, S7A-B). Among the other regions examined, p-aSyn (S129) was detected in the VN, and to a lesser extent in the DCN, RN, PAG and STh at this stage (Fig. 6A, S5A-E, S7B). The progressive nature of PFF aSyn-induced pathology was further corroborated by the analyses at terminal stage (DPI≥108), which revealed significantly increased detection of p-aSyn (S129) not only in the GRN (Fig. 5C, 6A-B, S7B) but also in additional brain regions. Among the latter, nuclei within pons, cerebellum and midbrain showed a higher degree of cellular p-aSyn (S129) accumulation, while STh and some regions in thalamus/hypothalamus were weakly immunopositive (Fig. 6A, S5A-E, S6A, S7B). Lastly, in line with the previous reports, there was visible lack of p-aSyn (S129) accumulation in M1/M2, Cpu and SN (S6A).

**Figure 5.**
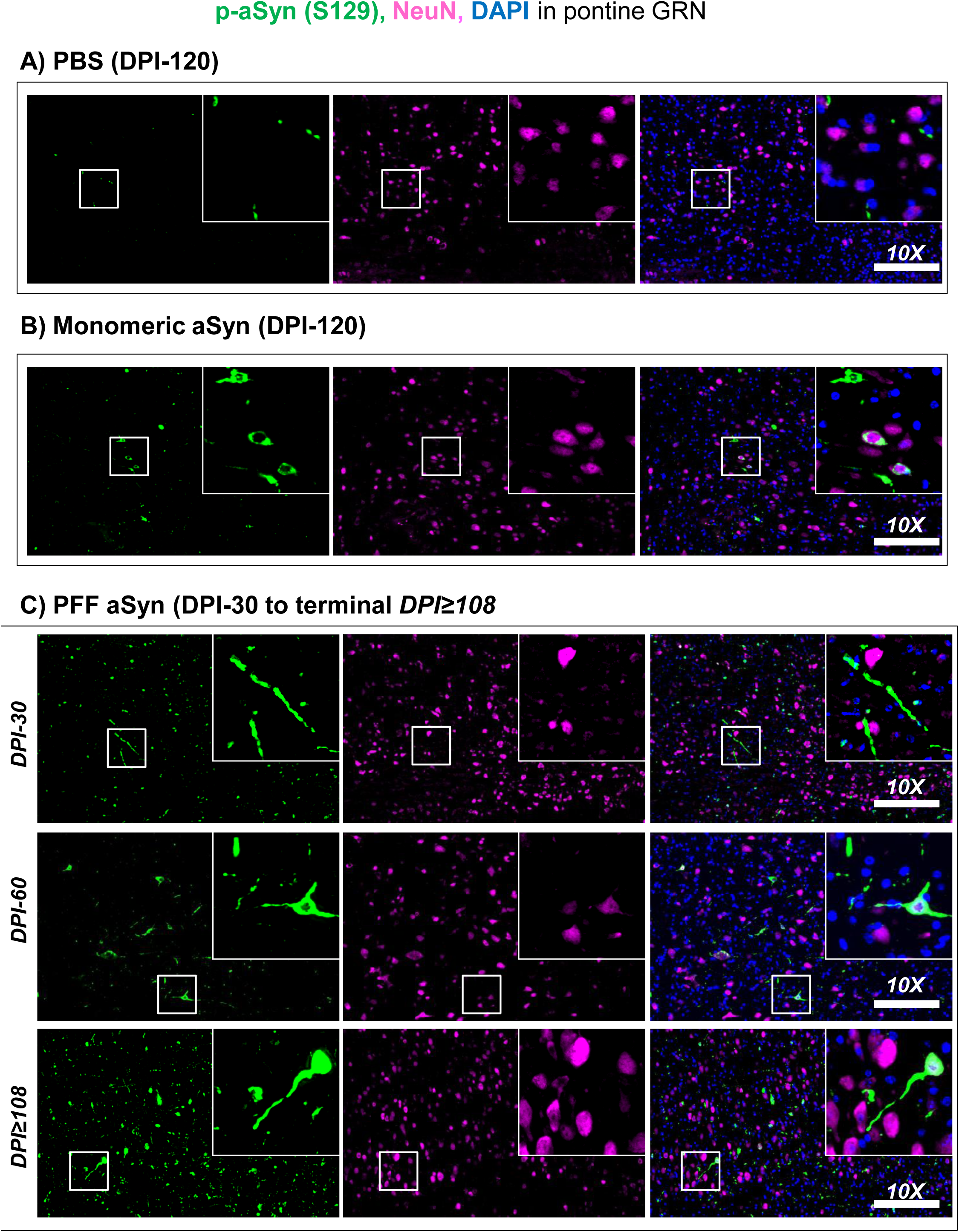
Dual immunofluorescence detection of phosphorylated alpha-synuclein (p-aSyn, S129) and neuronal nuclei (NeuN) in the GRN of young M83^+\-^ mice. **(A-C)** Representative low magnification (10X) images showing p-aSyn (S129) and neuronal nuclei (NeuN) immunofluorescence detection in the GRN/Gi of PBS (in A), monomeric aSyn (in B) and PFF aSyn (in C) injected M83^+/-^ mice at indicated time-points (DPI, days post-injection). The insets show 63X magnified views of the GRN/Gi, highlighted by the white-bordered square in the 10X views (scale bar= 200 µm). Notice the neuritic and cellular aSyn pathology in monomeric aSyn-injected cohort (DPI-120, in B) and the PFF aSyn-injected cohort, which increases over time in the latter (DPI-30 to terminal stage DPI≥108; also see Fig. 5B and S7A). A sparse degree of spontaneous aSyn pathology was also seen in the PBS-injected cohort (in 4A, DPI-120, age 7-8 months). Also see Supplementary Figures S5 (additional brain regions with p-aSyn, S129 positivity), S6-7 (markers of inflammatory gliosis) and S8 (p-aSyn S129 detection in the terminal-stage homozygous M83^+/+^ mice, ≥DPI-34). Primary antibodies in Figure 4A-C: p-aSyn (S129)-abcam EP1536Y and NeuN-EMD Millipore A60.

**Figure 6.**
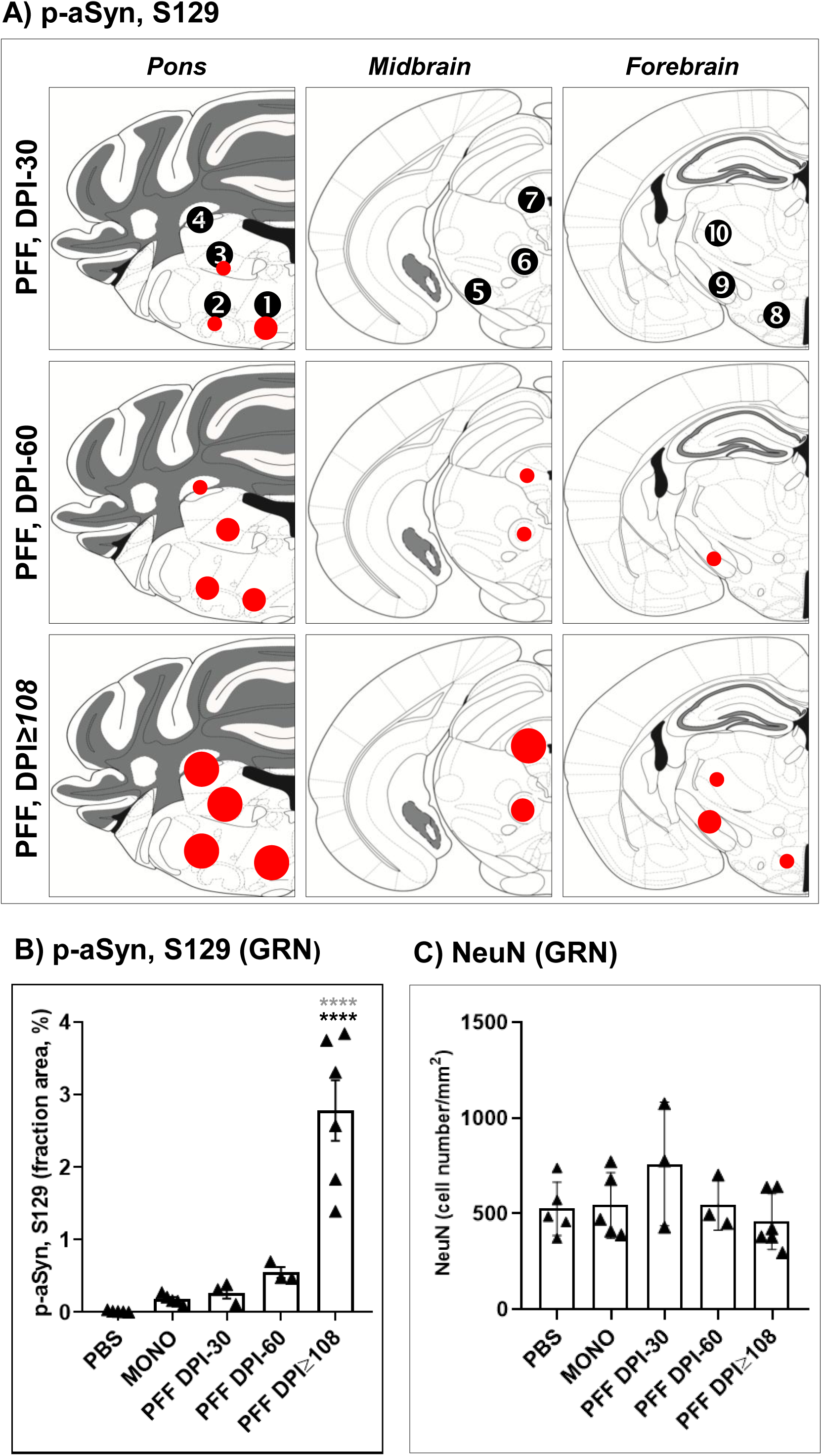
Composite schematic depiction of phosphorylated alpha-synuclein (p-aSyn, S129) immunofluorescence detection in the brains of M83^+/-^ mice. **(A)** Average p-aSyn (S129) fluorescence signal (percentage of area) in the regions is highlighted by colored circles as follows: small circle (0.05-0.2%), medium circle (0.2-1%) and large circle (1-5%). Also see Supplementary Figure S4 for the low magnification (2X) panoramic views annotating select brain regions and S7B. Regions shown in 5A: (*view in S4, Pons*) ❶ gigantocellular nuclei (GRN), ❷ pontine reticular nuclei (Rt), ❸ vestibular nuclei; ❹ deep cerebellar nuclei; (*view in S4, Midbrain*) ❺ substantia nigra, ❻ red nucleus, ❼ periaqueductal grey matter; (*view in S4, Thalamus*) ❽ medial and lateral hypothalamus, ❾ subthalamic nucleus, ❿ ventromedial and ventrolateral thalamic nuclei. **(B)** Quantitative estimates of p-aSyn S129 immunopositivity (immunofluorescence signal intensity from two serial sections) in the GRN/Gi in cohorts of M83^+/+^ mice, expressed as percentage of the area (total average area/image: 625,000 um^2^) in 10X views (see Fig. 4A-C). **(C)** Quantitative estimates of the total number of neurons (NeuN immunofluorescence) in of GRN/Gi from cohorts of M83^+/-^ mice. Statistics in 5B: One-Way ANOVA followed by Tukey’s multiple column comparisons (PBS, n=5; monomeric aSyn, n=5; PFF aSyn DPI-30 and DPI-60, n=3 and PFF aSyn DPI≥108, n=5; ****p<0.0001; Error bars, Mean ± SD). Black asterisks represent significant differences between PBS and PFF aSyn-injected cohorts, while the grey asterisks indicate significant differences between monomeric aSyn and PFF aSyn-injected cohorts. Only significant differences are highlighted.

### PFF aSyn-induced aSyn aggregation in GRN of M83^+/-^ mice is associated with neuroinflammation

To further characterize the cellular aSyn pathology, we performed dual IF detection of p-aSyn (S129) with p62 protein (Sequestosome 1, a marker of intracellular protein aggregates and autophagy), or markers of inflammatory gliosis in the PFF aSyn-injected cohort. For this purpose, we restricted our analyses to GRN (initiation site in pons), PAG (a propagation site in midbrain) and M1/M2 (motor cortex, with minimal aSyn aggregation) in the terminal stage animals (DPI≥108). As expected, p-aSyn (S129) aggregation within GRN and PAG was associated with substantial p62 accumulation, refelectd by IF co-detection of the two markers (Fig. S6B; compare M1/M2). In parallel, we also probed reactive astrogliosis and microglial infiltration, which have been reported as neuropathological features in the brains of M83 mice with advanced aSyn pathology (47, 49). In line with these reports, we found increased expression of astroglial cells (GFAP, in Fig. S6C and S7C), and phagocytic microglia (CD68, Fig. S6D and S7D) in the disease affected regions (GRN and PAG). In addition, we assessed the degree of neuronal loss in GRN, as a potential underlying factor for the observed sensorimotor phenotypes in the PFF aSyn-injected cohort. Our analyses suggest a slight (albeit, non-significant) decrease in the total number of neurons in the GRN and adjacent part of the pontine reticular formation (Fig. 6C).

In addition to the detailed quantitative studies in heterozygous M83^+/-^ cohorts presented above, we also examined p-aSyn (S129) accumulation in select brain regions of terminal stage homozygous M83^+/+^ mice, following aSyn PFF delivery in the GRN (pilot study, Fig. 2B). While we have not performed detailed quantitative assessments, IF and IHC (DAB chromogen) suggested a similar pattern of regional p-aSyn (S129) distribution, such that the GRN and the nearby VN (in pons) and PAG (in midbrain) were among the most conspicuously affected regions (Fig. S8A-D). Moreover, motor cortex (M1/M2) was largely devoid of p-aSyn (S129), as suggested by a sparse IF signal (Fig. S8E). Lastly, we aimed to rule out any bias in p-aSyn (S129) detection, which could potentially emanate from reliance on one antibody (EP1536Y) used in the main analyses (Fig. 5-6, S5-S7). To this end, we employed a different polyclonal antibody (rabbit, D1R1R), and assessed p-aSyn (S129) immunostaining in the GRN of the terminal stage heterozygous M83^+/-^ mice. By IF and IHC (DAB), both antibodies detected characteristic pattern of cellular p-aSyn (S129) immunostaining in the GRN, ie. in neuronal perikarya and neuropil (Fig. S8A-B). Taken together, these analyses (S7-S8) are compelling indicators of the pathological nature of p-aSyn (S129) accumulation in the brains of M83 mice, following PFF aSyn delivery in the GRN.

## DISCUSSION

Progressive movement disability, in the form of difficulty in movement initiation (bradykinesia) and impaired postural reflexes (shuffling gait, difficulty in turning) are among the cardinal features of clinical parkinsonism. It is also recognized that pathological aSyn accumulation affecting distinct extra-nigral loci in nervous system may precede the onset of clinical symptoms by several years, and depending on the initial site of aggregation, may dictate the course of disease progression (1, 3, 7, 10, 56–58). Historically, the significance of extra-nigral brainstem aSyn pathology- to a large extent- has been investigated in the context of non-motor symptoms (12–15, 50, 59). Hence, experimental models of extra-nigral brainstem aSyn pathology in the context of PD motor symptomatology are few and far between.

In this study, we investigated the significance of pathological aSyn aggregation in neuronal populations of GRN in relation to the features of movement disability in PD. Our data show that young heterozygous M83^+/-^ mice with aSyn aggregation (predominantly) localized to GRN (Fig. 5C, 6A-B, S7B), exhibit an early phenotype characterized by reduction in spontaneous locomotion (Fig. 3A-B, DVC), and concomitant defects in fine control of postural reflexes (Fig. 4A-C, narrow beam). However, gross defects in locomotion patterns (Fig. 3D-G, open field) and signs of incoordination in complex sensorimotor tasks (Fig. 4D-F) emerged at a later stage, when multi-focal aSyn pathology and inflammation was detected (S5-S7). Collectively, the spatiotemporal progression of cellular aSyn pathology and emergent sensorimotor phenotypes are encouraging findings for further refining this experimental paradigm (discussed below), towards establishing the pathogenic significance of aSyn aggregation in GRN.

Nevertheless, there are some key limitations that preclude major conclusions from the study presented above. The most obvious limitation is the small sample size and lack of detailed analyses for the controls at earlier time points (DPI-30 and DPI-60), especially monomeric aSyn injected animals. Close to the termination of the study (DPI-120), these animals also started to exhibit slight (although, non-significant) decline in the spontaneous activity (Fig. 3A-C) and altered patterns of locomotion in the open-filed (Fig. 3D-G), in association with localized aSyn pathology in GRN (Fig. 5B, 6B). Future studies could exploit this relatively slower aggregation paradigm and may yield better insights into the natural history of motor phenotypes in the context of progression of aSyn pathology in this (GRN synucleinopathy) model. Another major limitation of the study is that the observations have been made in transgenic M83 mice (overexpressing the aggregation prone, human mutant A53T aSyn), with an aggressive approach for inducing aSyn pathology (ie. PFF aSyn delivery in the GRN). It is well established that naive M83 mice (ie. without exogenous PFF aSyn injections) also exhibit age-related decline in locomotion, impaired performance in rotarod and a severe moribund phenotype near the terminal stage (lack of grooming, freezing, foot-drop and paralysis) (26, 47). However, it is noteworthy that the phenotypes reported in naive M83 mice appear at much later ages than the animals used in our study, ie. 8-12 months in homozygous M83^+/+^ and 20-24 months in heterozygous M83^+/-^ mice (26, 47).

It has also been reported that the onset and progression of the motor phenotypes in M83 mice are significantly exacerbated by exogenous delivery of PFF aSyn by peripheral routes, which results in initial aSyn pathology in spinal cord and subsequently across the neuraxis (43, 47, 49, 60). In other words, the movement disability in M83 mice following peripheral PFF delivery in these prior studies is not attributed to aSyn pathology in a distinct brain region(s), but largely considered to emanate from the loss of spinal motor neurons (47, 49). In contrast, our observations suggest that the early features of movement disability in the PFF aSyn-injected cohort (bradykinesia and postural instability, Fig. 3A-C, 4A-C) emerge in association with predominantly localized aSyn pathology in GRN (Fig. 5C, 6A-B, S7B). Moreover, our data also preclude widespread dysfunction in somatomotor system, since overall locomotion (open field, Fig. 3D-G), movement coordination (performance on wider beam, in Fig. 4B; pole test, in Fig. 4D; rotarod, in Fig. 4E) remained relatively intact till very late in the study (≥DPI-90). This conclusion (ie. aSyn aggregation in GRN leads to early phenotypes of bradykinesia) is also supported by the observations in monomeric aSyn injected M83^+/-^ mice, which exhibited altered locomotion patterns around DPI-120 (Fig. 3A-G) in association with incipient aSyn aggregation (5B, S7B; compare to PBS).

Nevertheless, given the similarities in the distribution of advanced aSyn pathology reported previously in M83 mice and our study, including the lack of involvement of SN and Cpu (26, 43, 49), a refined approach to validate this model (of aSyn aggregation in GRN) is warranted. As mentioned above, it is important that the behavioral phenotypes (bradykinesia and postural instability) are assessed in an experimental paradigm with localized and relatively slower propagation of aSyn aggregation in GRN, instead of the aggressive PFF aSyn-based approach used in this proof-of-concept study. This could be achieved using direct delivery of monomeric aSyn (eg., comparing wild type and mutant A53T aSyn) or ectopic expression of human aSyn transgene in GRN using viral vectors (13, 50). This later approach is also advantageous, since it could help dissect dysfunction in distinct neuronal sub-populations in the GRN, which potentially underlie the movement disability in this model. Technically, this is feasible in genetically modified rodents in which human transgene(s) expression can be achieved in target neuronal populations by the stereotaxic delivery of engineered viral vectors, as demonstrated for cre-recombinase expressing dopamingergic neurons *in vivo* (61). Thus, we consider that the validation of this model of neurodegenerative aggregate pathology in GRN, and further refinements to this experimental paradigm hold promise for novel translational approaches in PD and related disorders. To illustrate, recent studies show that chemogenetic activation of *Chx10* neurons in the GRN helps restore turning deficits in a rodent model of toxin-induced dopamine depletion in the striatum (18).

In a larger context, several animal models based on aSyn overexpression (transgenic or through viral vectors delivery) have been studied and differ with regards to their fidelity in recapitulating features of movement disability in PD ((Reviewed elsewhere- (13, 50)). These studies show that the phenotypes in each model are also affected by the promoter driving aSyn expression, which in turn potentially dictates the spatiotemporal features of aSyn aggregation in distinct neuronal populations (26, 62, 63). For instance, transgenic models with aSyn expression (A53T or E46K) under the control of prion promoter (*Prnp*) exhibit severe motor phenotypes, most likely due to age-related decline in spinal motor neurons as mentioned above (26, 47, 49). In comparison, the transgenic lines based on the thymus cell antigen 1 (*Thy1*) promoter driven aSyn overexpression exhibit age-related progression of motor features, as well as altered circadian rhythm and sleep (64, 65). Intriguingly, we found that the pattern of spontaneous locomotion in PFF aSyn injected mice was considerably affected during the nocturnal phase (Fig. 3B-C). We do not know whether this phenotype of reduced nocturnal activity potentially reflects altered sleep pattern, possibly compounded by reduced locomotion, and is worth further investigations. Lastly, there are isolated reports in the published literature showing that monomeric aSyn injection in the GRN of wild type mice leads to progressive decline in dopamine concentration and tyrosine hydroxylase mRNA expression in the striatum; however, behavioral outcomes were not assessed/reported (66). Thus, characterizing the neurochemical aspects within GRN and associated subcortical motor circuitry in this model could further facilitate the identification of neural substrate(s) underlying features of movement disability in this model.

## Conclusion

In conclusion, we consider that our study highlights a crucial role of prodromal aSyn pathology in GRN to clinically relevant phenotypes of movement disability in PD. We also anticipate that our data will stimulate novel hypotheses in model development for dissecting the neurological basis of PD symptomatology.

## Supporting information

Theologidis and Jan et al., Supplementary Information

## DECLARATIONS, AS APPLICABLE

### Funding

This work was supported by funding to AJ in the form of a Michael J. Fox Foundation Investigator grant (grant# MJFF-021498). HG and PHJ was supported by Lundbeck Foundation grants R383-2022-18**0**, R248-2016-2518 for Danish Research Institute of Translational Neuroscience-DANDRITE, Nordic-EMBL Partnership for Molecular Medicine.

### Conflict of interest

The authors declare no conflict of interest.

## Acknowledgements.

The authors would like to thank the animal caretaker staff at the Department of Biomedicine-AU (DK) for assistance, and J.G.J.M. (John) Bol at UMC Amsterdam (NL) for histopathology on the human tissue sections.

## Author contributions

VT, SAF, MRR and AJ designed research; VT, SAF, NMJ, DMG, MR, MWH, IF, HG and AJ performed research; SAF, NMJ, OAA and EDR performed data analyses; WDJvdB provided immunostaining data on human post-mortem tissue and contributed to discussion; PHJ, PB, JRN, CBV and NVDB contributed with infrastructural and personnel support, data evaluation and discussion; VT, SAF and AJ wrote the manuscript. All the authors read and approved the current version of the manuscript.

## Notes

https://doi.org/10.6084/m9.figshare.26936968

### Competing Interest Statement

The authors have declared no competing interest.

## REFERENCES

1. L. V. Kalia, A. E. Lang, Parkinson’s disease. Lancet 386, 896–912 (2015).

2. W. Poewe et al., Parkinson disease. Nat Rev Dis Primers 3, 17013 (2017).

3. A. H. V. Schapira, K. R. Chaudhuri, P. Jenner, Non-motor features of Parkinson disease. Nat Rev Neurosci 18, 435–450 (2017).

4. G. U. Hoglinger et al., A biological classification of Parkinson’s disease: the SynNeurGe research diagnostic criteria. Lancet Neurol 23, 191–204 (2024).

5. D. W. Dickson, Neuropathology of Parkinson disease. Parkinsonism Relat Disord 46 Suppl 1, S30–S33 (2018).

6. M. Goedert, R. Jakes, M. G. Spillantini, The Synucleinopathies: Twenty Years On. J Parkinsons Dis 7, S51–S69 (2017).

7. P. Borghammer, The alpha-Synuclein Origin and Connectome Model (SOC Model) of Parkinson’s Disease: Explaining Motor Asymmetry, Non-Motor Phenotypes, and Cognitive Decline. J Parkinsons Dis 11, 455–474 (2021).

8. H. Braak et al., Staging of brain pathology related to sporadic Parkinson’s disease. Neurobiol Aging 24, 197–211 (2003).

9. H. Braak et al., Parkinson’s disease: affection of brain stem nuclei controlling premotor and motor neurons of the somatomotor system. Acta Neuropathol 99, 489–495 (2000).

10. K. A. Jellinger, Is Braak staging valid for all types of Parkinson’s disease? J Neural Transm (Vienna) 126, 423–431 (2019).

11. K. Seidel et al., The brainstem pathologies of Parkinson’s disease and dementia with Lewy bodies. Brain Pathol 25, 121–135 (2015).

12. L. M. Butkovich et al., Transgenic Mice Expressing Human alpha-Synuclein in Noradrenergic Neurons Develop Locus Ceruleus Pathology and Nonmotor Features of Parkinson’s Disease. J Neurosci 40, 7559–7576 (2020).

13. J. B. Koprich, L. V. Kalia, J. M. Brotchie, Animal models of alpha-synucleinopathy for Parkinson disease drug development. Nat Rev Neurosci 18, 515–529 (2017).

14. E. Menozzi, J. Macnaughtan, A. H. V. Schapira, The gut-brain axis and Parkinson disease: clinical and pathogenetic relevance. Ann Med 53, 611–625 (2021).

15. N. Van Den Berge et al., Ageing promotes pathological alpha-synuclein propagation and autonomic dysfunction in wild-type rats. Brain 144, 1853–1868 (2021).

16. D. E. Haines, G. A. Mihailoff, Fundamental neuroscience for basic and clinical applications (Elsevier, Philadelphia, PA, ed. Fifth edition., 2018), pp. xi, 516 pages.

17. J. M. Cregg et al., Brainstem neurons that command mammalian locomotor asymmetries. Nat Neurosci 23, 730–740 (2020).

18. J. M. Cregg, S. K. Sidhu, R. Leiras, O. Kiehn, Basal ganglia-spinal cord pathway that commands locomotor gait asymmetries in mice. Nat Neurosci 27, 716–727 (2024).

19. M. Lemieux, F. Bretzner, Glutamatergic neurons of the gigantocellular reticular nucleus shape locomotor pattern and rhythm in the freely behaving mouse. PLoS Biol 17, e2003880 (2019).

20. H. Liang, C. Watson, G. Paxinos, Terminations of reticulospinal fibers originating from the gigantocellular reticular formation in the mouse spinal cord. Brain Struct Funct 221, 1623–1633 (2016).

21. J. W. Chopek, Y. Zhang, R. M. Brownstone, Intrinsic brainstem circuits comprised of Chx10-expressing neurons contribute to reticulospinal output in mice. J Neurophysiol 126, 1978–1990 (2021).

22. M. Fitzgerald, The development of nociceptive circuits. Nat Rev Neurosci 6, 507–520 (2005).

23. G. F. Martin, R. P. Vertes, R. Waltzer, Spinal projections of the gigantocellular reticular formation in the rat. Evidence for projections from different areas to laminae I and II and lamina IX. Exp Brain Res 58, 154–162 (1985).

24. A. R. Wilson-Poe, E. Pocius, M. Herschbach, M. M. Morgan, The periaqueductal gray contributes to bidirectional enhancement of antinociception between morphine and cannabinoids. Pharmacol Biochem Behav 103, 444–449 (2013).

25. C. Watson, G. Paxinos, L. Puelles, The mouse nervous system (Elsevier Academic Press, Amsterdam ; Boston, ed. 1st, 2012), pp. xvii, 795 p.

26. B. I. Giasson et al., Neuronal alpha-synucleinopathy with severe movement disorder in mice expressing A53T human alpha-synuclein. Neuron 34, 521–533 (2002).

27. N. Ferreira et al., Trans-synaptic spreading of alpha-synuclein pathology through sensory afferents leads to sensory nerve degeneration and neuropathic pain. Acta Neuropathol Commun 9, 31 (2021).

28. M. B. Thomsen et al., PET imaging reveals early and progressive dopaminergic deficits after intra-striatal injection of preformed alpha-synuclein fibrils in rats. Neurobiol Dis 149, 105229 (2021).

29. L. Piilgaard et al., Non-invasive detection of narcolepsy type I phenotypical features and disease progression by continuous home-cage monitoring of activity in two mouse models: the HCRT-KO and DTA model. Sleep 46 (2023).

30. T. N. Luong, H. J. Carlisle, A. Southwell, P. H. Patterson, Assessment of motor balance and coordination in mice using the balance beam. J Vis Exp 10.3791/2376 (2011).

31. K. E. Glajch, S. M. Fleming, D. J. Surmeier, P. Osten, Sensorimotor assessment of the unilateral 6-hydroxydopamine mouse model of Parkinson’s disease. Behav Brain Res 230, 309–316 (2012).

32. R. J. Carter et al., Characterization of progressive motor deficits in mice transgenic for the human Huntington’s disease mutation. J Neurosci 19, 3248–3257 (1999).

33. N. Ferreira et al., Prodromal neuroinvasion of pathological alpha-synuclein in brainstem reticular nuclei and white matter lesions in a model of alpha-synucleinopathy. Brain Commun 3, fcab104 (2021).

34. J. Yerger et al., Phenotype assessment for neurodegenerative murine models with ataxia and application to Niemann-Pick disease, type C1. Biol Open 11 (2022).

35. P. V. Turner, D. S. Pang, J. L. Lofgren, A Review of Pain Assessment Methods in Laboratory Rodents. Comp Med 69, 451–467 (2019).

36. M. Richner, O. J. Bjerrum, A. Nykjaer, C. B. Vaegter, The spared nerve injury (SNI) model of induced mechanical allodynia in mice. J Vis Exp 10.3791/3092 (2011).

37. B. L. van der Gaag et al., Distinct tau and alpha-synuclein molecular signatures in Alzheimer’s disease with and without Lewy bodies and Parkinson’s disease with dementia. Acta Neuropathol 147, 14 (2024).

38. P. Bankhead et al., QuPath: Open source software for digital pathology image analysis. Sci Rep 7, 16878 (2017).

39. C. Stringer, T. Wang, M. Michaelos, M. Pachitariu, Cellpose: a generalist algorithm for cellular segmentation. Nat Methods 18, 100–106 (2021).

40. K. B. J. Franklin, G. Paxinos, Paxinos and Franklin’s The mouse brain in stereotaxic coordinates (Academic Press, an imprint of Elsevier, Amsterdam, ed. Fourth edition., 2013), pp. 1 volume (unpaged).

41. J. P. Anderson et al., Phosphorylation of Ser-129 is the dominant pathological modification of alpha-synuclein in familial and sporadic Lewy body disease. J Biol Chem 281, 29739–29752 (2006).

42. M. G. Spillantini et al., Alpha-synuclein in Lewy bodies. Nature 388, 839–840 (1997).

43. J. I. Ayers et al., Localized Induction of Wild-Type and Mutant Alpha-Synuclein Aggregation Reveals Propagation along Neuroanatomical Tracts. J Virol 92 (2018).

44. W. T. Chu et al., alpha-Synuclein Induces Progressive Changes in Brain Microstructure and Sensory-Evoked Brain Function That Precedes Locomotor Decline. J Neurosci 40, 6649–6659 (2020).

45. H. K. Chung, H. A. Ho, D. Perez-Acuna, S. J. Lee, Modeling alpha-Synuclein Propagation with Preformed Fibril Injections. J Mov Disord 12, 139–151 (2019).

46. K. C. Luk et al., Intracerebral inoculation of pathological alpha-synuclein initiates a rapidly progressive neurodegenerative alpha-synucleinopathy in mice. J Exp Med 209, 975–986 (2012).

47. A. N. Sacino et al., Intramuscular injection of alpha-synuclein induces CNS alpha-synuclein pathology and a rapid-onset motor phenotype in transgenic mice. Proc Natl Acad Sci U S A 111, 10732–10737 (2014).

48. T. Uchihara, B. I. Giasson, Propagation of alpha-synuclein pathology: hypotheses, discoveries, and yet unresolved questions from experimental and human brain studies. Acta Neuropathol 131, 49–73 (2016).

49. Z. A. Sorrentino et al., Motor neuron loss and neuroinflammation in a model of alpha-synuclein-induced neurodegeneration. Neurobiol Dis 120, 98–106 (2018).

50. E. A. Konnova, M. Swanberg, “Animal Models of Parkinson’s Disease” in Parkinson’s Disease: Pathogenesis and Clinical Aspects, T. B. Stoker, J. C. Greenland, Eds. (Brisbane (AU), 2018), 10.15586/codonpublications.parkinsonsdisease.2018.ch5.

51. A. Aniszewska, J. Bergstrom, M. Ingelsson, S. Ekmark-Lewen, Modeling Parkinson’s disease-related symptoms in alpha-synuclein overexpressing mice. Brain Behav 12, e2628 (2022).

52. M. Delenclos et al., Neonatal AAV delivery of alpha-synuclein induces pathology in the adult mouse brain. Acta Neuropathol Commun 5, 51 (2017).

53. R. H. Earls et al., NK cells clear alpha-synuclein and the depletion of NK cells exacerbates synuclein pathology in a mouse model of alpha-synucleinopathy. Proc Natl Acad Sci U S A 117, 1762–1771 (2020).

54. E. Gallagher et al., Positron Emission Tomography with [(18)F]ROStrace Reveals Progressive Elevations in Oxidative Stress in a Mouse Model of Alpha-Synucleinopathy. Int J Mol Sci 25 (2024).

55. R. Lalonde, C. Strazielle, Brain regions and genes affecting limb-clasping responses. Brain Res Rev 67, 252–259 (2011).

56. H. McCann, C. H. Stevens, H. Cartwright, G. M. Halliday, alpha-Synucleinopathy phenotypes. Parkinsonism Relat Disord 20 Suppl 1, S62–67 (2014).

57. H. C. Cheng, C. M. Ulane, R. E. Burke, Clinical progression in Parkinson disease and the neurobiology of axons. Ann Neurol 67, 715–725 (2010).

58. J. H. Kordower et al., Disease duration and the integrity of the nigrostriatal system in Parkinson’s disease. Brain 136, 2419–2431 (2013).

59. K. F. Farrell et al., Non-motor parkinsonian pathology in aging A53T alpha-synuclein mice is associated with progressive synucleinopathy and altered enzymatic function. J Neurochem 128, 536–546 (2014).

60. J. I. Ayers et al., Robust Central Nervous System Pathology in Transgenic Mice following Peripheral Injection of alpha-Synuclein Fibrils. J Virol 91 (2017).

61. C. M. Backman et al., Characterization of a mouse strain expressing Cre recombinase from the 3’ untranslated region of the dopamine transporter locus. Genesis 44, 383–390 (2006).

62. E. Rockenstein et al., Differential neuropathological alterations in transgenic mice expressing alpha-synuclein from the platelet-derived growth factor and Thy-1 promoters. J Neurosci Res 68, 568–578 (2002).

63. K. L. Emmer, E. A. Waxman, J. P. Covy, B. I. Giasson, E46K human alpha-synuclein transgenic mice develop Lewy-like and tau pathology associated with age-dependent, detrimental motor impairment. J Biol Chem 286, 35104–35118 (2011).

64. T. Kudo, D. H. Loh, D. Truong, Y. Wu, C. S. Colwell, Circadian dysfunction in a mouse model of Parkinson’s disease. Exp Neurol 232, 66–75 (2011).

65. S. M. Rothman et al., Neuronal expression of familial Parkinson’s disease A53T alpha-synuclein causes early motor impairment, reduced anxiety and potential sleep disturbances in mice. J Parkinsons Dis 3, 215–229 (2013).

66. I. Joniec-Maciejak et al., Effects of alpha-Synuclein Monomers Administration in the Gigantocellular Reticular Nucleus on Neurotransmission in Mouse Model. Neurochem Res 44, 968–977 (2019).

