## Supplementary material for "Bradykinesia and postural instability in a model of prodromal Synucleinopathy with α-Synuclein aggregation in the gigantocellular nuclei": Theologidis and Jan et al., Supplementary Information

- Table S1: Demographics, human cases used for IHC
- Table S2, Video clips of behaviors in the animal model with link to figshare repository
- Supplementary Figures Legends
- Supplementary Figures
- DVC analytics manual

**Table S1. Demographics of the cases examined for aSyn pathology in the GRN (medulla oblongata, p-aSyn S129)**

| ID | Clinical Diagnosis | Pathology | AD Braak Stage | PD Braak Stage | Cause of death | Gender | Age (yrs) | Remarks |
|---|---|---|---|---|---|---|---|---|
| Control-1 | Control | C | 2O | 0 | | Female | 83 | |
| Control-2 | Control | C | 1A | 0 | Myocard infarction | Female | 90 | |
| Control-3 | Control | C | 1A | 0 | Cachexia | Male | 79 | |
| Control-4 | Control | C | 3B | 0 | | Female | 88 | |
| PD-1 | PD | PSP | O | 5 | Cachexia and dehydration with Parkinson's disease | Female | 86 | PSP + LBD |
| PD-2 | PDD | PDD | 2O | 6 | Respiratory insufficiency and dehydration | Male | 74 | |
| PD-3 | C | iLBD | 1B | 4 | Respiratory insufficiency by pneumonia and exhaustion | Female | 87 | Non-demented with LBs |
| PD-4 | PD | PDD | 3B | 6 | Cachexia with end stage Parkinson's disease | Female | 90 | |

**Abbreviations:** C= Control; iLBD = Incidental Lewy body disease; LB = Lewy body pathology; LBD = Lewy body disease; PD = Parkinson disease; PDD = Parkinson's disease with dementia; PSP = Progressive supranuclear palsy.

**Table S2. Supplemental video files**

**figshare FOLDER NAME: TABLE S2- GRN MODEL VIDEOS**

| SUBFOLDER/TEST | Group | Days post-injection | File Name | Remark |
|---|---|---|---|---|
| OPEN FIELD | PBS | 120 | VIDEO 1 | $M83^{+/-}$ |
| | Monomeric aSyn | 120 | VIDEO 2 | $M83^{+/}$ |
| | PFF aSyn | 60 | VIDEO 3 | $M83^{+/}$ |
| | PFF aSyn | 120 | VIDEO 4 | $M83^{+/}$ |
| | PFF aSyn | 108* terminal | VIDEO 5 | $M83^{+/}$ |
| | PFF aSyn | 34 | VIDEO 18 | $M83^{+/+}$ |
| BALANCE BEAM | PBS | 120 | VIDEO 6 | $M83^{+/}$ |
| | Monomeric aSyn | 120 | VIDEO 7 | $M83^{+/}$ |
| | PFF aSyn | 60 | VIDEO 8 | $M83^{+/}$ |
| | PFF aSyn | 120 | VIDEO 9 | $M83^{+/}$ |
| | PFF aSyn | 27 | VIDEO 19 | $M83^{+/+}$ |
| POLE TEST | PBS | 120 | VIDEO 10 | $M83^{+/}$ |
| | Monomeric aSyn | 120 | VIDEO 11 | $M83^{+/}$ |
| | PFF aSyn | 60 | VIDEO 12 | $M83^{+/}$ |
| | PFF aSyn | 120 | VIDEO 13 | $M83^{+/}$ |
| HINDLIMB CLASPING | PBS | 120 | VIDEO 14 | $M83^{+/}$ |
| | Monomeric aSyn | 120 | VIDEO 15 | $M83^{+/}$ |
| | PFF aSyn | 60 | VIDEO 16 | $M83^{+/}$ |
| | PFF aSyn | 120 | VIDEO 17 | $M83^{+/}$ |
| | PFF aSyn | 27 | VIDEO 20 | $M83^{+/+}$ |

Link: https://doi.org/10.6084/m9.figshare.26936968

**Supplementary Figure Legends**

**S1. Spontaneous activity and locomotion in cohorts of heterozygous M83$^{+\backslash-}$ mice using non-invasive DVC monitoring. (A-D)** Non-invasive monitoring of spontaneous activity by cohorts of M83$^{+/-}$ mice (injected with PBS, monomeric aSyn or PFF aSyn) in digitally ventilated cages (DVC) over a longitudinal period up to 56 days post-injection (DPI) starting at DPI-36. The line chart (in A) represents the daily locomotion index (also presented as cumulative locomotion index in Fig. 3A), which is further represented for the light and dark periods for PBS (in B, n=5/2 cages), monomeric aSyn (in C, n=5/2 cages) or PFF aSyn- injected M83$^{+/-}$ mice (in D, n=12/4 cages). Also see Fig. 3B-C.

**S2. Measurements of locomotion behavior and motor strength in cohorts of heterozygous M83$^{+\backslash-}$ mice. (A-C)** Measurements of spontaneous activity by cohorts of M83$^{+/-}$ mice in the open-field arena recorded over 10 minutes (ANY-maze, see Material and Methods) at DPI-60 (in A) and DPI-90 (in B) and DPI-120 (in C); DPI, days post-injection. Bar graphs with individual points represent quantitative measurements of overall distance travelled, mean speed and time freezing as indicated. Also see Fig. 3D-G. **(D)** Assessment of motor strength (grip strength, front paws) of M83$^{+/-}$ cohorts over a longitudinal period (shown on x-axis as DPI, days post-injection). Statistics in S2A: Manny-Whitney test and in B-D: One-Way ANOVA followed by Tukey's multiple column comparisons (PBS, n=5, monomeric aSyn, n=5 and PFF aSyn, n=6-9; Error bars, Mean ± SD).

**S3. Measurements of gait stance during spontaneous locomotion in cohorts of heterozygous M83$^{+\backslash-}$ mice. (A)** Schematic depiction of the measures of gait stance, showing stride length, step width, rear base of support (RBOS), frontal base of support (FBOS) and step alternation, with rear paws painted in black and front paws painted in red (see Material and Methods). **(B)** Representative images showing gait coordination during spontaneous locomotion in PFF-injected M83$^{+/-}$ mice with scans from a male in the top panel and scans from a female in the bottom panel. Notice the pattern of coordinated gait at DPI-90 (exemplified by green arrows) and altered gait stance 1 day prior to euthanasia (indicated as pre-terminal), with a male (DPI-106) exhibiting abnormal placing of paws (instances exemplified by red arrows) and a female (DPI-121) starting to develop unilateral foot-drop (scale bar= 5 cm). **(C)** Bar graphs showing quantitative measurements of gait stance (indicated in S3A) of cohorts of M83$^{+/-}$ mice injected with PBS or PFF aSyn at DPI-45 and DPI-90 (DPI, days post-injection). Statistics in S3C: One-Way ANOVA followed by Dunn's multiple column comparisons (PBS, n=5 and PFF aSyn, n=10; Error bars, Mean ± SEM). No significant statistical differences were found in the measurements of gait stance. **(D)** Bar graphs showing quantitative measurements of the gait stance (indicated in S3A) of PFF aSyn-injected M83$^{+/-}$ mice at DPI-90 (DPI, days post-injection) or 1-2 days prior to euthanasia (termed pre-terminal stage). Statistics in S3D: Wilcoxon matched-pair signed rank test comparing mean measures (2-3 consecutive steps) for individual animals at DPI-90 and pre-terminal stage (Error bars, Mean ± SEM).

**S4. Neuroanatomical regions studied for the detection of phosphorylated alpha-synuclein (p-aSyn, S129) in the GRN-PFF aSyn mouse model. (A)** Representative images showing 2X low magnification annotated coronal views (scale bar= 1 mm). Bregma co-ordinates (in mm) are derived from Paxinos and Franklin's 4$^{th}$ edition with the following topographical landmarks: (*view in S4- Pons*) GRN/Gi- gigantocellular reticular nuclei, VN- vestibular nuclei; Rt-pontine intermediate reticular nuclei, DCN- deep cerebellar nuclei and *4V*- fourth ventricle; (*view in S4- Midbrain*) RN- red nucleus; PAG- periaqueductal grey, SN- substantia nigra, DG- dentate gyrus and *Aq*. cerebral aqueduct; (*view in S4- Thalamus*) DG- dentate gyrus, LH- lateral (and dorsomedial) hypothalamus, VPL/VPM, ventralposterior thalamus, STh- subthalamic nucleus, S1- somatosensory cortex and *LV*- lateral ventricle, and (*view in S4- Caudatoputamen*) Cpu-caudate putamen, M1/M2- primary motor cortex and *LV*- lateral ventricle. Also see Fig. 6A.

**S5. Dual immunofluorescence detection of phosphorylated alpha-synuclein (p-aSyn, S129) and neuronal nuclei (NeuN) in select brain regions of PFF aSyn-injected M83$^{+\backslash-}$ mice. (A-E)** Representative low magnification (10X) images showing p-aSyn (S129) and NeuN immunofluorescence detection in the brain regions of PFF aSyn-injected M83$^{+/-}$ mice at the highlighted time-points (DPI, days post-injection). Regions shown include: (*view in S4- Pons*) VN- vestibular nuclei (in A) and DCN- deep cerebellar nuclei (in B; *gr*. granule cell layer); (*view in S4- Midbrain*) RN- red nucleus (in C) and PAG- periaqueductal grey (in D; *Aq*. cerebral aqueduct), and (*view in S4- Thalamus*) STh- subthalamic nucleus. The insets show 63X magnified views of the selected region, highlighted by the white-bordered square in the 10X views (scale bar= 200 µm). Also see Fig. 5C (p-aSyn, S-129 in pontine GRN/Gi), Fig. 6A, Supplementary Figures S6-S7 (markers of inflammatory gliosis) and S8 (aSyn pathology in the terminal-stage M83$^{+/+}$ mice). Primary antibodies in Figure S5A-E: p-aSyn (S129)- abcam EP1536Y and NeuN- EMD Millipore A60.

**S6. Immunofluorescence detection of phosphorylated alpha-synuclein (p-aSyn, S129) and inflammatory gliosis in select brain regions of PFF aSyn-injected M83$^{+\backslash-}$ mice. (A)** Representative low magnification (10X) images showing p-aSyn (S129) and NeuN immunofluorescence detection in the brain regions of PFF aSyn-injected M83$^{+/-}$ mice at terminal stage DPI≥108 (DPI, days post-injection). Regions shown: (*view in S4- Caudatoputamen*) M1/M2- primary motor cortex and Cpu- striatum; (*view in S4- Thalamus*) VPL/VPM, ventralposterior thalamus and DM/LH- dorsomedial and lateral hypothalamus; (*view in S4- Midbrain*) SN- substantia nigra (*view in S4- Pons*) Rt- pontine intermediate reticular nuclei. The insets show 63X magnified views of select regions with sparse detection/negative for p-aSyn S129 (except pontine Rt), highlighted by the white-bordered square in the 10X views (scale bar= 200 µm). Also see Fig. 5C (p-aSyn, S-129 in pontine GRN/Gi), Fig. 6A, Supplementary Figures S6-S7 (markers of inflammatory gliosis) and S8 (aSyn pathology in the terminal-stage

M83$^{+/+}$ mice). Primary antibodies in Figure S6A: p-aSyn (S129)- abcam EP1536Y and NeuN- EMD Millipore A60. **(B-D)** Representative low magnification (10X) images showing dual immunofluorescence detection of p-aSyn (S129) and p62 (Sequestosome 1- a marker of protein inclusions, in B), glial fibrillary acidic protein (GFAP- astroglial marker, in C), CD68 (phagocyte/microglia marker, in D) in select brain regions of PFF aSyn-injected M83$^{+/-}$ mice at DPI≥108 (DPI, days post-injection). Regions shown: (*view in S4- Pons*) GRN/Gi- gigantocellular reticular nuclei; (*view in S4- Midbrain; Aq*. cerebral aqueduct) PAG- periaqueductal grey and (*view in S4- Caudatoputamen; LV.* Lateral ventricle) M1/M2- primary motor cortex. The insets in all panels show 63X magnified views of the select region, highlighted by the white-bordered square in the 10X views (scale bar= 200 µm). The inset in M1/M2 (S6B, GFAP) is included as an example of astroglial lining in the vicinity of lateral ventricle (*LV.*), i.e., positive detection by the antibody away from M1/M2. Primary antibodies in Figure S6: (S6A) p-aSyn (S129)- abcam EP1536Y, (S6B) p62- Nordic Biosite GP62-C, (S6C) GFAP- abcam ab4674, and (S6D) CD68- Novus Biologicals FA-11. Also see Fig. S7B-D.

**S7. Immunodetection of phosphorylated alpha-synuclein (p-aSyn, S129), neuronal loss and inflammatory gliosis in cohorts of M83$^{+\backslash -}$ mice. (A)** Quantitative estimates of p-aSyn S129 immunopositivity (immunofluorescence signal intensity from two serial sections) in the GRN/Gi in cohorts of M83$^{+/+}$ mice expressed as percentage of total p-aSyn S129 (Fig. 6B) detected in neuronal soma (overlapping with NeuN staining) or away from soma (no overlap with NeuN). Also see Fig. 5A-C. Statistics in S7A: Wilcoxon matched-pair signed rank test (PBS, n=5; monomeric aSyn, n=5; PFF aSyn DPI-30 and DPI-60, n=3 and PFF aSyn DPI≥108, n=6; *$p<0.05$; Error bars, Mean ± SEM). Black asterisks (on DPI≥108) represent significant differences between somatic or other p-aSyn S129 immunofluorescence. Only significant differences are highlighted. **(B)** Quantitative estimates of p-aSyn S129 immunopositivity (immunofluorescence signal from two serial sections) in select brain regions in cohorts of PFF aSyn-injected M83$^{+/+}$ mice, expressed as percentage of the area (total average area/image: 625,000 um$^2$) in 10X views (Also see Fig. 5A). Regions included in S7B: (GRN) gigantocellular nuclei, (VN) vestibular nuclei, (DCN) deep cerebellar nuclei. (RN) red nucleus, (PAG) periaqueductal grey matter, (STh) subthalamic nucleus and (M1/M2) motor cortex. Also see Supplementary Figure S4 for the low magnification (2X) panoramic views annotating select brain regions. Statistics in S7B: Two-Way ANOVA followed by Tukey's multiple column comparisons: DPI-30 (n=3), DPI-60 (n=3) and DPI≥108, n=6; *p<0.05, **p<0.01, ***p<0.001, ****p<0.0001; Error bars, Mean ± SEM). Grey asterisks represent significant differences between DPI-30 and DPI≥108, while the black asterisks significant differences between DPI-60 and DPI≥108 (ns= not significant). Also see Fig. 6A. **(C-D)** Quantitative estimates of astrogliosis (GFAP, in C) and microglial infiltration (CD68, in D) in 10X views of GRN/Gi from cohorts of M83$^{+/-}$ mice (also see Fig. S6C-D).

**S8. Immunodetection of phosphorylated alpha-synuclein (p-aSyn, S129) in the brains of PFF aSyn-injected homozygous M83$^{+\backslash+}$ and heterozygous M83$^{+\backslash-}$ mice. (A-B)** Representative low magnification (2X) images showing p-aSyn (S129) detection in pons of PFF aSyn-injected homozygous M83$^{+/+}$ mice at terminal stage (DPI≥34), with chromogenic method (DAB, in A) and immunofluorescence (in B, dual detection with NeuN). The insets show 20X magnified views of the GRN/Gi (scale bar= 1 mm). **(C-E)** Representative low magnification (10X) immunofluorescence images showing dual detection of p-aSyn (S129) and NeuN in select brain regions of PFF aSyn-injected M83$^{+/+}$ mice at terminal stage (DPI≥34). Regions shown: (*view in S4- Pons*) VN- vestibular nuclei (in C), (*view in S4- Midbrain; Aq*. cerebral aqueduct) PAG- periaqueductal grey (in D) and (*view in S4- Caudatoputamen*) M1/M2- primary motor cortex (in E). The insets show 63X magnified views of the select region, highlighted by the white-bordered square in the 10X views (scale bar= 200 µm). Primary antibodies: p-aSyn (S129)- abcam EP1536Y (all panels) and NeuN- EMD Millipore A60 (S8B-E). **(F)**

Representative low magnification (10X) images showing p-aSyn (S129) immunostaining by EP1536Y antibody in GRN/Gi of PFF aSyn-injected heterozygous $M83^{+/-}$ mice at terminal stage DPI≥108 (DPI, days post-injection), with chromogenic method (DAB). The inset on top right show 63X magnified views of the GRN/Gi highlighted by the black-bordered square in the 10X views (scale bar= 200 µm). For comparison, an example of 63X magnified view of the GRN/Gi from a control (PBS-injected, DPI-120) animal is shown in the inset on bottom left. **(G)** Representative low magnification (10X) images showing p-aSyn (S129) detection by D1R1R antibody in GRN/Gi of PFF aSyn-injected heterozygous $M83^{+/-}$ mice at terminal stage DPI≥108 (DPI, days post-injection), with chromogenic method (DAB, on left) and dual immunofluorescence detection (on right). The insets show 63X magnified views highlighted by the squares in the 10X views (scale bar= 200 µm).

#### A) Daily Locomotion Index- all M83$^{+/-}$ cohorts

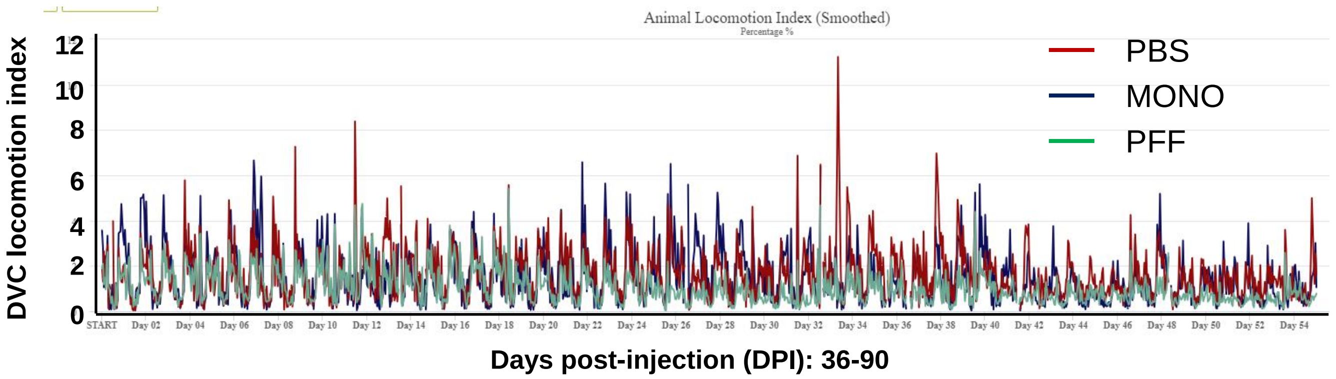

#### B) Daily Rhythym- PBS, M83$^{+/-}$ (n=5/2 cages)

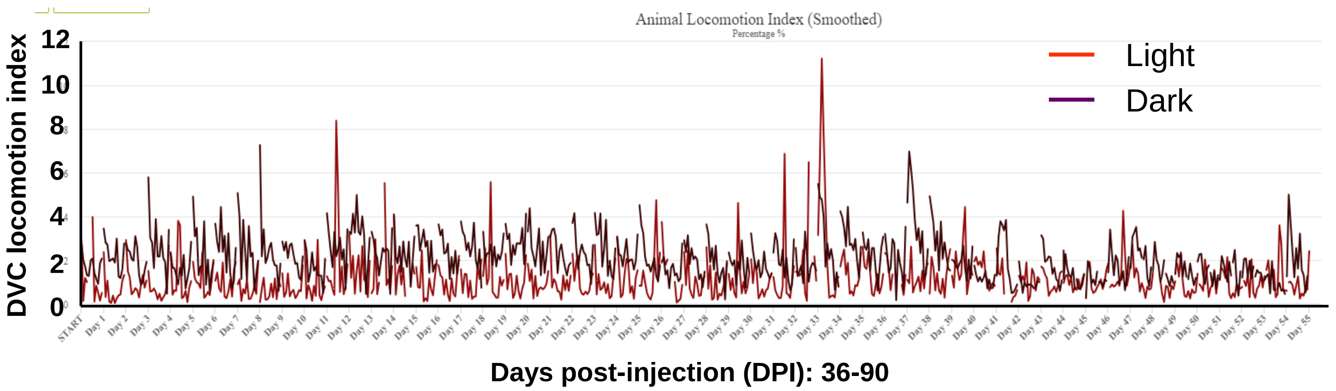

#### C) Daily Rhythym- Monomeric aSyn, M83$^{+/-}$ (n=5/2 cages)

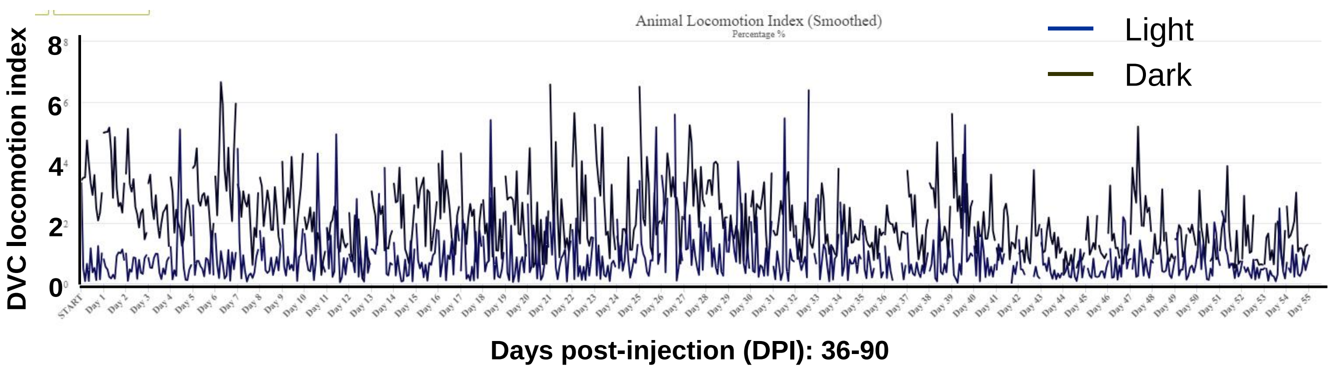

#### D) Daily Rhythym- PFF aSyn, M83$^{+/-}$ (n=12/4 cages)

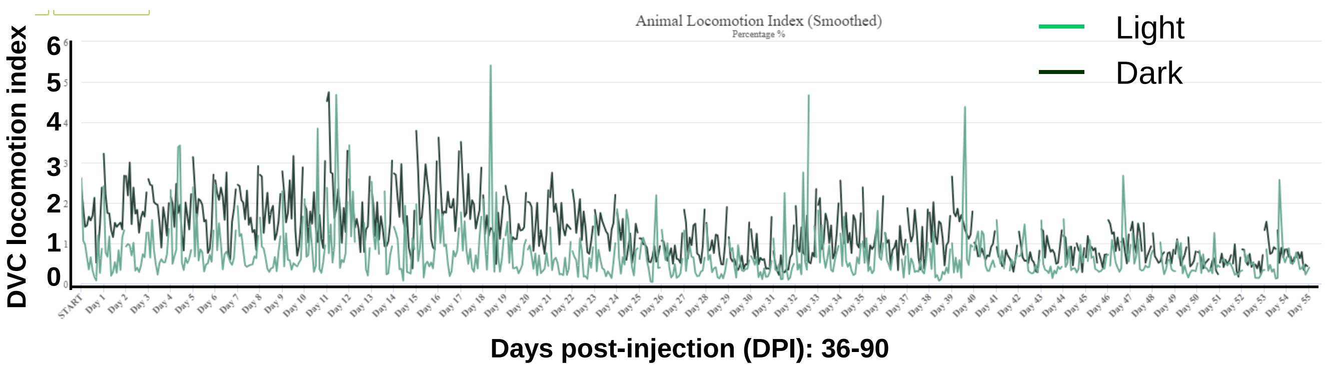

S2

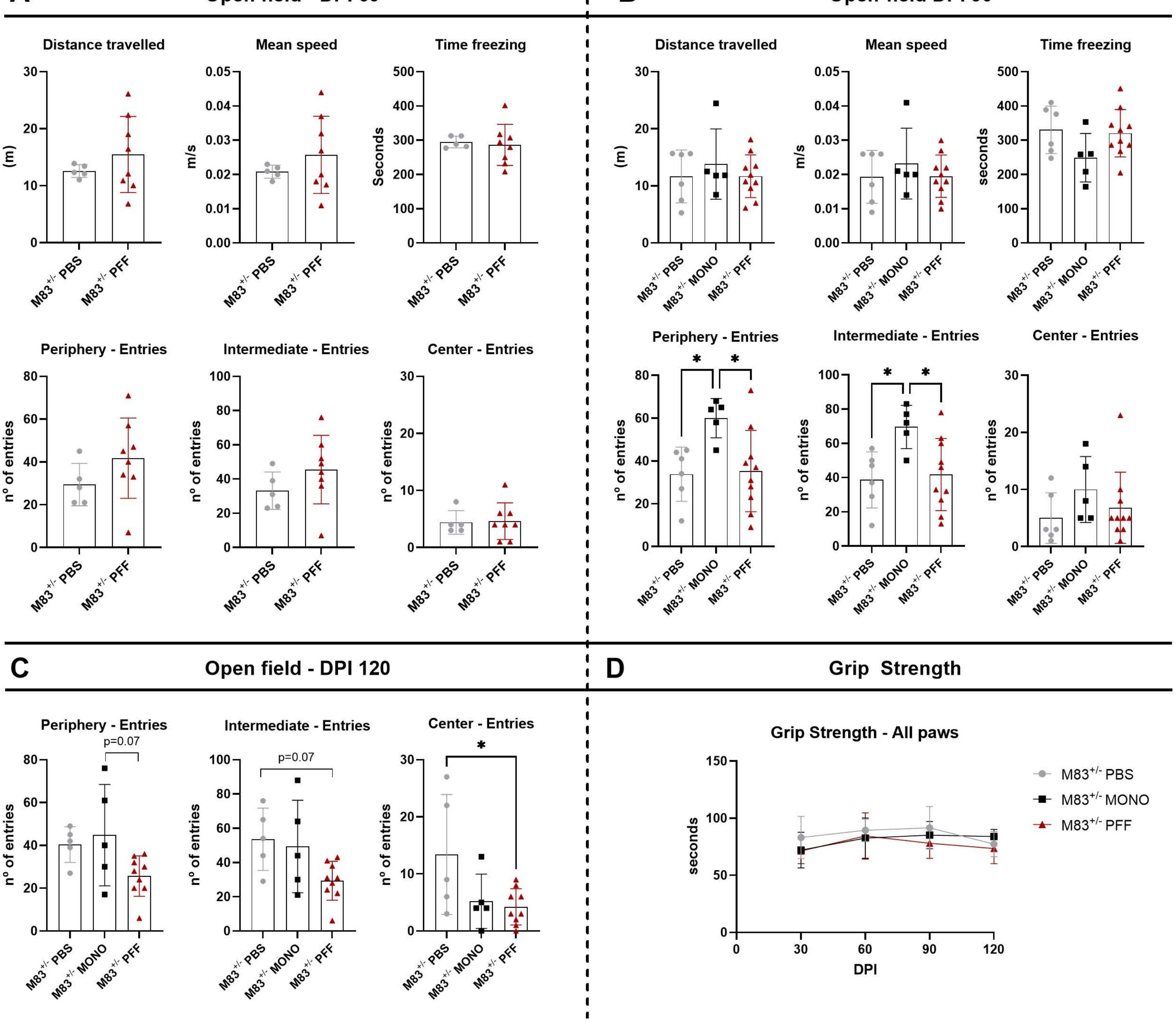

A
Open field - DPI 60
Distance travelled
(m)
Mean speed
m/s
Time freezing
Seconds
Periphery - Entries
Intermediate - Entries
Center - Entries
nº of entries
M83+/- PBS
M83+/- PFF
B
Open-field DPI 90
Distance travelled
(m)
Mean speed
m/s
Time freezing
seconds
Periphery - Entries
Intermediate - Entries
Center - Entries
nº of entries
M83+/- PBS
M83+/- MONO
M83+/- PFF
C
Open field - DPI 120
Periphery - Entries
p=0.07
Intermediate - Entries
p=0.07
Center - Entries
nº of entries
M83+/- PBS
M83+/- MONO
M83+/- PFF
D
Grip Strength
Grip Strength - All paws
seconds
DPI
M83+/- PBS
M83+/- MONO
M83+/- PFF

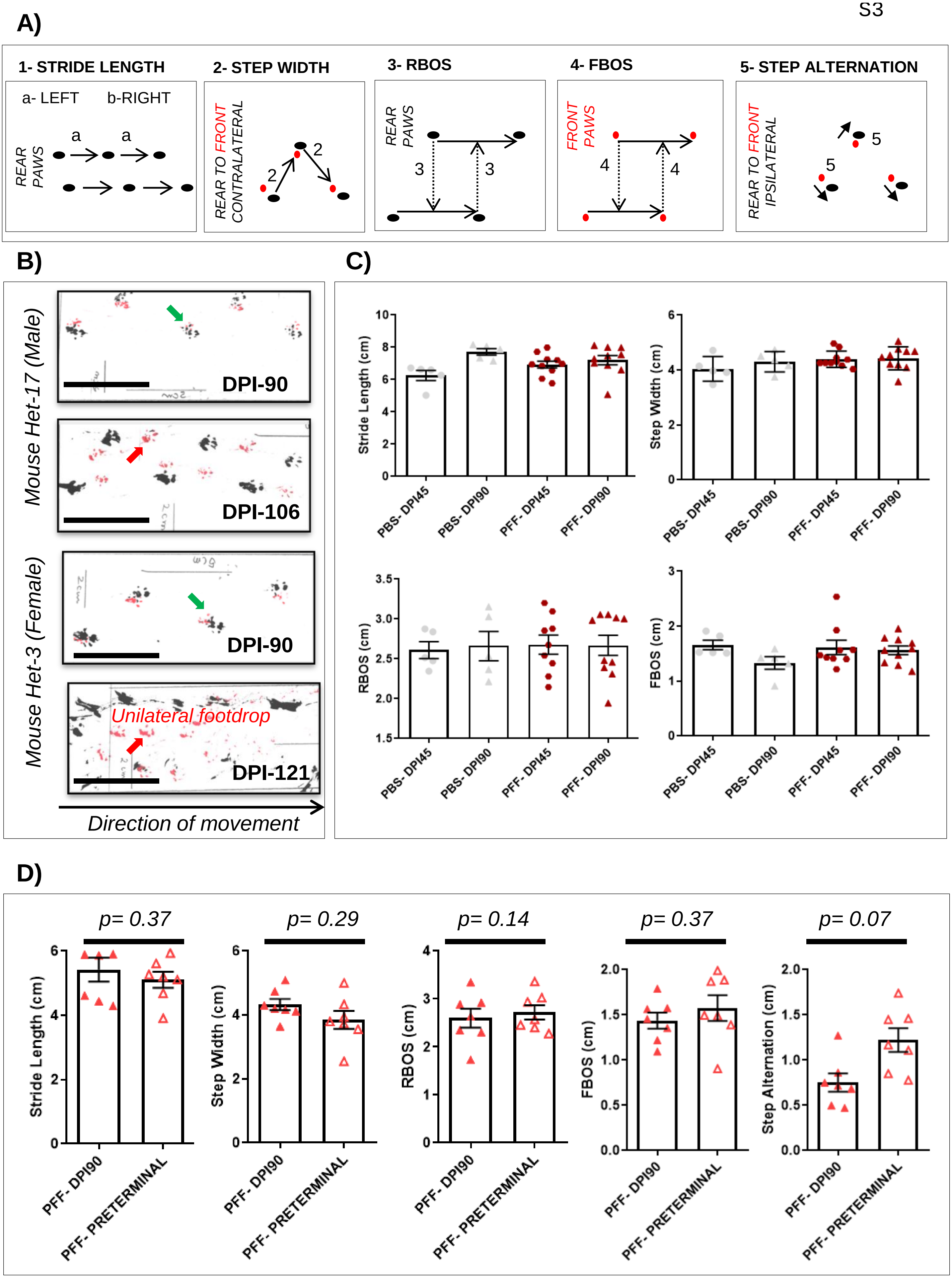
A)
1- STRIDE LENGTH
a- LEFT
b-RIGHT
REAR PAWS
2- STEP WIDTH
REAR TO FRONT CONTRALATERAL
3- RBOS
REAR PAWS
4- FBOS
FRONT PAWS
5- STEP ALTERNATION
REAR TO FRONT IPSILATERAL
B)
Mouse Het-17 (Male)
DPI-90
DPI-106
Mouse Het-3 (Female)
DPI-90
Unilateral footdrop
DPI-121
Direction of movement
C)
Stride Length (cm)
Step Width (cm)
RBOS (cm)
FBOS (cm)
PBS- DPI45
PBS- DPI90
PFF- DPI45
PFF- DPI90
D)
p= 0.37
p= 0.29
p= 0.14
p= 0.37
p= 0.07
Stride Length (cm)
Step Width (cm)
RBOS (cm)
FBOS (cm)
Step Alternation (cm)
PFF- DPI90
PFF- PRETERMINAL

A)

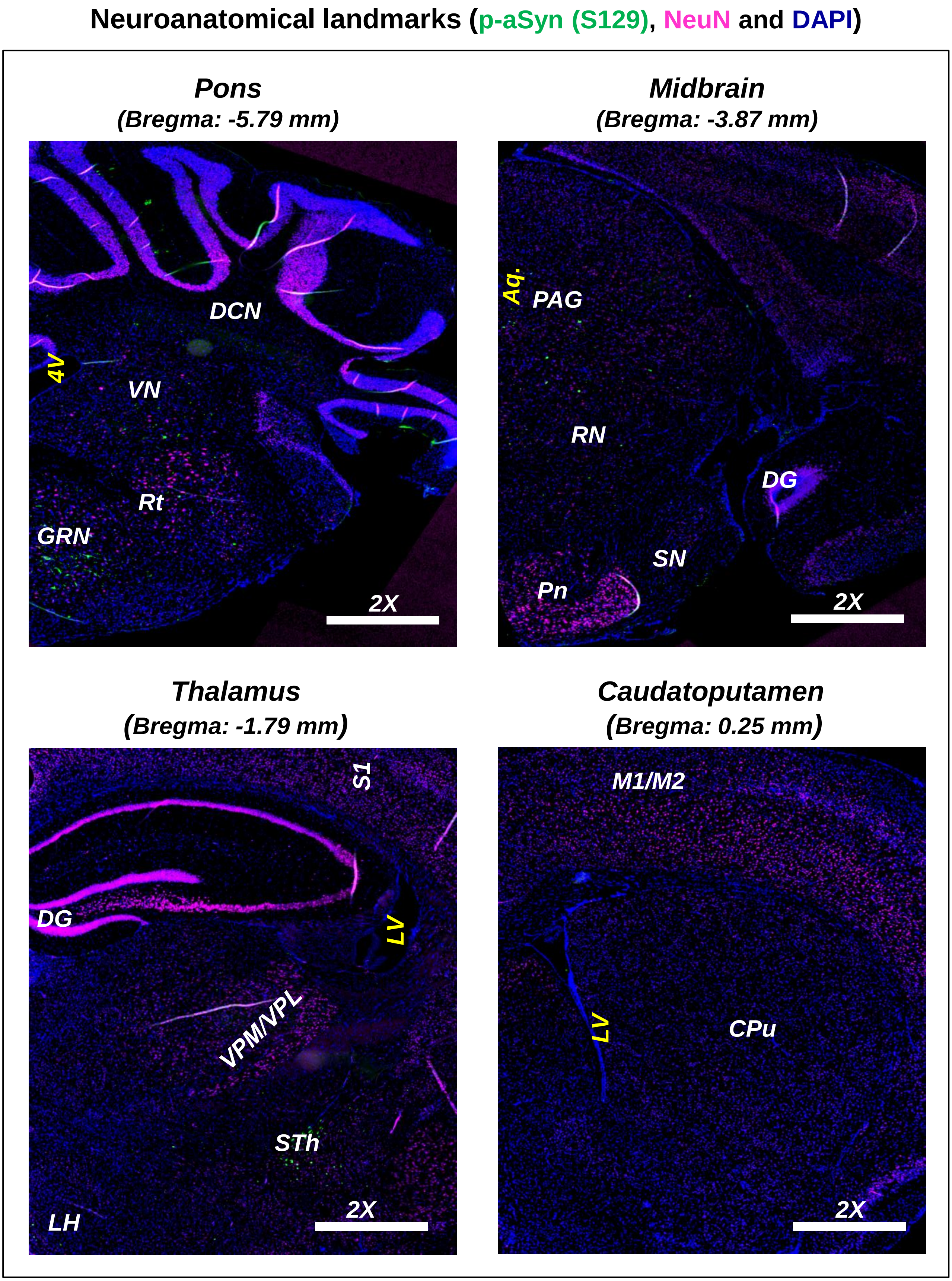
Neuroanatomical landmarks (p-aSyn (S129), NeuN and DAPI)
Pons
(Bregma: -5.79 mm)
DCN
4V
VN
Rt
GRN
2X
Midbrain
(Bregma: -3.87 mm)
Aq.
PAG
RN
DG
SN
Pn
2X
Thalamus
(Bregma: -1.79 mm)
S1
DG
LV
VPM/VPL
STh
LH
2X
Caudatoputamen
(Bregma: 0.25 mm)
M1/M2
LV
CPu
2X

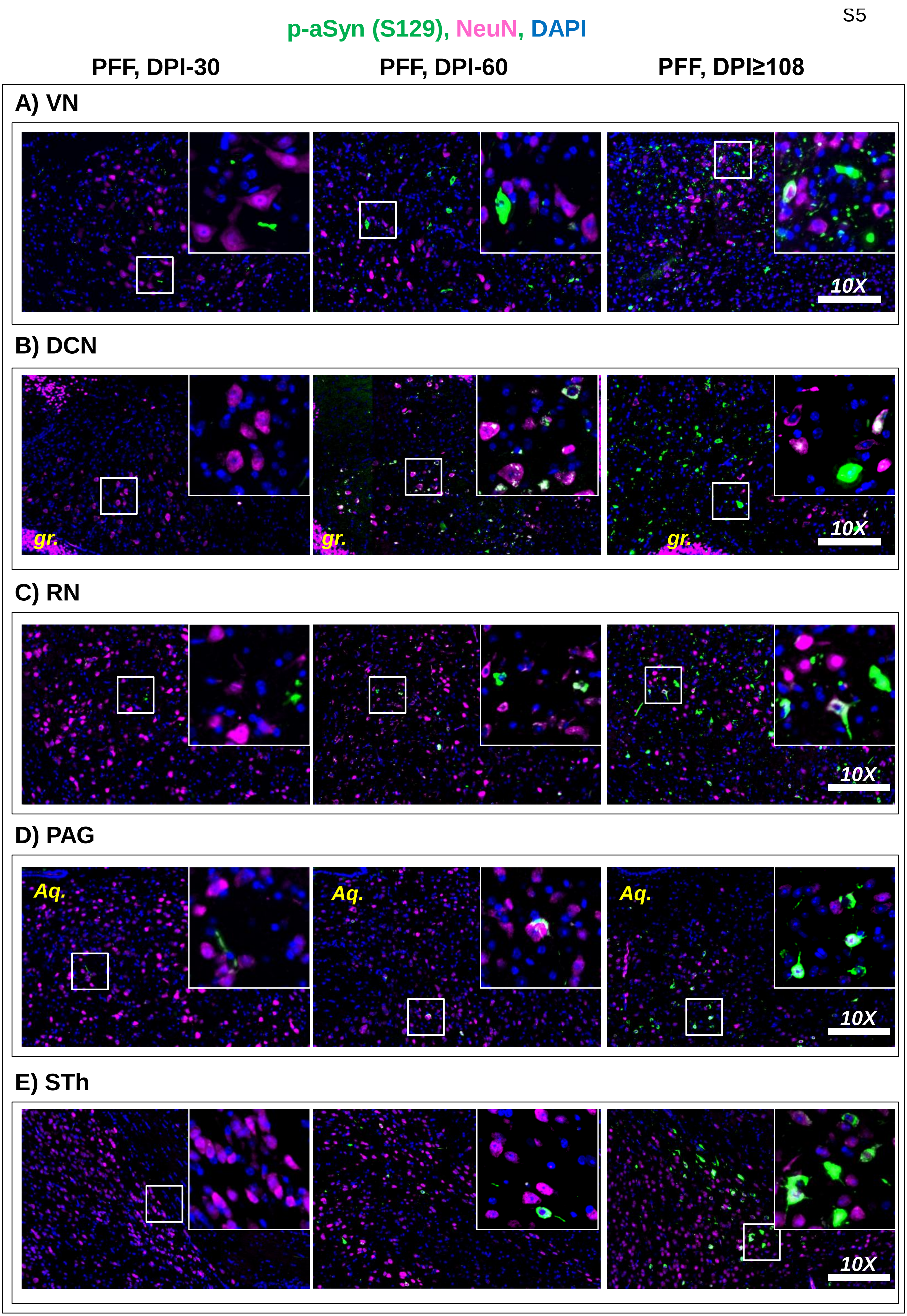
p-aSyn (S129), NeuN, DAPI
PFF, DPI-30
PFF, DPI-60
PFF, DPI≥108
A) VN
10X
B) DCN
gr.
gr.
gr.
10X
C) RN
10X
D) PAG
Aq.
Aq.
Aq.
10X
E) STh
10X

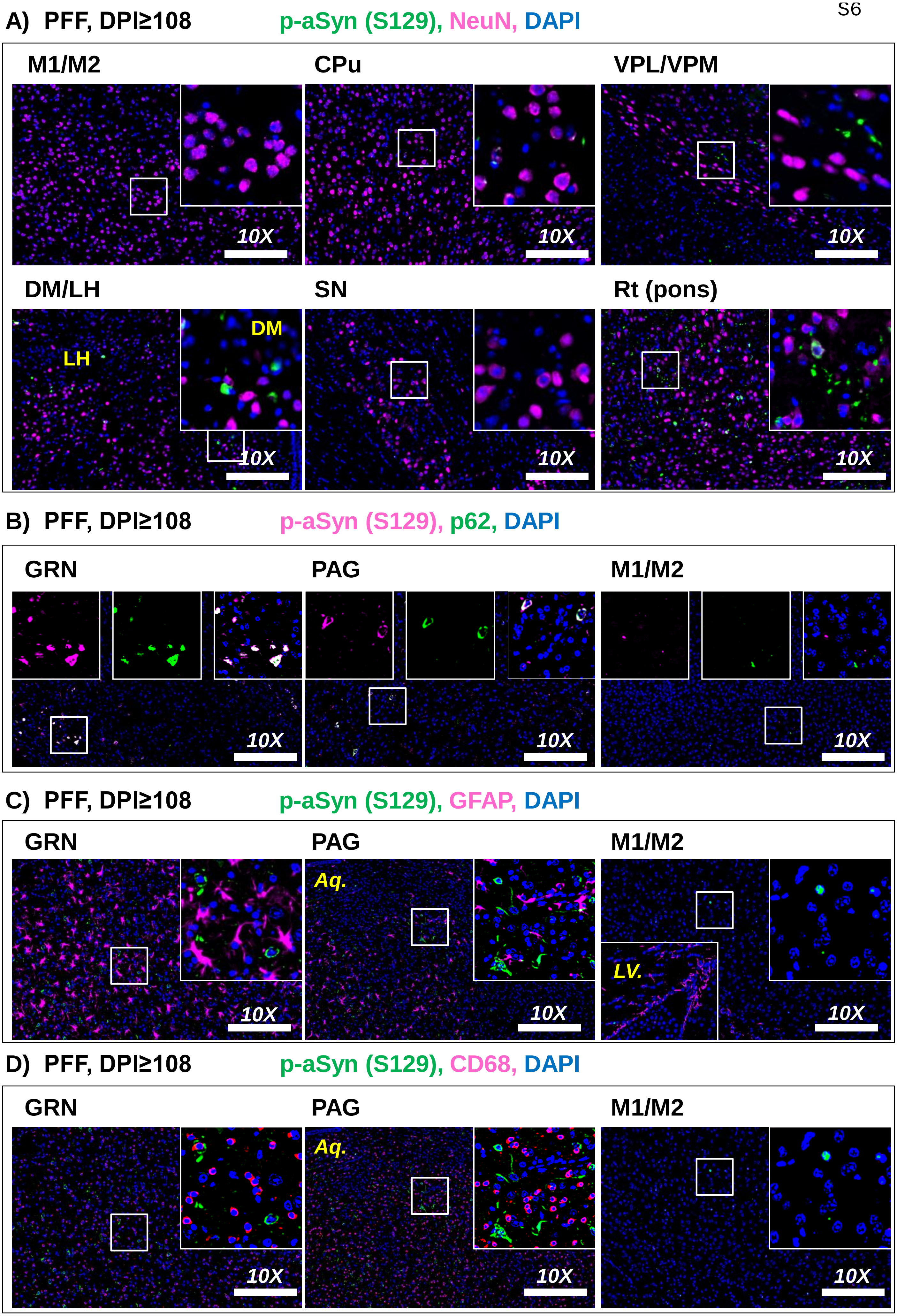
A) PFF, DPI≥108
p-aSyn (S129), NeuN, DAPI
M1/M2
CPu
VPL/VPM
DM/LH
LH
DM
SN
Rt (pons)
10X
B) PFF, DPI≥108
p-aSyn (S129), p62, DAPI
GRN
PAG
M1/M2
10X
C) PFF, DPI≥108
p-aSyn (S129), GFAP, DAPI
GRN
PAG
Aq.
M1/M2
LV.
10X
D) PFF, DPI≥108
p-aSyn (S129), CD68, DAPI
GRN
PAG
Aq.
M1/M2
10X

A) p-aSyn, S129 (GRN)

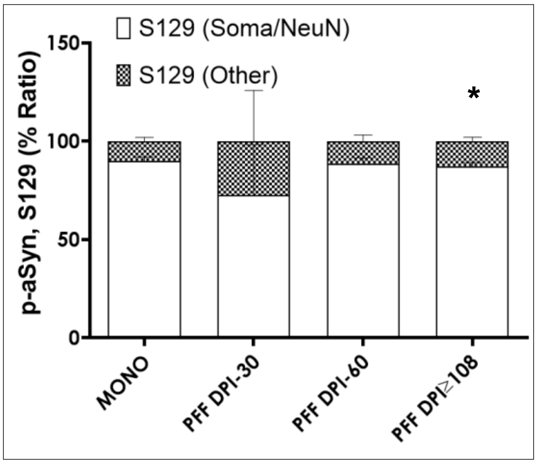

B) p-aSyn, S129 (regions)

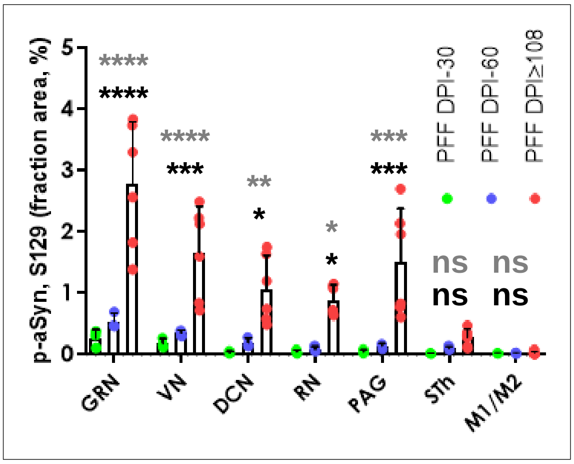

C) GFAP (GRN)

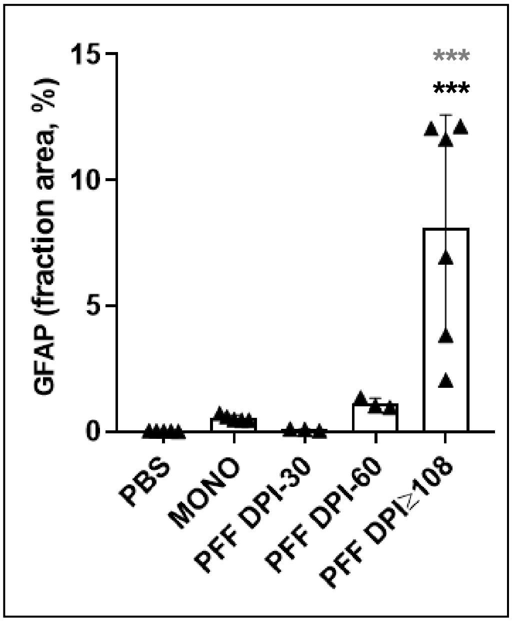

D) CD68 (GRN)

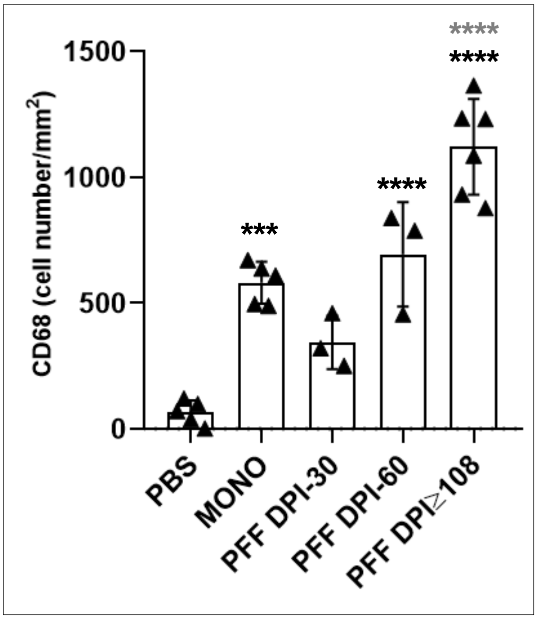

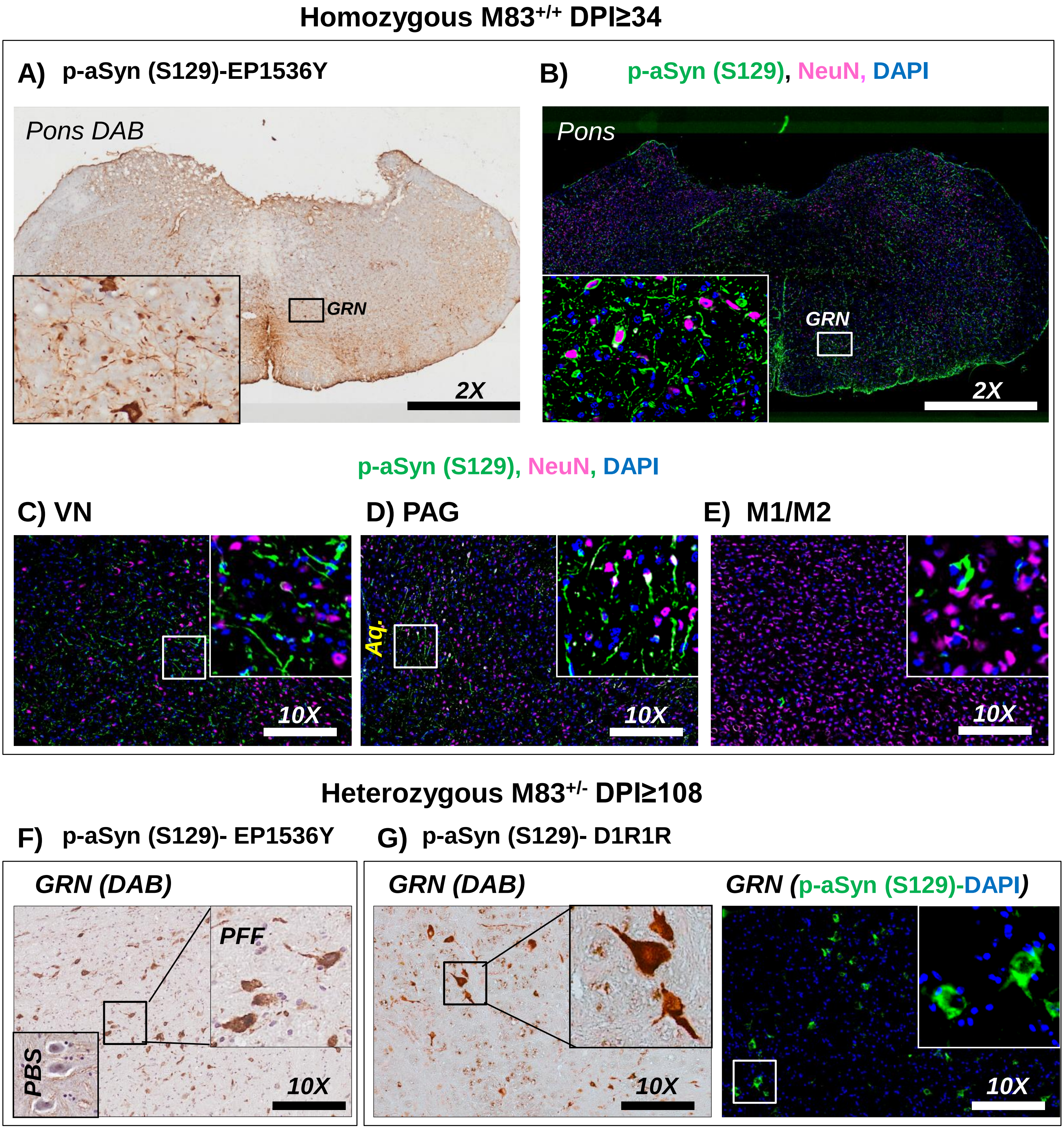
Homozygous M83+/+ DPI≥34
A) p-aSyn (S129)-EP1536Y
Pons DAB
GRN
2X
B) p-aSyn (S129), NeuN, DAPI
Pons
GRN
2X
p-aSyn (S129), NeuN, DAPI
C) VN
10X
D) PAG
Aq.
10X
E) M1/M2
10X
Heterozygous M83+/- DPI≥108
F) p-aSyn (S129)- EP1536Y
GRN (DAB)
PFF
PBS
10X
G) p-aSyn (S129)- D1R1R
GRN (DAB)
10X
GRN (p-aSyn (S129)-DAPI)
10X

### DVC® ANALYTICS INSTRUCTION MANUAL

#### Summary

**Revision History**

| Date | Version | Author | Summary of Changes |
|---|---|---|---|
| 15/05/17 | V1.0 | Giorgio Rosati | First Draft |
| 16/05/17 | V1.1 | Giorgio Rosati | Explanation how to download raw data |
| 27/07/2018 | V2.0 | Giorgio Rosati | DVC Analytics version 2 |
| 19/09/2019 | V3.0 | Giorgio Rosati | DVC Analytics version 3 |

DVC® ANALYTICS INSTRUCTION MANUAL

#### 1 Login in page

In order to login into the system, go to: https://analytics.dvc.tecniplast.it/login/XXX (XXX provided to you by Tecniplast) and enter your username and password that have been previously communicated.

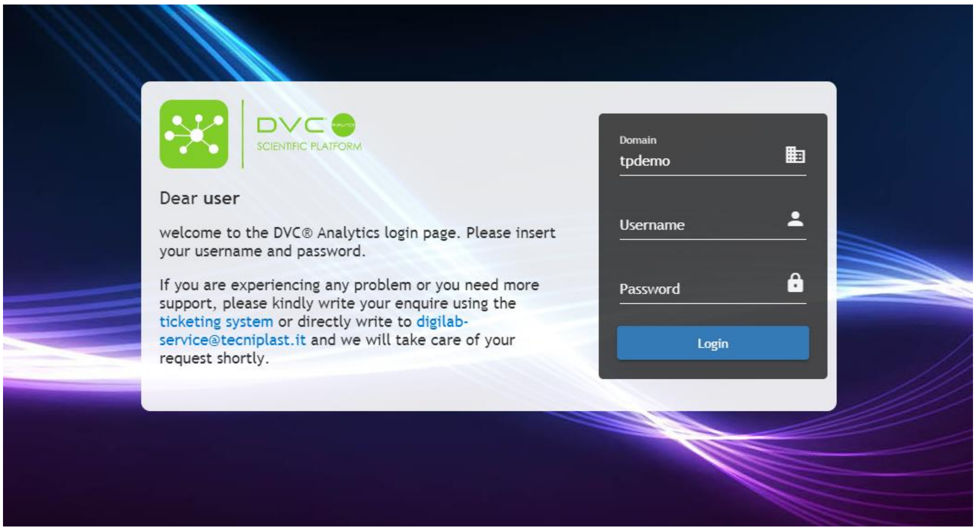

#### 2 Home Page

The top part of the home page (dark blue background) displays general information about the Facility (e.g. Facility name). The lower part of the home page (white background) displays the DVC® Analytics menu.

The home page varies based on the user role whether Facility Manager or Researcher (the User Setting button is hidden to the Researcher(s) that will have access only to the cages assigned previously by the Facility Manager)

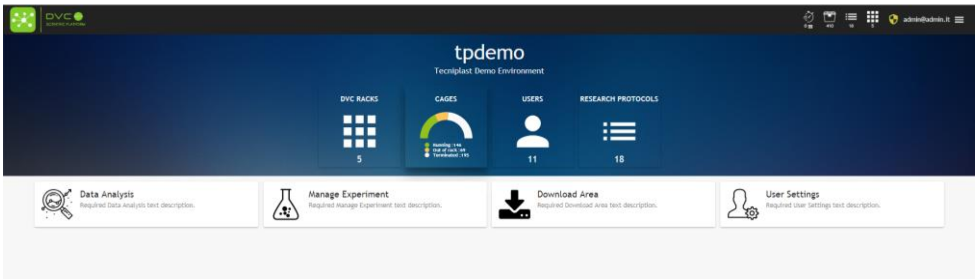

**DVC RACKS**: how many DVC® Racks currently connected to the DVC® Analytics platform

**CAGES**: how many cages:

- **Running (green):** # of registered cages in the DVC® system currently inserted in the DVC® Rack
- **Out of Rack: (yellow):** # of registered cages in the DVC® system currently removed from the DVC® Rack (but still registered)
- **Terminated (grey):** # of cages properly terminated in the DVC® system

**USERS:** how many registered DVC® Analytics Users in the specific Facility

**RESEARCH PROTOCOLS**: how many DVC® Research Protocols have been currently received by the DVC® Analytics (sent by the DVC® System)

In The lower part of the home page shows different available buttons dependently upon your role (Facility/Researcher):

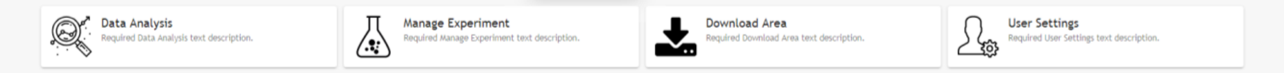

Through those buttons you can analyze data and set different Facility rules.

#### 2.1 Facility Manager and Researcher Roles

Facility Manager role has been designed in order to perform any task and have access to all functionalities of the DVC® Analytics system. Researcher role has been designed to provided limited access to only a subset of information (only to the Researcher's data) in order to increase privacy and limit any possibility of stolen data.

More specifically, these are the different enabled functionalities for the different roles:

| **FACILITY MANAGER**<br>(has full access to the DVC® Analytics menu) | **RESEARCHER**<br>(limited access to the DVC® Analytics menu) |
|---|---|
| • is enabled to view all DVC® cages connected to the DVC® Analytics.<br>• is enabled to register new DVC® Analytics users and associated them to DVC® cages.<br>• is enabled to group together DVC® users and associate them to the DVC® cages.<br>• is enabled to set and change the facility settings (e.g. dark hour). | • is enabled to view only DVC® cages that have been previously associated to her/him.<br>• Is enabled to create cage and mice groups using only his/her cages |

#### 2.2 Data Analysis

In this section it is possible to analyze in depth an individual cage or group of cages selecting across multiple choices from temporal and data presentation perspective. Just clicking on the "Data Analysis" icon, you can enter in a menu section where multiple choices are available:

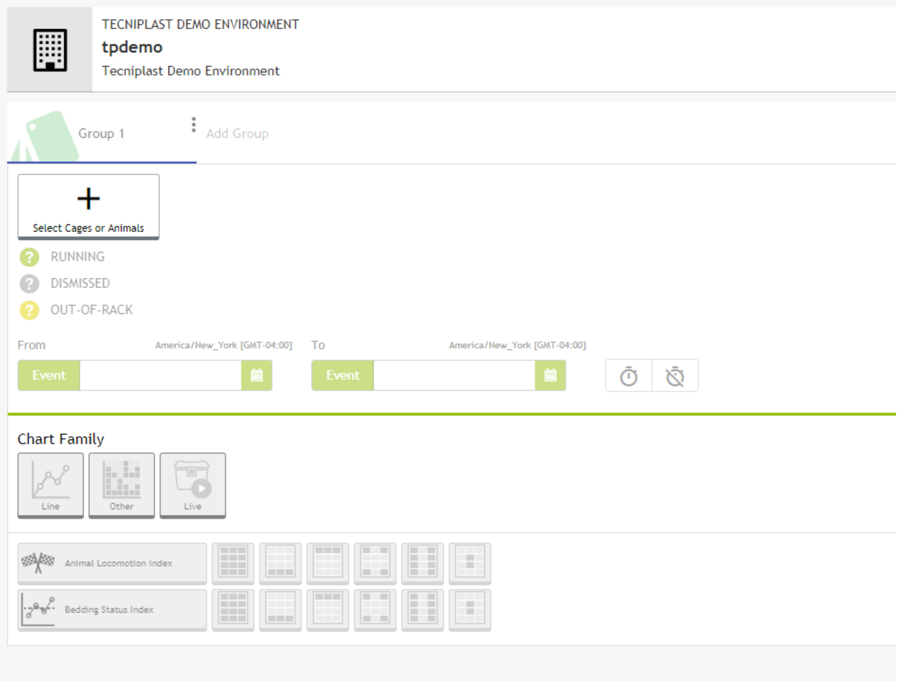

The first requested information is to select "Cages" or "Animals" by simply clicking on the corresponding button (all the other fields are disabled until this first selection is performed):

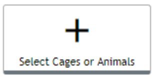

Once you click, a pop-up menu appears with multiple choices:

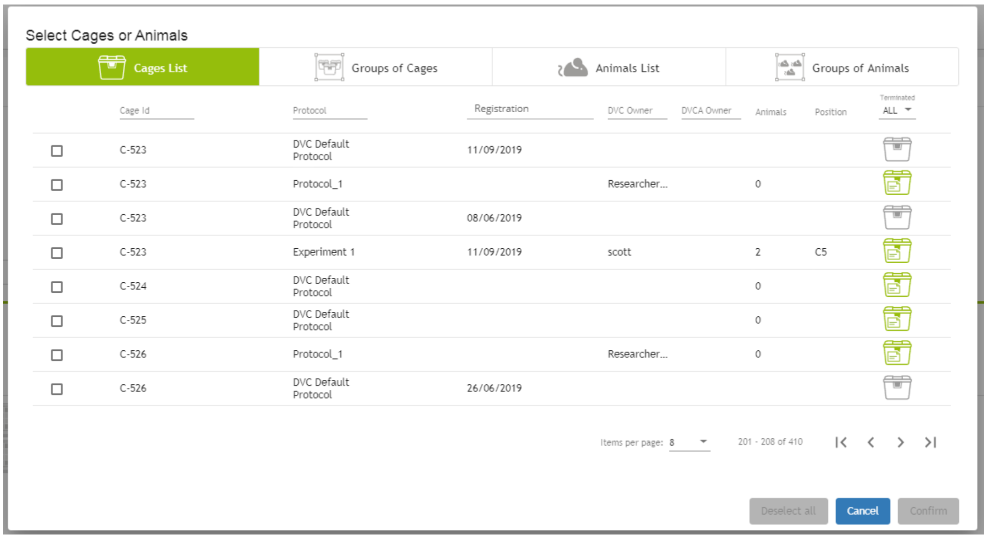

You can immediately select a cage (or multiple) searching for its Cage Id, you can filter by (research) Protocols (set in the DVC® system), time of Registration (when it has been registered in the DVC® system), DVC® Owner (set in the DVC® system). Moreover, there are other information like # animals in the cage ID and current rack position.

You can also select an already created “Group of Cages”

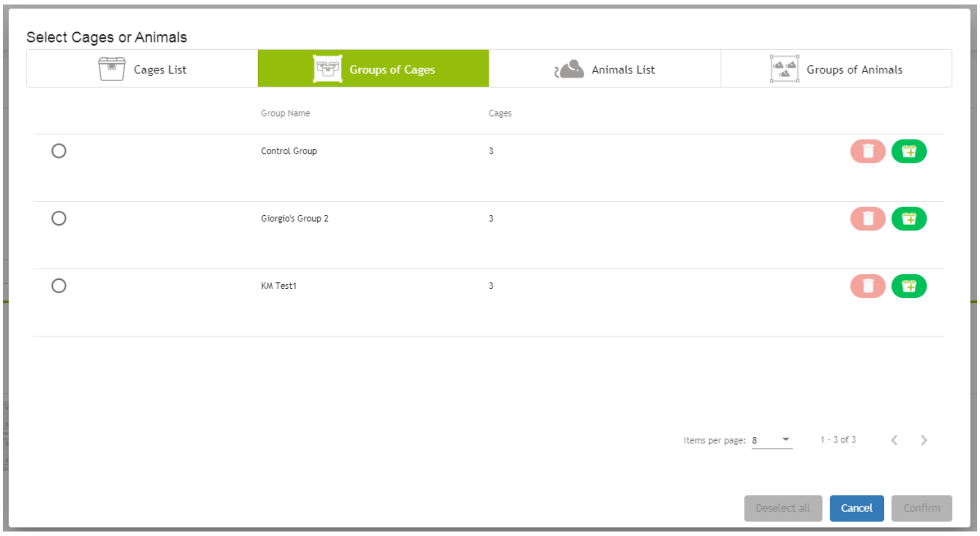

As well as you can select directly Animal IDs:

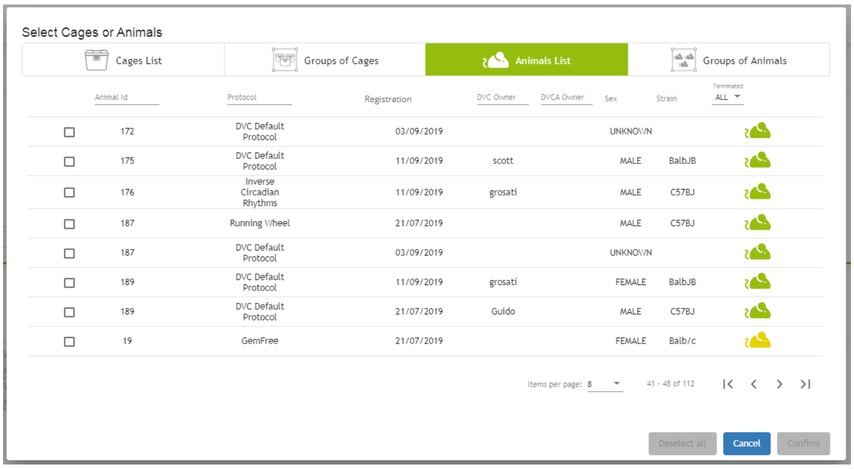

Or “Groups of Animals” if already created:

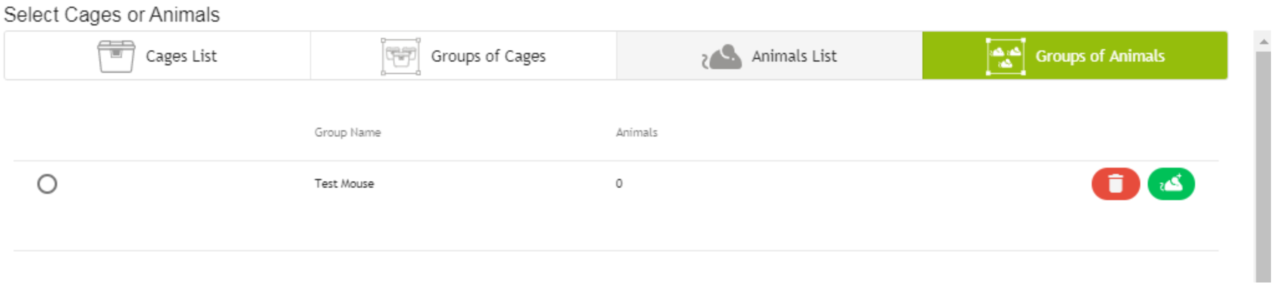

**PLEASE NOTE**: you can only select same type of elements (cages with cages or animals with animals, but not cages with animals at the same time).

Then, when at least one element (cage or animal ID) has been selected, other submenus become available.

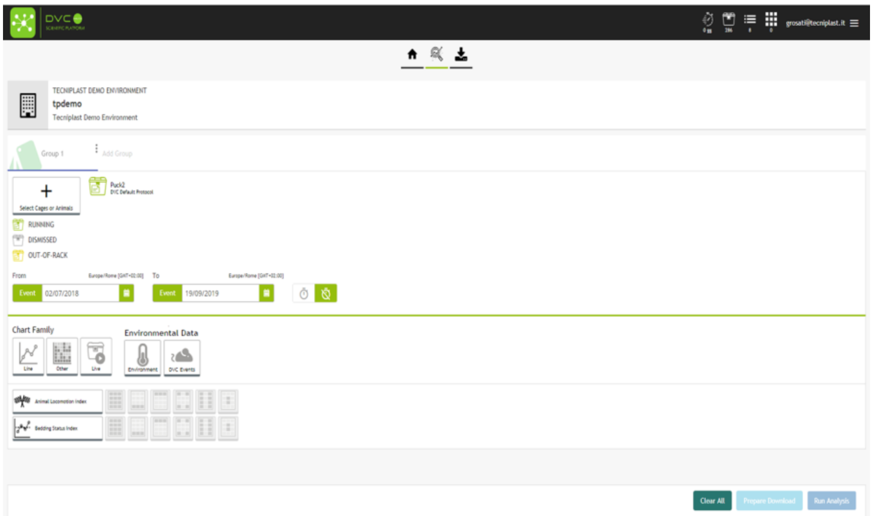

- **Group name and creation**: you can save this selection for further analysis.
- **From - To**: automatically filled with the date & Time of Registration and Termination (if any, otherwise the actual time if the cage(s) is still running).
- **Chart Family**: different available charts.

- **Environmental Data**: events generated in the DVC® system and environmental data (captured by REM if it is installed).
- **DVC® available metrics**: Animal Locomotion Index and Bedding Status Index

**PLEASE NOTE**: you cannot "Run Analysis" neither "Prepare Download" until all the selections have been performed.

##### 2.2.1 Group name and creation

It is possible to quickly save the cage(s) or animal(s) selection just performed. Clicking on the small icon ⋮

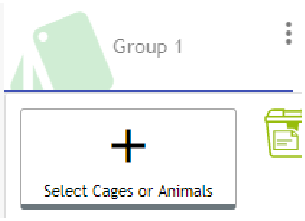

You can then change the name (by default it is Group 1), color in the graphs and finally save the group:

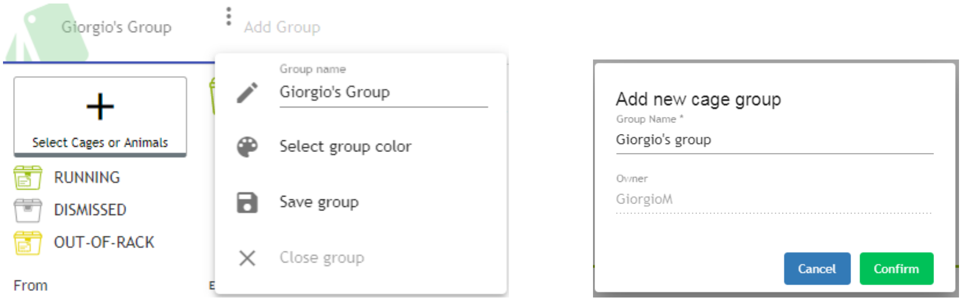

##### 2.2.2 Selection by event

Any period (From - To) can be either selected clicking on the date (and a specific date pop-up appears) or clicking on the event button:

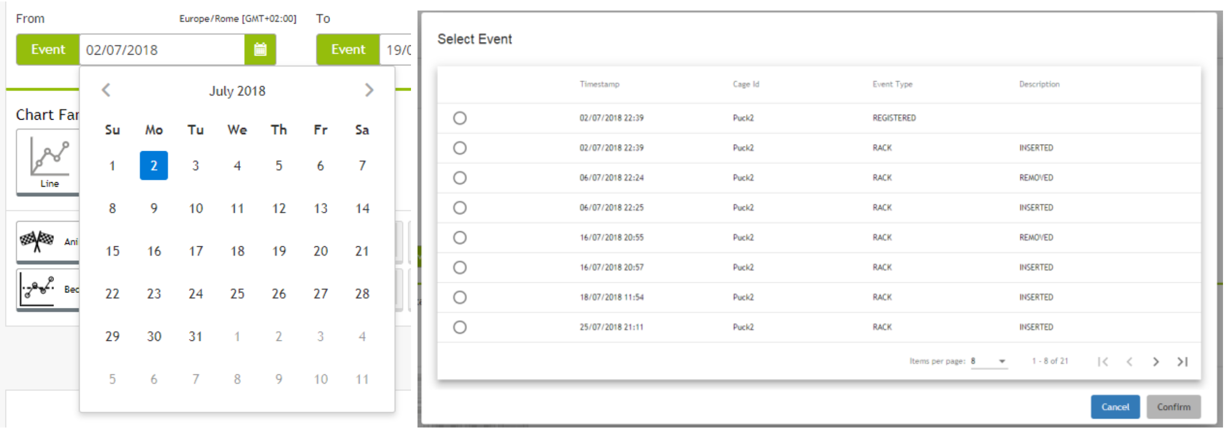

In this latter case, the entire (event) history of the selected element (cage(s) or animal(s)) appears in the chronological list. The currently managed events received from the usage of the DVC® systems are:

When the element is a Cage:

- **REGISTERED** (when the cage has been registered into the DVC® system)
- **INSERTED** (when the cage has been inserted into the DVC® rack)

- **REMOVED** (when the cage has been removed from the DVC® rack)
- **BEDDING_CHANGE** (when the cage bedding has been properly changed in the DVC® system)
- **CAGE_DISMISS** (when the cage has been properly terminated in the DVC® system).
- **RECONCILIATION** (when the cage has been reconciliated in the DVC® system, i.e. from cage missing to specific inserted cage. This is a functionality implemented in the DVC® system)

When the element is an Animal, it also has some extra events:

- **MOVED** (when the animal has been moved from a registered cage to the current under analysis)
- **CULLED** (when the animal is culled)
- **ADD** (when the animal is added for the first-time in the cage. It could be during the cage registration but also added from scratch in an already existing cage)

**PLEASE NOTE**: To disable the time selection click on the time disabling icon 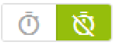 and the time will be considered from 00:00

#### 2.3 Select Chart Type

There are 3 different choices for you to start displaying data: Line, Other and Live.

##### 2.3.1.1 Chart Family Line

Selecting "Line", by default the interface proposes you the standard selection composed by the next selections:

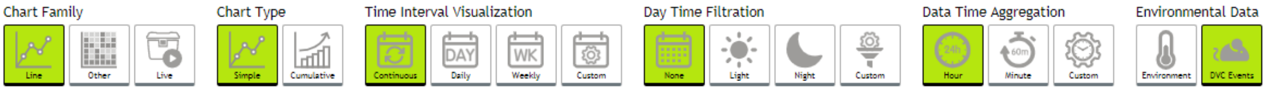

Chart Type:

- **Simple**: displays a continuous line in the selected time. Every point is the data collected by the DVC® system.
- **Cumulative**: all data are summed up and show a cumulative progression of the selected data along the selected time

Time Interval Visualization:

- **Continuous**: the line is continuously showing data along the selected time interval
- **Daily**: 7 different graphs corresponding to the 7 days of the week are displayed. If the time interval is longer than 1-week, multiple lines will be created for each day of the week.
- **Weekly**: one single weekly graph showing multiple lines corresponding to the multiple weeks (if any) in the selected period.
- **Custom**: multiple weekly graph based on number of selected weeks.

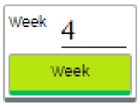

Day Time Filtration:

- **None**: no filtration applied. All day data are displayed.
- **Light**: Only light data are showed (light period based on Facility settings)
- **Night**: Only dark data are showed (night period based on Facility settings)
- **Custom**: it is possible to select days and hours to show data

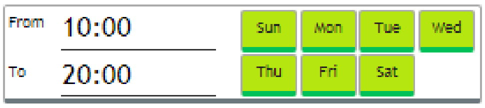

Data Time Aggregation:

- **Hour**: data are aggregated by hour
- **Minute**: data are aggregated by minute
- **Custom**: data can be aggregated by multiple minutes or multiple hours

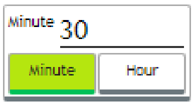

##### 2.3.1.2 Chart Family Other

Selecting "Other", by default the interface proposes you the standard selection composed by the next selections:

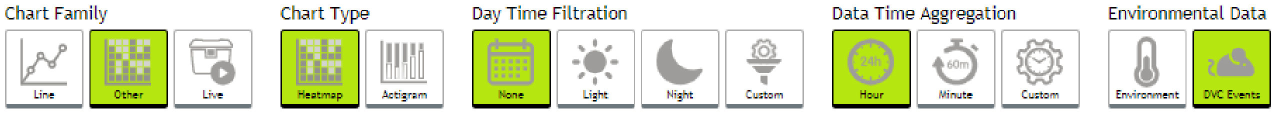

The only differences compared to the previous "Line" option is on the Chart Type selection that offers 2 different types of charts:

- **Heatmap**: data are aggregated day by day (each line is 24h data – from midnight to midnight) and every point is corresponding to the "Data Time Aggregation" selection (hour, minute, custom). The data are color-coded (blue is a low value, red is a high value)
- **Actigram**: data are aggregated day by day (each line is 24h data – from midnight to midnight) and every point is corresponding to minute aggregation (it is not possible to select other aggregation in this current version) and the magnitude of the line is corresponding to the data value collected.

##### 2.3.1.3 Chart Family Live

The DVC® Analytics features a very powerful opportunity related to "see" live data coming from selected elements (cage(s)/animal(s)).

Selecting "Live", by default the interface proposes to "see" the last 15min of the selected elements.

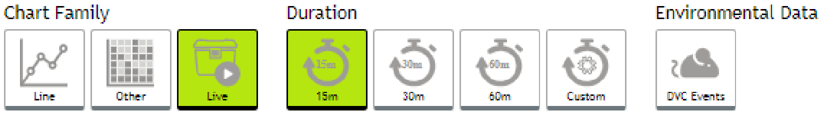

You can select other predefined time intervals (15-30-60min) as well as setting custom's ones simply clicking on the corresponding icon and then inputting your choices.

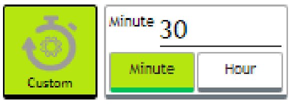

In this powerful feature, data are updated every minute.

**PLEASE NOTE**: this feature is available ONLY for RUNNING cages.

##### 2.3.2 Select Metric Type

In your current version, there are 2 different DVC® metrics available:

- **Animal Locomotion Index**: it is expressed in arbitrary unit normalized between 0% and 100% representative of the animal activity performed in the cage by the animals (no activity = 0% - all electrodes simultaneously activated by the movements of the animals = 100%)
- **Bedding Status Index**: it is expressed in arbitrary unit representative of the dielectric (cage, bedding, moisture) materials immersed into the Electromagnetic Field (EMF) generated by the DVC® board.

Moreover, it is possible to apply these metrics only to specific set of electrodes of the DVC® board simply selecting the proper icon representing the board next to animal locomotion index and Bedding status index button. The representation can be divided into: all, front, rear, corner, wall, center.

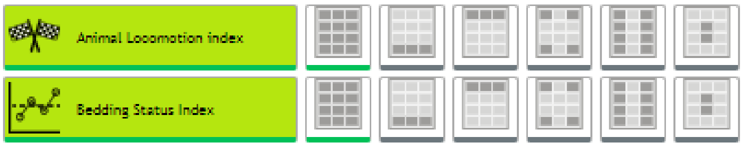

**PLEASE NOTE**: In this version of the DVC® Analytics you can select multiple metrics at once.

##### 2.3.3 Run the analysis

Once all the selections have been made (elements, time intervals, charts, metrics), it is possible to start analyzing data (the buttons are now activated and they can be clicked).

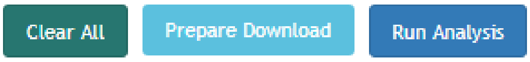

Clicking on “Run Analysis”, dependently on your specific selections, the corresponding graphs appear.

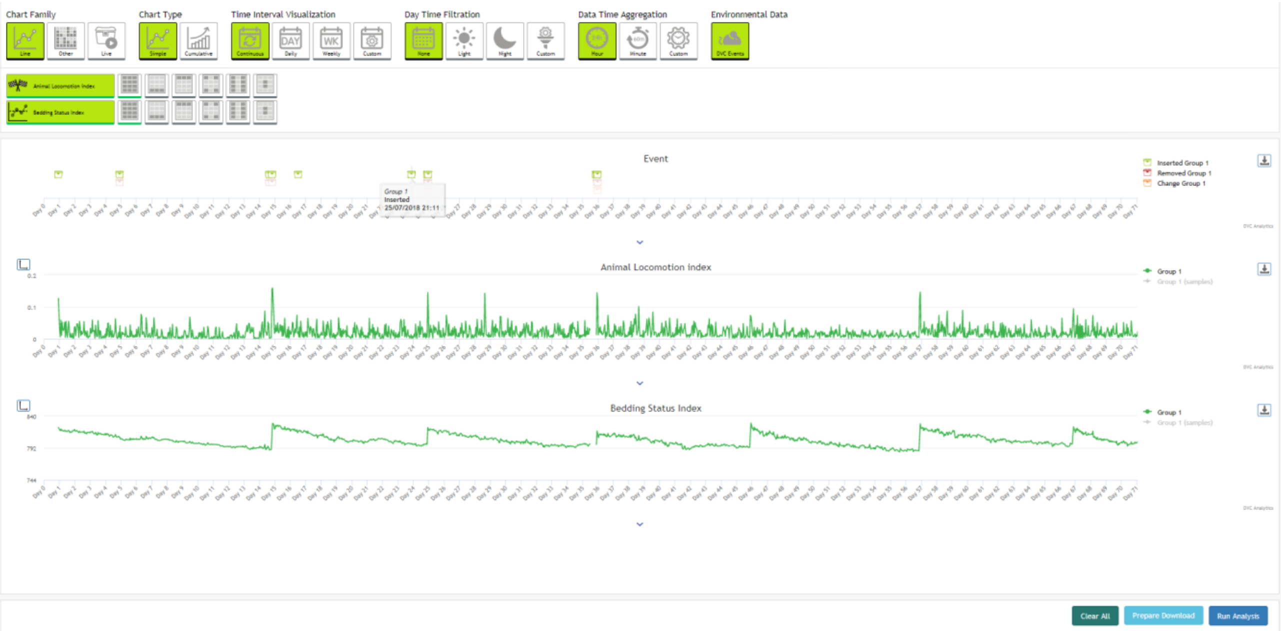

You can easily zoom any graph simply clicking in any position and keep pressed the left mouse button till the end of the area you want to zoom:

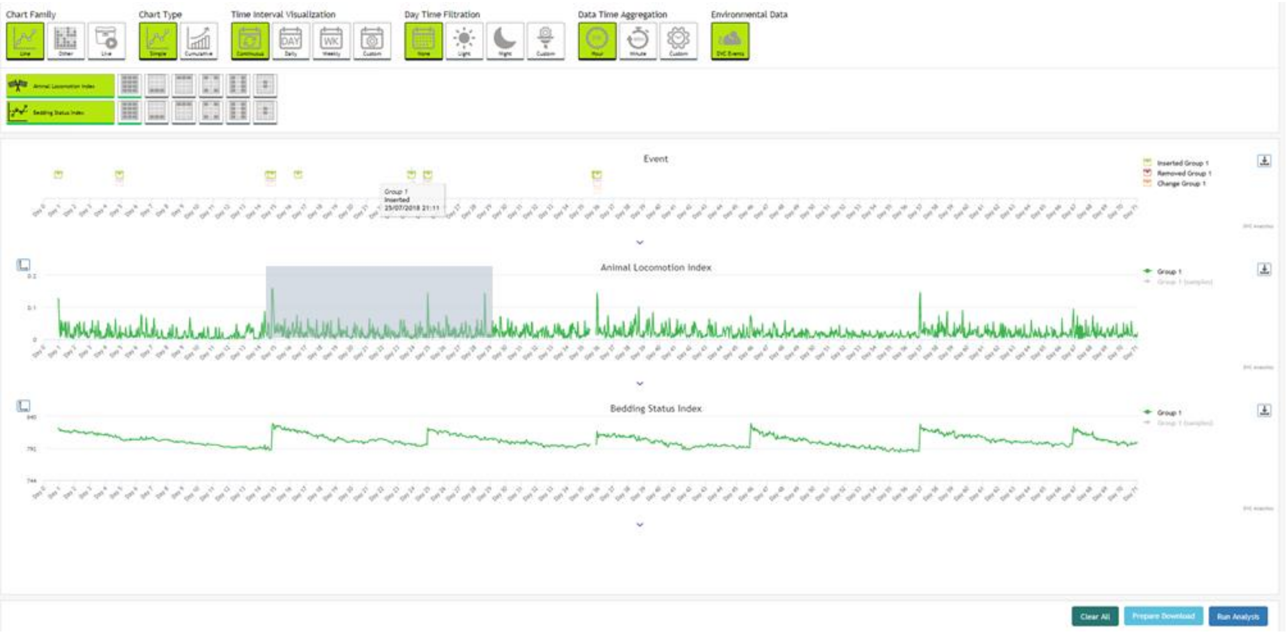

Releasing the button of the mouse, the graph is zooming in the selected area as well as all the other graphs are zoomed to show the corresponding data:

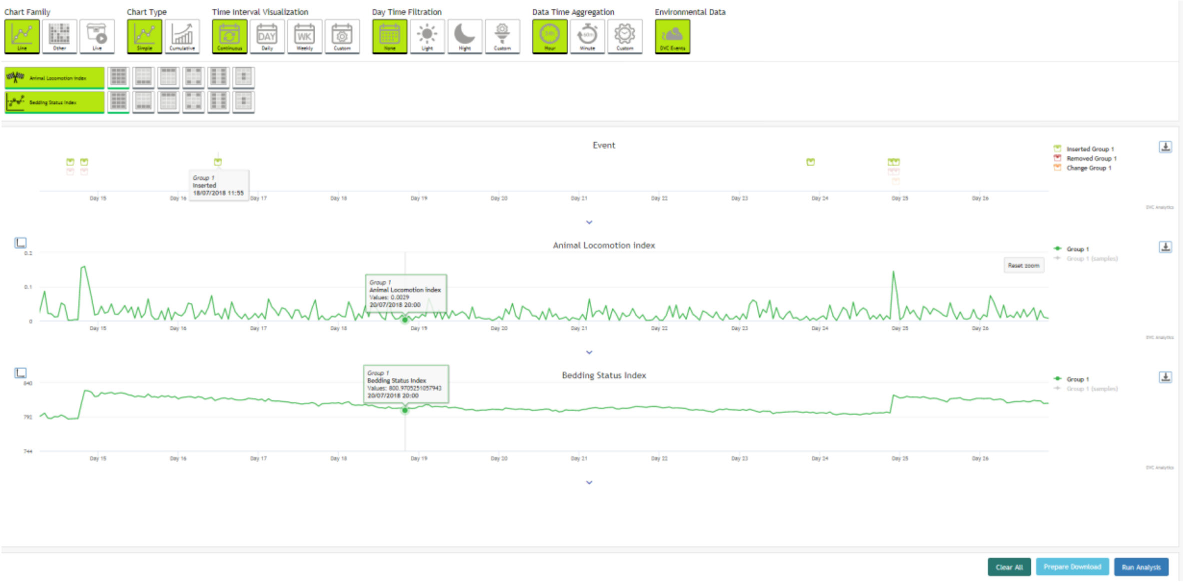

To reset the zoom, simply click on the corresponding icon of the zoomed graph Reset zoom .

Moreover, if you want to better analyze any graph, there is the opportunity to magnify it simply clicking on the corresponding icon ˅ below the graph itself.

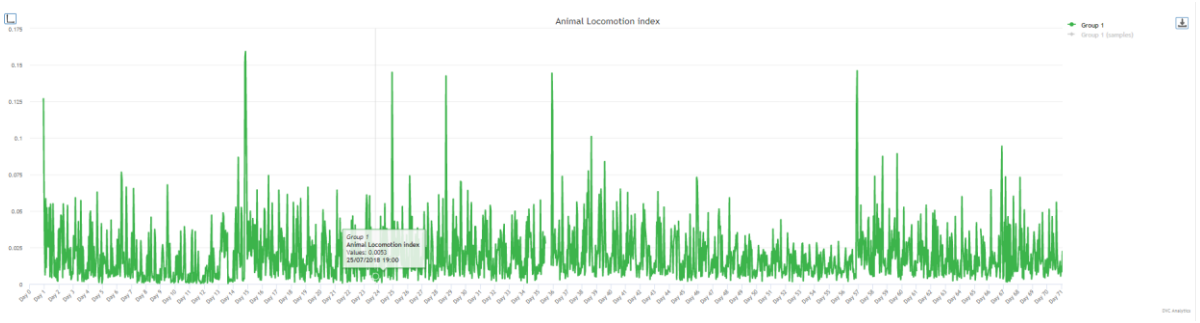

Furthermore, when you perform comparison between different groups, by default the y-axis autoscales on the highest value present in the selected time window. Otherwise, you can manually set the range of y-axis (minimum and maximum) accordingly to your need simply by clicking on this icon 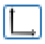 , located on the top-left side of the graph.

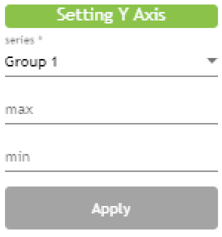

##### 2.3.3.1 Multiple groups – SEM, INTERQUARTILE and BOX PLOT

Selecting multiple groups while performing the "Data Analysis" enables extra functionalities of the application. Selecting "Line", in the "Chart Type" is then possible to select also the Simple Line with Standard Error of the Mean (SEM) or INTERQUARTILE. In this case, extra vertical lines (SEM or INTERQUARTILE) are added to any extra point of the consequent graphs.

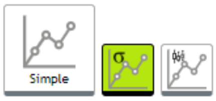

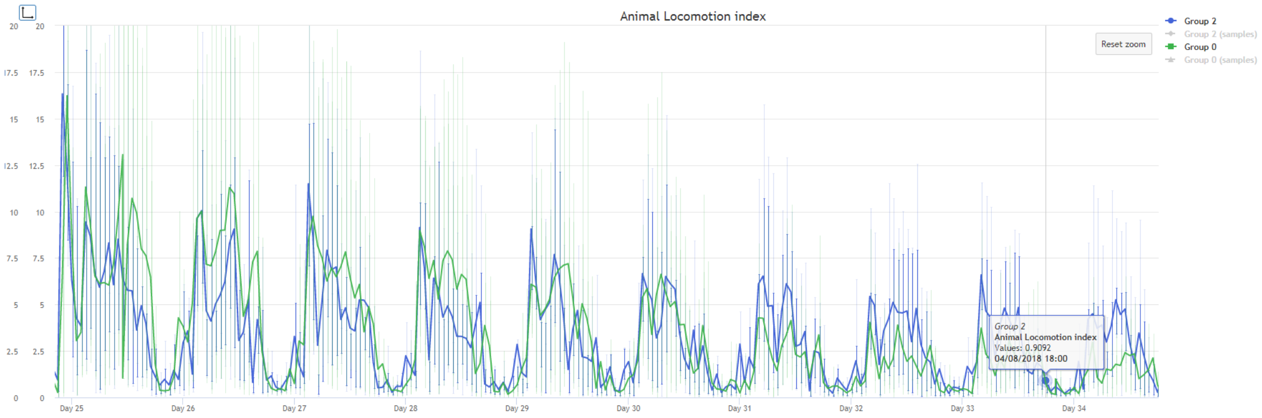

If you select "Other" in the Chart Family, a new graph becomes available "Box Plot":

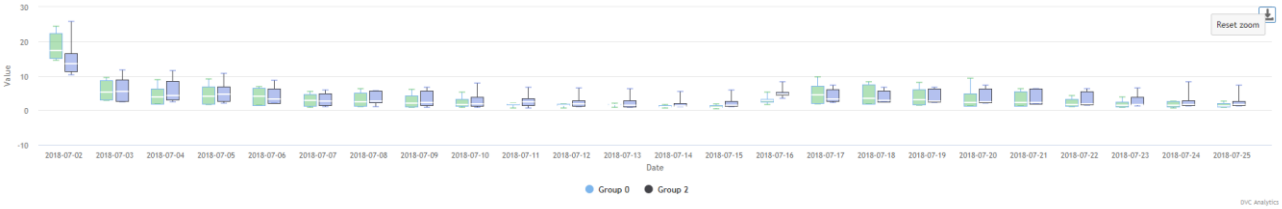

#### 2.3.4 Prepare Download

When all the selections have been performed, it is also possible to prepare the download of the corresponding selected data.

Clicking on the icon **Prepare Download** it is then possible to choose between 2 different options:

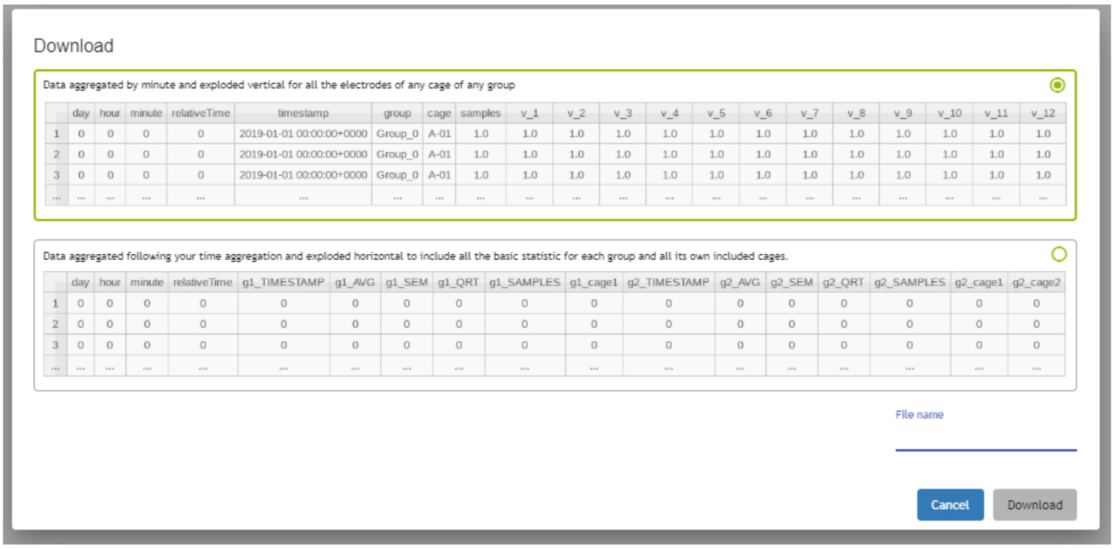

**Type 1 (upper choice):**

Reporting data aggregated in rows by minute or by hour (or custom aggregation, dependently on your specific selection) for all the electrodes of any cage of any group In sequential order.

| | day | hour | minute | relativeTime | timestamp | group | cage | samples | v_1 | v_2 | v_3 | v_4 | v_5 | v_6 | v_7 | v_8 | v_9 | v_10 | v_11 | v_12 |
|---|---|---|---|---|---|---|---|---|---|---|---|---|---|---|---|---|---|---|---|---|
| 1 | 0 | 0 | 0 | 0 | 2019-01-01 00:00:00+0000 | Group_0 | A-01 | 1.0 | 1.0 | 1.0 | 1.0 | 1.0 | 1.0 | 1.0 | 1.0 | 1.0 | 1.0 | 1.0 | 1.0 | 1.0 |
| 2 | 0 | 0 | 0 | 0 | 2019-01-01 00:00:00+0000 | Group_0 | A-01 | 1.0 | 1.0 | 1.0 | 1.0 | 1.0 | 1.0 | 1.0 | 1.0 | 1.0 | 1.0 | 1.0 | 1.0 | 1.0 |
| 3 | 0 | 0 | 0 | 0 | 2019-01-01 00:00:00+0000 | Group_0 | A-01 | 1.0 | 1.0 | 1.0 | 1.0 | 1.0 | 1.0 | 1.0 | 1.0 | 1.0 | 1.0 | 1.0 | 1.0 | 1.0 |
| ... | ... | ... | ... | ... | ... | ... | ... | ... | ... | ... | ... | ... | ... | ... | ... | ... | ... | ... | ... | ... |

| | |
|---|---|
| **Day** | Day from date selected in the observation time |
| **Hour** | Hour of that day 1-24 |
| **Minute** | Minute of that hour 1-60 |
| **relative Time** | Absolute value (in seconds) of the timing from the previous midnight of the starting date |
| **Timestamp** | Absolute Time Stamp in UTC time (https://en.wikipedia.org/wiki/Coordinated_Universal_Time) |
| **Group** | Name Group |
| **Cage** | Cage name within the selected cage group |

| | |
|---|---|
| **Samples** | Number of collected samples, detected in the specified time frame. Such information are useful to understand if the cage has been removed from the rack thus providing an estimation on how much data have been lost due to this event. |
| **V_1/2/3/4/5/6/7/8/9/10/11/12** | Value of activity detected on electrode 1/2/3/4/5/6/7/8/9/10/12 of the DVC® board |

**Type 2 (lower choice):**

Reporting data aggregated in rows following your time aggregation (minute, hour or custom) and cage groups and cages by coloumns with some basic descriptive statistics (i.e. average, quartile, SEM)

| | day | hour | minute | relativeTime | g1_TIMESTAMP | g1_AVG | g1_SEM | g1_QRT | g1_SAMPLES | g1_cage1 | g2_TIMESTAMP | g2_AVG | g2_SEM | g2_QRT | g2_SAMPLES | g2_cage1 | g2_cage2 |
|---|---|---|---|---|---|---|---|---|---|---|---|---|---|---|---|---|---|
| 1 | 0 | 0 | 0 | 0 | 0 | 0 | 0 | 0 | 0 | 0 | 0 | 0 | 0 | 0 | 0 | 0 | 0 |
| 2 | 0 | 0 | 0 | 0 | 0 | 0 | 0 | 0 | 0 | 0 | 0 | 0 | 0 | 0 | 0 | 0 | 0 |
| 3 | 0 | 0 | 0 | 0 | 0 | 0 | 0 | 0 | 0 | 0 | 0 | 0 | 0 | 0 | 0 | 0 | 0 |
| ... | ... | ... | ... | ... | ... | ... | ... | ... | ... | ... | ... | ... | ... | ... | ... | ... | ... |

| | |
|---|---|
| **day** | Day from date selected on the observation time |
| **hour** | Hour of that day 1-24 |
| **minute** | Minute of that hour 1-60 |
| **relative Time** | Absolute value (in seconds) of the timing from the previous midnight of the starting date |
| **g1_TIMESTAMP** | Absolute Time Stamp in UTC time for group 1 (https://en.wikipedia.org/wiki/Coordinated_Universal_Time) |
| **g1_AVG** | Metric Average of group 1 |
| **g1_SEM** | SEM Group 1 |
| **g1_QRT** | Quartile Group 1<br>[Minimum, Lower Quartile, Median, Upper Quartile, Maximum] |
| **g1_SAMPLES** | Collected samples, detected in the specified time frame and for the entire group. Such information are useful to understand if the cage has been removed from the rack thus providing an estimation on how much data have been lost due to this event. |
| **g1_cage1** | Metric Average of the first selected cage within group 1 |
| **g2_TIMESTAMP** | Absolute Time Stamp group 2 |
| **g2_AVG** | Metric Average of group 2 |
| **g2_SEM** | Standard Error Group 2 |
| **g2_QRT** | Quartile Group 2<br>[Minimum, Lower Quartile, Median, Upper Quartile, Maximum] |
| **g2_SAMPLES** | Amount of information, samples, detected in the specified time frame and for the entire group. Such information are useful to understand if the cage has been removed from the rack thus providing an estimation on how much data have been lost due to this event. |
| **g2_cage1** | Metric Average of the first selected cage within group 2 |
| **g2_cage2** | Metric Average of the second selected cage within group 2 |

In order to start preparing the Download, it is mandatory to fill the File name section to activate the Download button

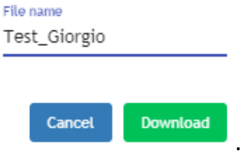

**PLEASE NOTE**: you find the "Downloaded data" in the dedicated section called Download Area

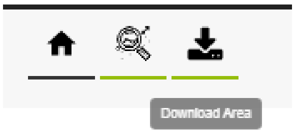

Where you can finally now really download on your PC the data simply clicking on the icon.

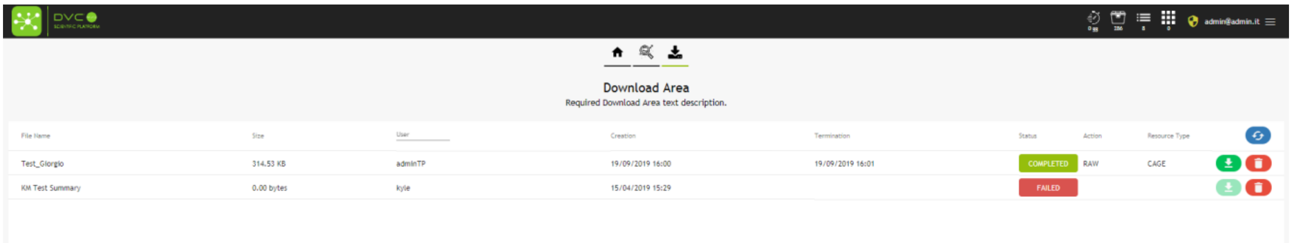

The file you download is a .zip file that contains .csv files (comma separated file), one each metric you have previously selected (in this case average = Bedding Status Index, activation = Animal Locomotion Index, events = DVC events)

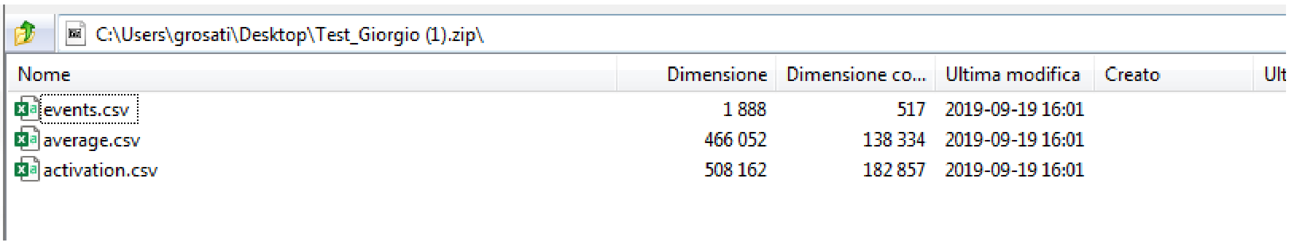

C:\Users\grosati\Desktop\Test_Giorgio (1).zip\

| Nome | Dimensione | Dimensione co... | Ultima modifica | Creato | Ult |
|---|---|---|---|---|---|
| events.csv | 1 888 | 517 | 2019-09-19 16:01 | | |
| average.csv | 466 052 | 138 334 | 2019-09-19 16:01 | | |
| activation.csv | 508 162 | 182 857 | 2019-09-19 16:01 | | |

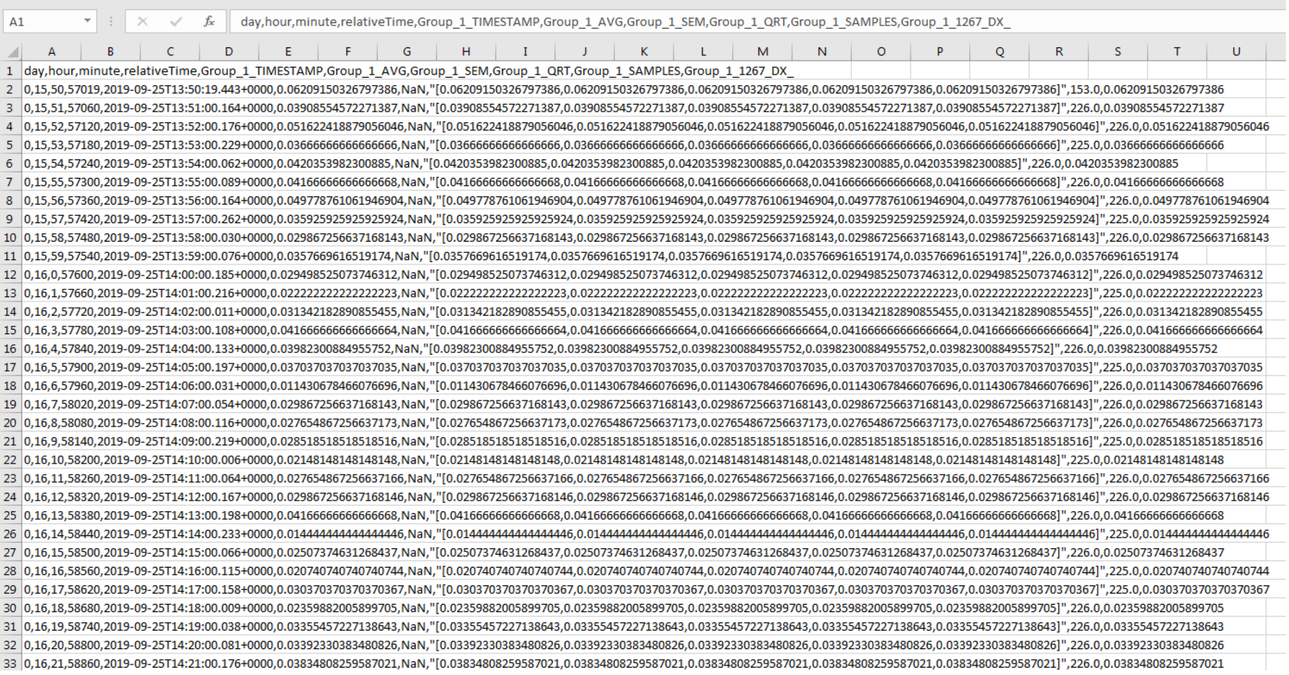

| | |
|---|---|
| 1 | day,hour,minute,relativeTime,Group_1_TIMESTAMP,Group_1_AVG,Group_1_SEM,Group_1_QRT,Group_1_SAMPLES,Group_1_1267_DX_ |
| 2 | 0,15,50,57019,2019-09-25T13:50:19.443+0000,0.06209150326797386,NaN,"[0.06209150326797386,0.06209150326797386,0.06209150326797386,0.06209150326797386,0.06209150326797386]",153.0,0.06209150326797386 |
| 3 | 0,15,51,57060,2019-09-25T13:51:00.164+0000,0.03908554572271387,NaN,"[0.03908554572271387,0.03908554572271387,0.03908554572271387,0.03908554572271387,0.03908554572271387]",226.0,0.03908554572271387 |
| 4 | 0,15,52,57120,2019-09-25T13:52:00.176+0000,0.051622418879056046,NaN,"[0.051622418879056046,0.051622418879056046,0.051622418879056046,0.051622418879056046,0.051622418879056046]",226.0,0.051622418879056046 |
| 5 | 0,15,53,57180,2019-09-25T13:53:00.229+0000,0.0366666666666666,NaN,"[0.0366666666666666,0.0366666666666666,0.0366666666666666,0.0366666666666666,0.0366666666666666]",225.0,0.0366666666666666 |
| 6 | 0,15,54,57240,2019-09-25T13:54:00.062+0000,0.0420353982300885,NaN,"[0.0420353982300885,0.0420353982300885,0.0420353982300885,0.0420353982300885,0.0420353982300885]",226.0,0.0420353982300885 |
| 7 | 0,15,55,57300,2019-09-25T13:55:00.089+0000,0.04166666666666668,NaN,"[0.04166666666666668,0.04166666666666668,0.04166666666666668,0.04166666666666668,0.04166666666666668]",226.0,0.04166666666666668 |
| 8 | 0,15,56,57360,2019-09-25T13:56:00.164+0000,0.049778761061946904,NaN,"[0.049778761061946904,0.049778761061946904,0.049778761061946904,0.049778761061946904,0.049778761061946904]",226.0,0.049778761061946904 |
| 9 | 0,15,57,57420,2019-09-25T13:57:00.262+0000,0.035925925925925924,NaN,"[0.035925925925925924,0.035925925925925924,0.035925925925925924,0.035925925925925924,0.035925925925925924]",225.0,0.035925925925925924 |
| 10 | 0,15,58,57480,2019-09-25T13:58:00.030+0000,0.029867256637168143,NaN,"[0.029867256637168143,0.029867256637168143,0.029867256637168143,0.029867256637168143,0.029867256637168143]",226.0,0.029867256637168143 |
| 11 | 0,15,59,57540,2019-09-25T13:59:00.076+0000,0.0357669616519174,NaN,"[0.0357669616519174,0.0357669616519174,0.0357669616519174,0.0357669616519174,0.0357669616519174]",226.0,0.0357669616519174 |
| 12 | 0,16,0,57600,2019-09-25T14:00:00.185+0000,0.029498525073746312,NaN,"[0.029498525073746312,0.029498525073746312,0.029498525073746312,0.029498525073746312,0.029498525073746312]",226.0,0.029498525073746312 |
| 13 | 0,16,1,57660,2019-09-25T14:01:00.216+0000,0.022222222222222223,NaN,"[0.022222222222222223,0.022222222222222223,0.022222222222222223,0.022222222222222223,0.022222222222222223]",225.0,0.022222222222222223 |
| 14 | 0,16,2,57720,2019-09-25T14:02:00.011+0000,0.03134218289085545,NaN,"[0.03134218289085545,0.03134218289085545,0.03134218289085545,0.03134218289085545,0.03134218289085545]",226.0,0.03134218289085545 |
| 15 | 0,16,3,57780,2019-09-25T14:03:00.108+0000,0.04166666666666664,NaN,"[0.04166666666666664,0.04166666666666664,0.04166666666666664,0.04166666666666664,0.04166666666666664]",226.0,0.04166666666666664 |
| 16 | 0,16,4,57840,2019-09-25T14:04:00.133+0000,0.03982300884955752,NaN,"[0.03982300884955752,0.03982300884955752,0.03982300884955752,0.03982300884955752,0.03982300884955752]",226.0,0.03982300884955752 |
| 17 | 0,16,5,57900,2019-09-25T14:05:00.197+0000,0.037037037037037035,NaN,"[0.037037037037037035,0.037037037037037035,0.037037037037037035,0.037037037037037035,0.037037037037037035]",225.0,0.037037037037037035 |
| 18 | 0,16,6,57960,2019-09-25T14:06:00.031+0000,0.011430678466076696,NaN,"[0.011430678466076696,0.011430678466076696,0.011430678466076696,0.011430678466076696,0.011430678466076696]",226.0,0.011430678466076696 |
| 19 | 0,16,7,58020,2019-09-25T14:07:00.054+0000,0.029867256637168143,NaN,"[0.029867256637168143,0.029867256637168143,0.029867256637168143,0.029867256637168143,0.029867256637168143]",226.0,0.029867256637168143 |
| 20 | 0,16,8,58080,2019-09-25T14:08:00.116+0000,0.027654867256637173,NaN,"[0.027654867256637173,0.027654867256637173,0.027654867256637173,0.027654867256637173,0.027654867256637173]",226.0,0.027654867256637173 |
| 21 | 0,16,9,58140,2019-09-25T14:09:00.219+0000,0.028518518518518516,NaN,"[0.028518518518518516,0.028518518518518516,0.028518518518518516,0.028518518518518516,0.028518518518518516]",225.0,0.028518518518518516 |
| 22 | 0,16,10,58200,2019-09-25T14:10:00.006+0000,0.0214814814814148148,NaN,"[0.0214814814814148148,0.0214814814814148148,0.0214814814814148148,0.0214814814814148148,0.0214814814814148148]",225.0,0.0214814814814148148 |
| 23 | 0,16,11,58260,2019-09-25T14:11:00.064+0000,0.027654867256637166,NaN,"[0.027654867256637166,0.027654867256637166,0.027654867256637166,0.027654867256637166,0.027654867256637166]",226.0,0.027654867256637166 |
| 24 | 0,16,12,58320,2019-09-25T14:12:00.167+0000,0.029867256637168146,NaN,"[0.029867256637168146,0.029867256637168146,0.029867256637168146,0.029867256637168146,0.029867256637168146]",226.0,0.029867256637168146 |
| 25 | 0,16,13,58380,2019-09-25T14:13:00.198+0000,0.04166666666666668,NaN,"[0.04166666666666668,0.04166666666666668,0.04166666666666668,0.04166666666666668,0.04166666666666668]",226.0,0.04166666666666668 |
| 26 | 0,16,14,58440,2019-09-25T14:14:00.233+0000,0.014444444444444446,NaN,"[0.014444444444444446,0.014444444444444446,0.014444444444444446,0.014444444444444446,0.014444444444444446]",225.0,0.014444444444444446 |
| 27 | 0,16,15,58500,2019-09-25T14:15:00.066+0000,0.02507374631268437,NaN,"[0.02507374631268437,0.02507374631268437,0.02507374631268437,0.02507374631268437,0.02507374631268437]",226.0,0.02507374631268437 |
| 28 | 0,16,16,58560,2019-09-25T14:16:00.115+0000,0.020740740740740744,NaN,"[0.020740740740740744,0.020740740740740744,0.020740740740740744,0.020740740740740744,0.020740740740740744]",225.0,0.020740740740740744 |
| 29 | 0,16,17,58620,2019-09-25T14:17:00.158+0000,0.030370370370370367,NaN,"[0.030370370370370367,0.030370370370370367,0.030370370370370367,0.030370370370370367,0.030370370370370367]",225.0,0.030370370370370367 |
| 30 | 0,16,18,58680,2019-09-25T14:18:00.009+0000,0.0235988200589905,NaN,"[0.0235988200589905,0.0235988200589905,0.0235988200589905,0.0235988200589905,0.0235988200589905]",226.0,0.0235988200589905 |
| 31 | 0,16,19,58740,2019-09-25T14:19:00.038+0000,0.0335545722713643,NaN,"[0.0335545722713643,0.0335545722713643,0.0335545722713643,0.0335545722713643,0.0335545722713643]",226.0,0.0335545722713643 |
| 32 | 0,16,20,58800,2019-09-25T14:20:00.081+0000,0.0339233038348026,NaN,"[0.0339233038348026,0.0339233038348026,0.0339233038348026,0.0339233038348026,0.0339233038348026]",226.0,0.0339233038348026 |
| 33 | 0,16,21,58860,2019-09-25T14:21:00.176+0000,0.03834808259587021,NaN,"[0.03834808259587021,0.03834808259587021,0.03834808259587021,0.03834808259587021,0.03834808259587021]",226.0,0.03834808259587021 |

**PLEASE NOTE**: TIMESTAMP are produced in UTC time (https://en.wikipedia.org/wiki/Coordinated_Universal_Time). If you are using MS Excel to open these files, remember to set the "." as decimal separator.

#### 2.4 Manage Experiment

Clicking on the icon "Manage Experiment" other choices appear:

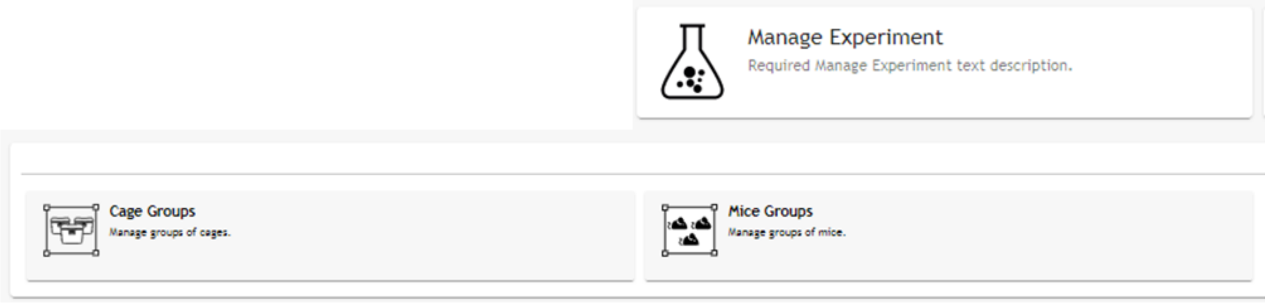

##### 2.4.1 Cage Groups

In this section it is possible to create specific groups of cages and assign to (already) existing DVC® Analytics users.

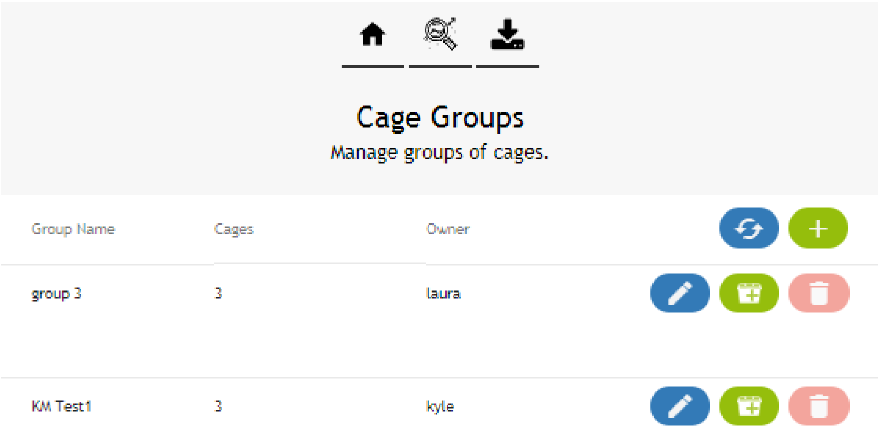

First step is to create the group, simply clicking on the corresponding icon . A specific pop-up area appears and you are requested to insert the name of this group as well as the owner of the group.

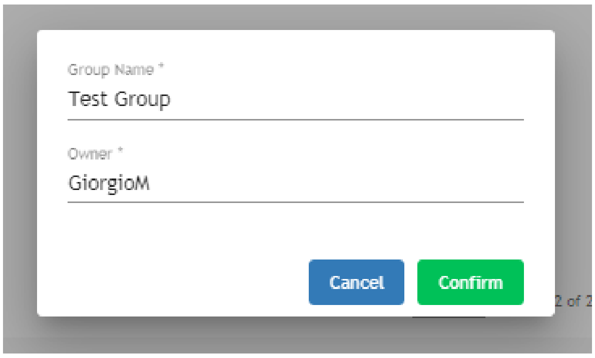

Then, you can start adding cages to this group simply clicking on the icon and then selecting the cages to be included in this group.

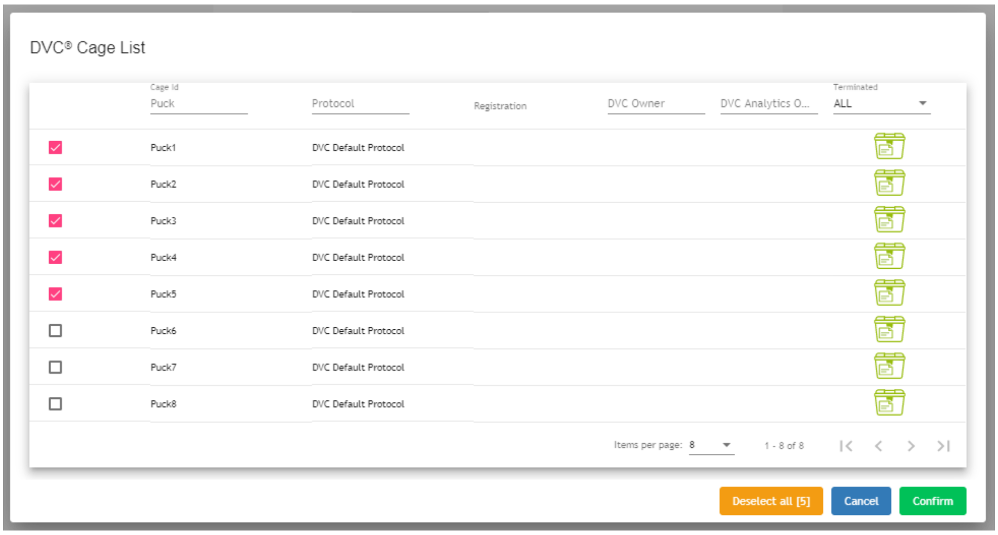

**PLEASE NOTE**: same cage(s) can be assigned to different groups.

**PLEASE NOTE**: in order to delete a group, you must deselect all the cages before being able to delete the Cage Group with the corresponding icon .

##### 2.4.2 Mice groups

To create a Mouse group, please follow exactly the previous workflow with the only different that you must choose between available mice IDs (instead of cage IDs).

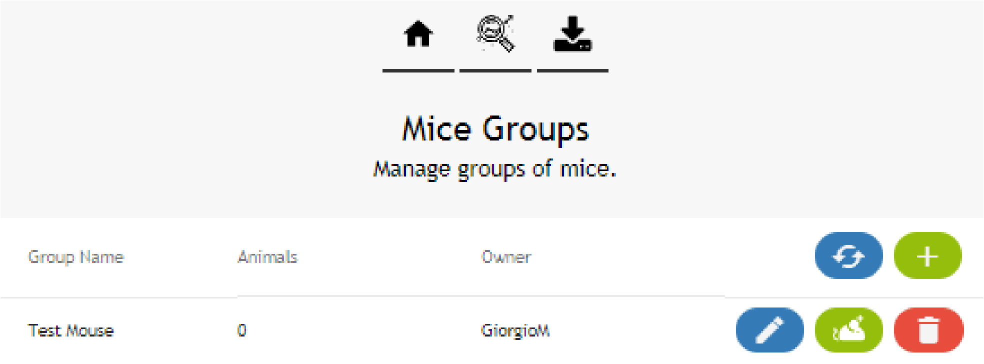

#### 2.5 Download Area

As already anticipated, in this section, you can immediately find all the requested-to-be-prepared data.

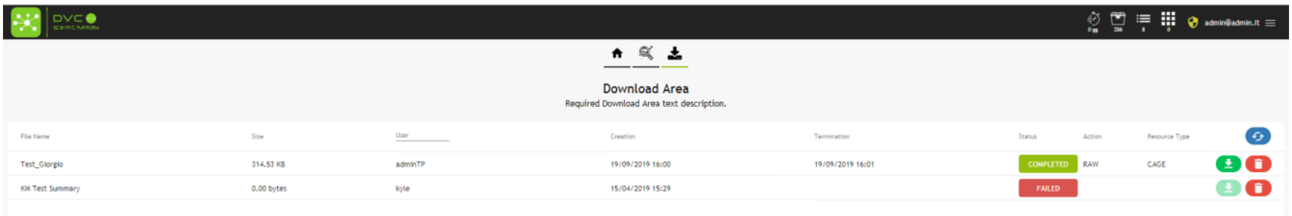

Additionally, you have other information such the status of the requested task (Running, Failed, Executed) as well as who requested it, the size of the file and the date of starting (Creation) and Termination.

#### 2.6 User settings

As described in section 2.1, this section is available only for the DVC® Analytics users registered as Facility Manager and not available to the users registered as Researchers.

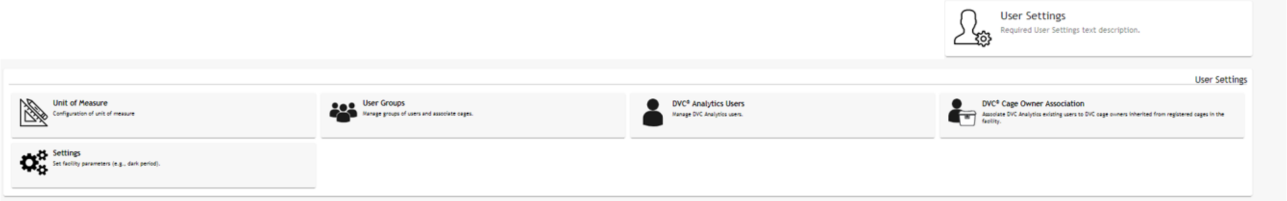

There are different options you can set

#### 2.7 Unit of Measure

Clicking on the corresponding button, you can see which are your current settings set by Tecniplast

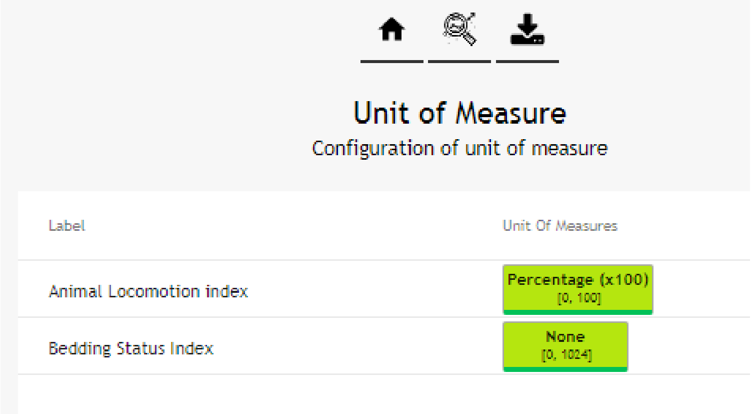

##### 2.7.1 User Groups

You can create groups of users able to access to groups of cages.

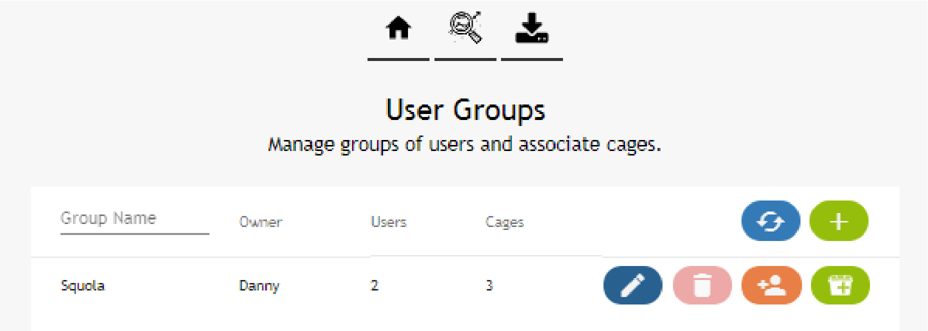

This functionality is similar to the abovementioned Cage group (sect. 2.4.1) but with the difference that more users can now access to the same cages. You can simply add more users to the group clicking on the corresponding icon and then chose from the list, as well adding cages clicking on the icon and then choosing the selected ones from the list of available.

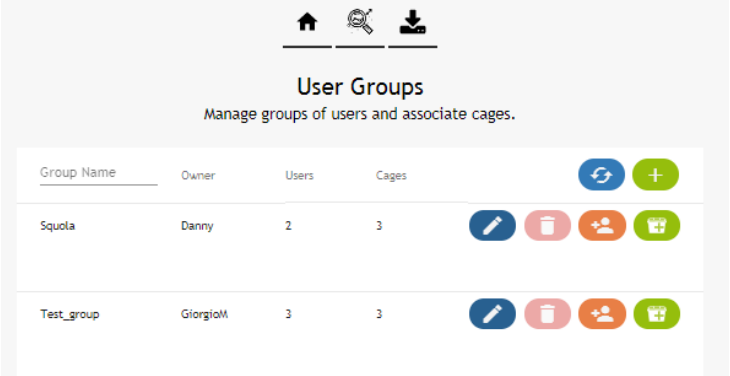

**PLEASE NOTE**: to delete a User group, you must deselect all the users and all the cages from the User group.

##### 2.7.2 DVC® Analytics Users

In this section you can create unlimited users simply clicking on the corresponding icon .

You must enter different information for any user:

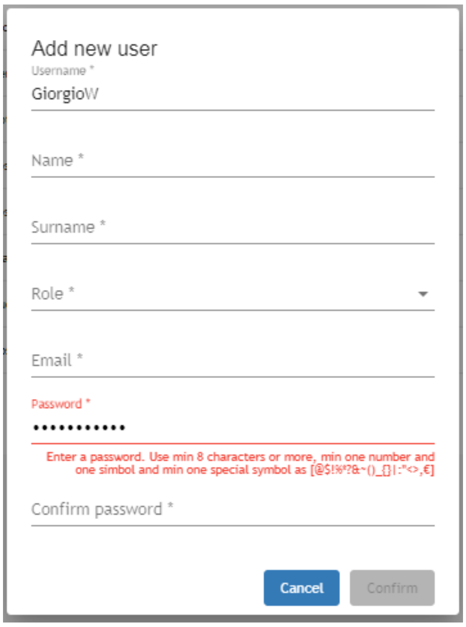

**PLEASE NOTE**: The minimum requirements to create your password are: 8 characters or more, one number and one symbol and one special symbol as [@$!%*?&~()_{}|:"<>,€]

Any new user must have an initial password that can be easily changed when entering for the first time in the application and clicking on the top right area 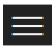

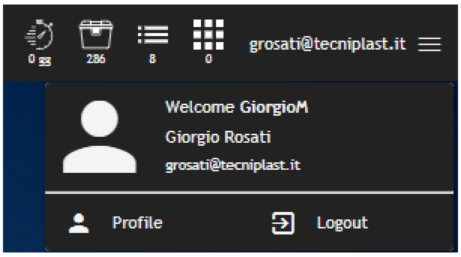

And then "Profile" and finally "Change Password" where the (new) user is requested to insert OLD password and new one.

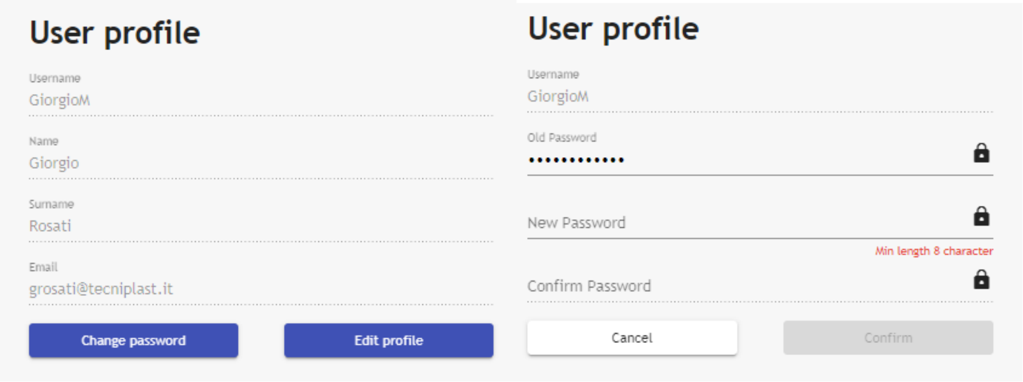

##### 2.7.3 DVC® Cage Owner Association

This section is fundamental especially if you are (or have) Researcher users, because it is the only way to analyze cages when using this profile.

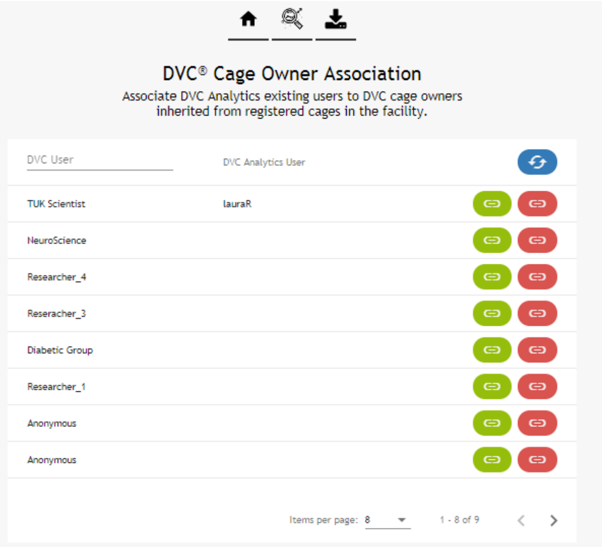

Every cage prepared in the DVC® system can have a DVC® Owner associated to it (it is not mandatory but highly suggested when used in combination with DVC® Analytics). If so, this DVC® Owner is pushed to the DVC® Analytics and it is called DVC User and can be associated to already existing DVC® Analytics users simply clicking on the icon.

**PLEASE NOTE**: only 1 DVC® Analytics user can be associate to the DVC User (it is not possible to associate multiple DVC® Analytics users to the same cages – you can manage this situation using.

##### 2.7.4 Settings

This button allows you to set different Facility information

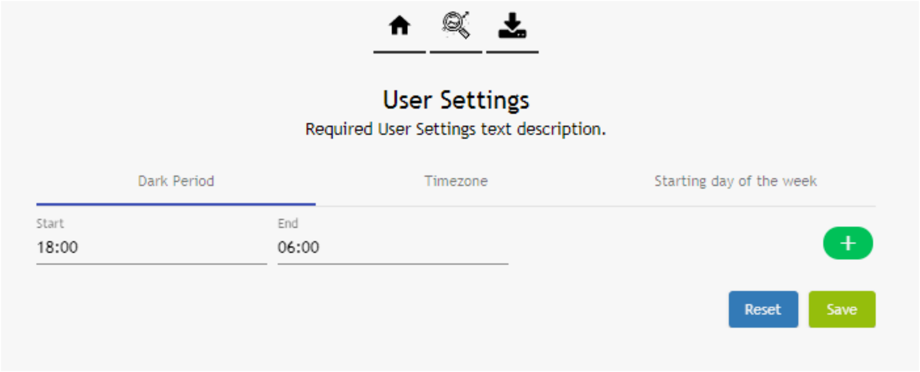

###### 2.7.4.1 Dark Period

This section is fundamental to properly set your official night period in the Facility. This setting is used in the "Day Time Filtration" section.

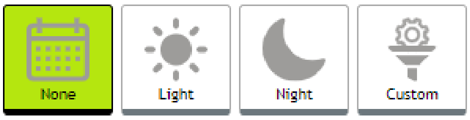

If you manage time-shift in your facility because of summer-winter time, you can click on the 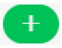 and then select the new time interval and the date of start from this interval

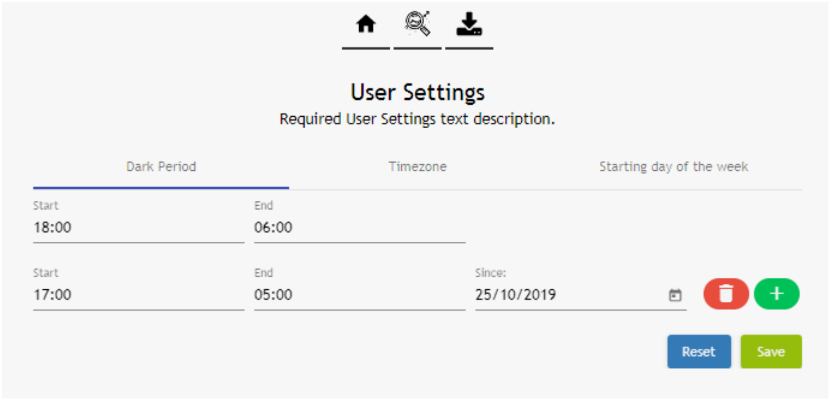

**PLEASE NOTE**: you can change/delete this time interval and original data are not affected.

###### 2.7.4.2 Time Zone

This is also important to "see" your data in the graph/charts considering your Facility Time zone. Simply start typing in the corresponding area and the field will autocomplete with your timezone

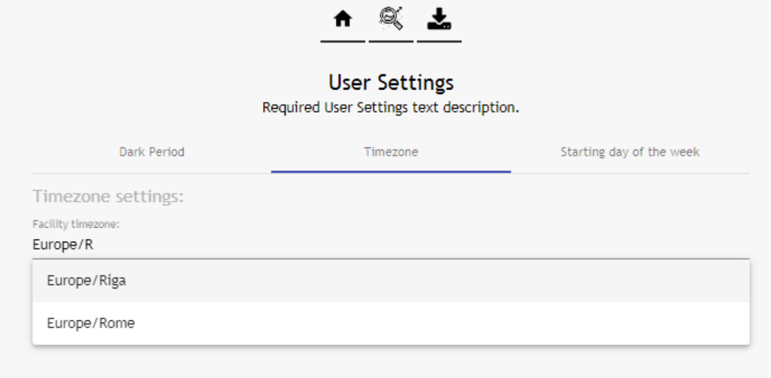

##### 2.7.4.3 Starting of the week

In order to allow you maximum flexibility, it is possible to set the "first" day of the week that will be then used to display data especially in the daily and weekly graph selection:

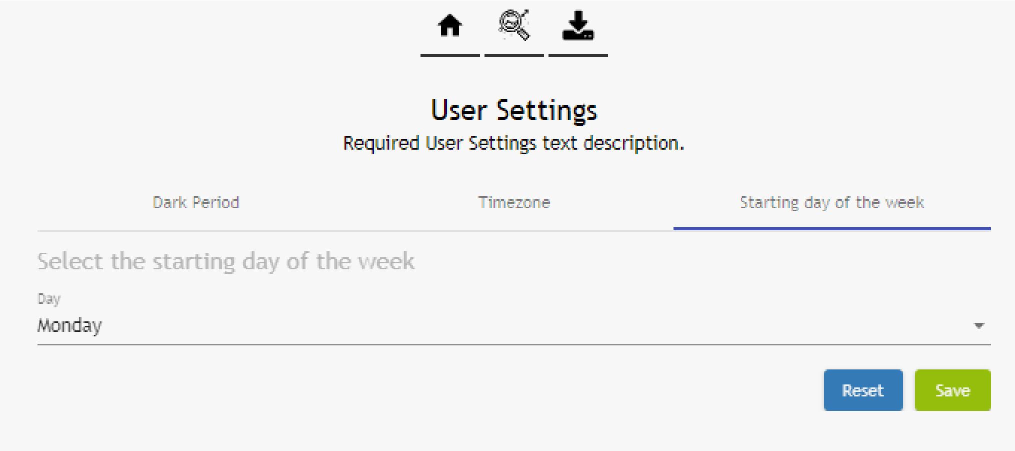

**PLEASE NOTE**: Sunday and Monday are currently the available "first" days of the week.

#### 3 Useful information

In this section, we would like to provide you some tips and information that would be important to know in order to better understand how the DVC® Analytics works and get the most from it.

##### 3.1 DVC® board

The DVC® board is the core of the DVC® system. There are 12 different electrodes that are mapping the entire base of the cage. These electrodes are numbered in the next way:

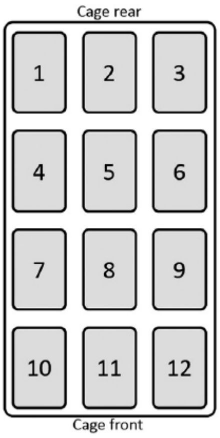

For some metrics (Animal Activity Index and Bedding Status Index) it is possible to select ONLY some electrodes (corners, walls, center, etc) if you want to deeply analyze specific patterns.

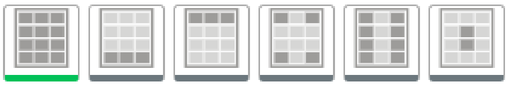

For some other metrics, such as Running Wheel or Animal Tracking for instance, it is not possible to select specific electrodes because the data are calculated using the Running Wheel or the entire DVC® board respectively.

##### 3.2 DVC® working principle and derived metrics

Basically, the working principle of the DVC® system is based on an electrical capacitance sensing technology (CTS). As said, the DVC® board is composed of 12 electrodes connected to an integrated circuit that continuously measures their electrical capacitance every 250msec (roughly). Since capacitance is influenced by the matter present in each electrode's surrounding, its measurements are affected by the presence of, e.g., water and animals. Note that, materials with high water content are characterized by large values of relative permittivity (with respect to air), which in turns has a direct effect on capacitance (high relative permittivity means higher capacitance). Since mice are characterized by high water content, their movements performed while close to an electrode induce significant capacitance changes, and thus, by properly tracking these changes over time it is possible to monitor animal activity. Note that, capacitance remains substantially unchanged when material compositions around an electrode is unvaried. Additionally, the capacitance readings are affected by the presence of water (due to e.g., bottle leakage) or urine. However, animal activity occurs on a time scale substantially different than that of water leakage or urine and thus the two variables can be easily distunguished. Furthermore, even when water/urine are present in an electrode surrounding (clearly not a flooded cage, but common amount of water/urine in a dirty cage) the capability of the system to discern animal movements is substantially unchanged. In fact, the presence of water/urine can change absolute capacitance readings, but not capacitance variations due to animal movements.

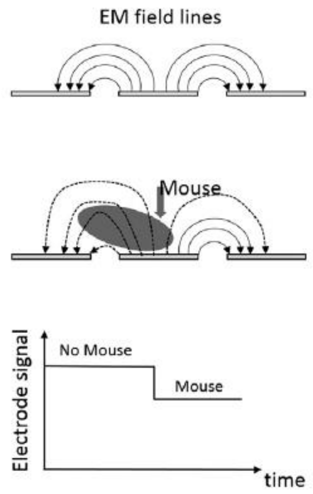

Keeping in mind this working principle, there are some important metrics that can be applied. The currently available metrics in the DVC® Analytics systems are:

- **Animal Activity Index**
- **Bedding Status Index**
- **Animal Tracking Distance**
- **Animal Tracking Speed**
- **Running Wheel Rotation**
- **Running Wheel Distance**
- **Running Wheel Speed**

**PLEASE NOTE**: every element (cage or animal) has specific capabilities assigned by the DVC® system that enable or not correspondent DVC® metrics (a cage without Running Wheel doesn't enable the Running Wheel metric).

##### 3.2.1 Animal Activity Index

This DVC® metric is extremely robust because it uses the so called "Activation Density" metric that has been extensively validated in the field across different experiments and validation processes (you can find detailed information in this publication https://www.heliyon.com/article/e01454).

An electrode is considered activated when its measurements are perturbed significantly over a limited time interval, which generally occurs when a mouse performs activity while sufficiently close to an electrode (see below). Density indicates that the total number of activations are divided by the duration of the time interval considered and the number of electrodes of interest (up to twelve). A sketch of the Capacity Sensing Technology activation density metric is the following. Recall that the Capacity Sensing Technology board provides measurements related to electrode capacitance every 250ms and let $c_k(t)$ be the (filtered) capacitance measurements from the kth electrode at time t. Then, we compare the difference between two adjacent capacitance measurements as

$$\Delta_k(t) = c_k(t) - c_k(t-1).$$

The rationale behind this is that when no animal movements occur the difference Δk(t) is approximately zero as there are no variations in electrode capacitance, while absolute values |Δk(t)|>0 indicate capacitance variations that are generally caused by animal movements. According to these observations, we consider that an electrode is activated when we observe a change in adjacent measurements larger than a fixed threshold λ. The threshold is conveniently chosen to separate noise induced capacitance variations from animal movements. Finally, the binary information indicating whether the electrode is activated at time t is given by:

$$a_k(t) = 1[|\Delta_k(t)| \geq \lambda]$$

where 1[x] is the indicator function for the event x, with 1[x]=1 if event x is true and 1[x]=0 otherwise. Finally, one is generally interested in measuring the average amount of activations, occurring across a given set of electrodes (i.e., area of the cage) and within a given time interval. To do so, the CST activation density within time periods t1 and t2, across set of electrode S, can be computed as

$$A_{CST}(t_1, t_2) = \frac{1}{|S|\,(t_2 - t_1)} \sum_{k \in S} \sum_{t=t_1}^{t_2-1} a_k(t)$$

where |S| indicates the cardinality (i.e., number of electrodes) of set S. Note that, the CST activation density does not indicate the type of movement performed, but it only accounts whether activity occurred close to an electrode.

This Animal Activity Index is expressed in % arbitrary unit and it is normalized between 0% and 100%.

##### 3.2.2 Bedding Status Index

This DVC® metric has been developed in order to provide the possibility to determine and analyze the status of the bedding. There are basically 2 different events that affect bedding status:

- **Growing moisture due to latrine creation**
- **Water flooding due to water bottle leakage and/or Automatic Watering System valve failure**

In both cases, this DVC® metric is calculated starting from the absolute value collected by the CST and applied a specific-time-interval average function (minute, hour, custom)

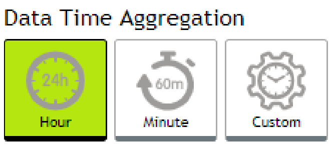

**PLEASE NOTE**: this DVC® metric lends itself well to use the different mapping electrodes opportunity

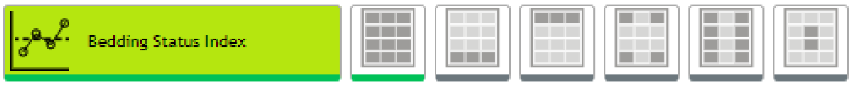

##### 3.2.3 Animal Tracking Distance and Speed

The distance walked accounts for the total distance covered by the mouse within a given time interval, while the average speed is the distance walked divided by the duration of the time interval considered. We assume that the mouse position on the cage floor is identified in terms of its centroid, while the distance walked is computed via the sum of the Euclidean distances of the mouse centroid in successive frames within the time interval of interest. The distance walked is defined as follows. Let $\mathbf{p}(t) = [p_x(t), p_y(t)]$be a 2 ×1 vector of coordinates on the plane (cage floor) representing the position of the centroid of the mouse at time $t$. Then, the distance walked within the time interval $t_1$and $t_2$can be computed as:

$$S(t_1, t_2) = \sum_{t=t_1+1}^{t_2} d(t)$$

Where:

$$d(t) = \sqrt{(p_x(t) - p_x(t-1))^2 + (p_y(t) - p_y(t-1))^2}$$

is the Euclidean distance between two positions in adjacent frames.

The average speed is instead defined as the ratio between the cumulative walked distance and the duration of the time interval:

$$V(t_1, t_2) = \frac{1}{t_2 - t_1} S(t_1, t_2)$$

##### 3.2.4 Running Wheel Rotation, Distance and Speed

Using the product DVC® Running wheel, it is possible to automate several metrics. The diameter of the plastic DVC® Running wheel is 110,4 mm (4,35 inch) that corresponds to a perimeter of about 34,54 cm (13,6 inch).

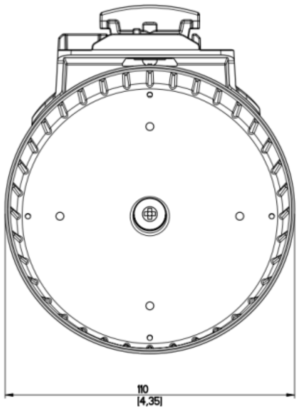

The minimum time resolution is the minute and the metrics are expressed in:

- **Running Wheel rotation:** # complete rotations in the selected time-resolution (minute, hour, custom)
- **Running Wheel distance**: # complete rotations * 34,54 cm (13,6 inch) in the selected time-resolution
- **Running Wheel Speed**: expressed in cm/min or m/min

#### 3.3 How data are calculated and aggregated

Considering the different data and the different charts, it is fundamental to understand how these are calculated.

##### 3.3.1 Line Chart

Every point of the line is calculated in the next way:

**SPATIAL AGGREGATION**: average of the selected DVC® boards electrodes (by default the 12 electrodes). The result is one single data point (if you select only the corners, the data point is the average of the 4 electrodes).

**TEMPORAL AGGREGATION:** the default temporal window (minute and hour) are automatically calculated by the DVC® system while they are occurring. Vice versa, if you have selected a custom temporal window, the result is the average of all the minutes included into the custom temporal interval if the time is not a multiple of the hour.

**Activation (3min) = [activation (min 1°) + activation (min 2°) + activation (min 3°)] / 3**

If the custom temporal interval is a multiple of the hour:

**Activation (3h) = [activation (hour 1°) + activation (hour 2°) + activation (hour 3°)] / 3**

**GROUP AGGREGATION:** the data point in the graph is the calculated as the average of the single calculated metric.

**Activation (3 cages) = [activation (cage 1°) + activation (cage 2°) + activation (cage 3°)] / 3**

##### 3.3.2 Line chart with SEM

As explained in section 2.3.3.1, the SEM is enabled by selecting multiple cages in the same group. Everything is calculated the same as above in terms of spatial and temporal aggregation and then, the SEM (Standard Error of the Mean), it is simply calculated as (where SD = Standard Deviation):

$$\textbf{SEM = SD / } \sqrt{(\#\boldsymbol{samples})}$$

In the correspondent Linechart it is shown as average (central data point) ± SEM.

##### 3.3.3 Line chart with Interquantile

As explained in section 2.3.3.1, the INTERQUANTILE is enabled by selecting multiple cages in the same group. Everything is calculated the same as above in terms of spatial and temporal aggregation and then, the INTERQUANTILE feature enables 6 different points per data sample:

- Average
- Median
- Quantile
- 3* Quantile
- Interquantile min range
- Interquantile max range

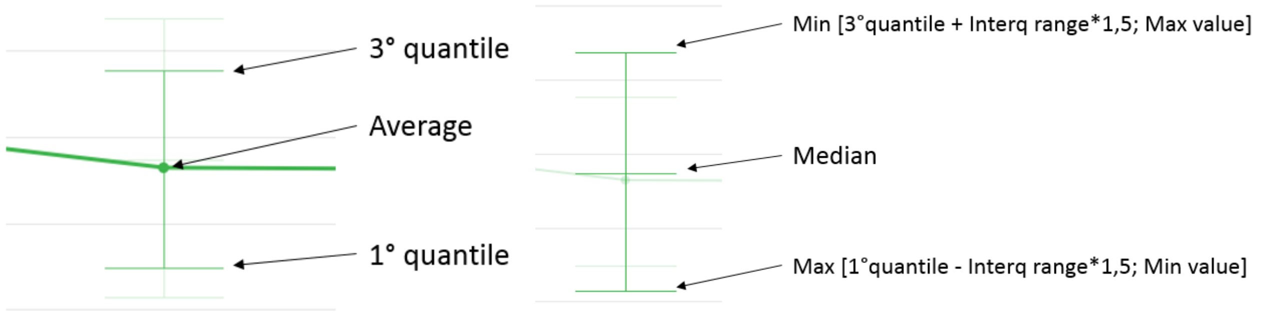

##### 3.3.4 Line Chart cumulative

Spatial and temporal aggregation follow the abovementioned scheme, and in this specific case, every data point is the sum of the previous ones:

$$d(0) = A(0)$$

$$d(1) = A(0) + A(1)$$

$$d(2) = A(0) + A(1) + A(2)$$

$$d(n) = A(0) + A(1) + A(2) + \ldots + A(n)$$

##### 3.3.5 Heatmap

Spatial, temporal and group aggregation are exactly calculated as above, the only difference with the Line Chart is the chromatic visualization (from blue as lower value to red as higher value). Every block is representative of the Data Time Aggregation chosen.

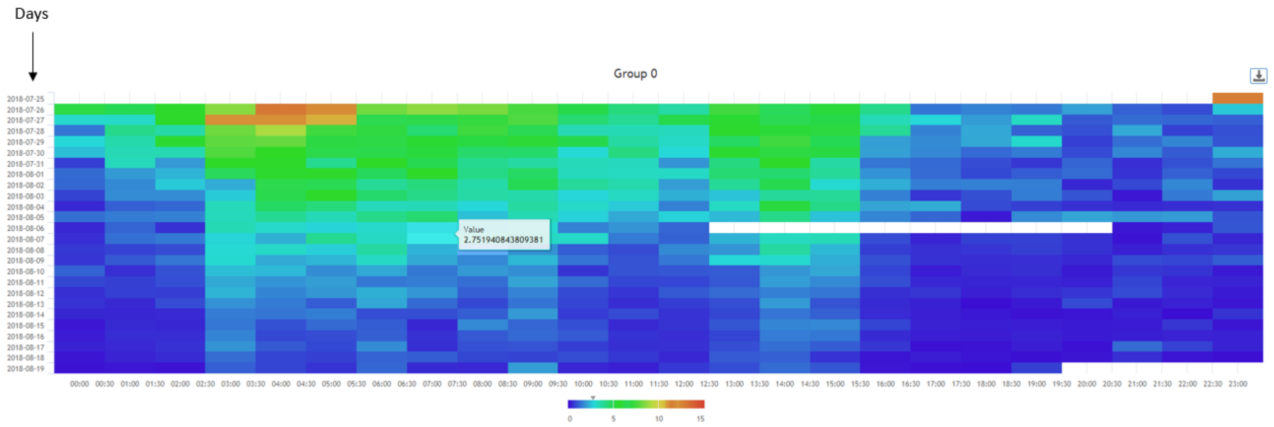

24 hours of the day, starting from midnight to midnight

##### 3.3.6 Box Plot

Spatial aggregation is the same of the Line Chart. Temporal aggregation is forced to be calculated on 24 hours (the day). Result is a single data point for each selected element (cage or animal). The Box Plot chart is available only when multiple elements (cages or animals) are selected.

The BOX PLOT feature enables 5 different points per data sample:

- Median
- 1st Quantile
- 3rd Quantile
- Min
- Max

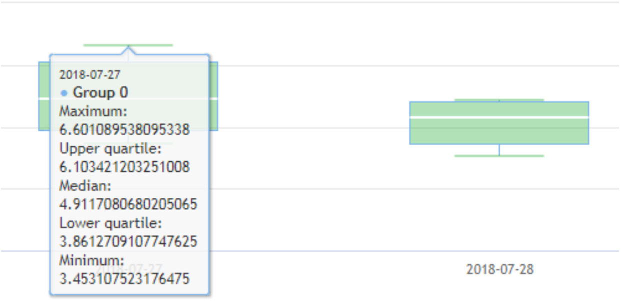
